# Pupil dilation reflects effortful action invigoration in overcoming aversive Pavlovian biases

**DOI:** 10.1101/2023.12.28.573353

**Authors:** Johannes Algermissen, Hanneke E. M. den Ouden

## Abstract

“Pavlovian” or “motivational” biases describe the phenomenon that the valence of prospective outcomes modulates action invigoration: Reward prospect invigorates action, while punishment prospect suppresses it. The adaptive role of these biases in decision-making is still unclear. One idea is that they constitute a fast-and-frugal decision strategy in situations characterized by high arousal, e.g., in presence of a predator, which demand a quick response. In this pre-registered study (*N* = 35), we tested whether such a situation—induced via subliminally presented angry vs. neutral faces—leads to increased reliance on Pavlovian biases. We measured trial-by-trial arousal by tracking pupil diameter while participants performed an orthogonalized Motivational Go/NoGo Task. Pavlovian biases were present in responses, reaction times, and even gaze, with lower gaze dispersion under aversive cues reflecting “freezing of gaze”. The subliminally presented faces did not affect responses, reaction times, or pupil diameter, suggesting that the arousal manipulation was ineffective. However, pupil dilations reflected facets of bias suppression, specifically the physical (but not cognitive) effort needed to overcome aversive inhibition: Particularly strong and sustained dilations occurred when participants managed to perform Go responses to aversive cues. Conversely, no such dilations occurred when they managed to inhibit responses to Win cues. These results suggest that pupil diameter does not reflect response conflict per se nor the inhibition of prepotent responses, but specifically effortful action invigoration as needed to overcome aversive inhibition. We discuss our results in the context of the “value of work” theory of striatal dopamine.

---

Humans and other animals are assumed to have different, parallel decision-making systems at their disposal that solve decision problems in different ways (Kahneman, 2011; Loewenstein & O’Donoghue, 2004; Metcalfe & Mischel, 1999; Milli, Lieder, & Griffiths, 2021; Shiffrin & Schneider, 1977). Some of these systems prioritize speed on behalf of accuracy, yielding quick, but seemingly inaccurate or “irrational” decisions. Other systems prioritize accuracy and yield more “rational” decisions, but at the cost of lower speed and increased mental resource demand (Dayan, 2014). One particularly simple, but quick system might be the so-called “Pavlovian” system, responsible for “Pavlovian” or “motivational” biases in behavior (Dayan, Niv, Seymour, & Daw, 2006; Guitart-Masip, Duzel, Dolan, & Dayan, 2014). This system allows the value of cues in the environment—associated with rewards (positive value) or punishments (negative value)—to influence action selection: in the presence of stimuli which signal that a reward can be gained (appetitive cues or “Win cues”), it invigorates behavior and drives more frequent and faster responses. In contrast, in the presence of stimuli which signal that a loss needs to be stimuli (aversive cues or “Avoid” cues), it suppresses behavior and leads to fewer and slower responses. Given that these biases seem to be altered in depression (Huys et al., 2016; Nord, Lawson, Huys, Pilling, & Roiser, 2018), traumas (Ousdal et al., 2018), anxiety disorders (Mkrtchian, Aylward, Dayan, Roiser, & Robinson, 2017), and alcohol addiction (Chen et al., 2023; Schad et al., 2020), understanding their role in everyday life might shed light on the etiology and maintenance of these disorders.

The presence of multiple decision systems necessitates an arbitration of which system to rely on in a particular situation, potentially driven by which class of situations or ecological niche each system is most “adaptive” in. Previous frameworks have suggested that different decision systems are selected based on their performance in achieving an optimal tradeoff between speed and accuracy (Daw, Niv, & Dayan, 2005; Keramati, Dezfouli, & Piray, 2011; Milli et al., 2021). Under this framework, Pavlovian biases have been suggested to constitute “default response options” in unfamiliar and/ or seemingly uncontrollable environments in which the recruitment of more effortful, “instrumental” control systems does not increase the rate of returned rewards (Dorfman & Gershman, 2019), In such situations, Pavlovian biases might constitute sensible “priors” about which action-outcome contingencies might hold in an environment (Moutoussis et al., 2018). Other frameworks have characterized Pavlovian control as an “emergency break” that takes over behavior in presence of particularly large rewards or threats, e.g., when facing a dangerous predator (O’Doherty, Cockburn, & Pauli, 2017). Under such circumstances, the Pavlovian system might trump other systems and induce a global inhibition of all motor effectors, characteristic of the freezing response (Roelofs, 2017; Roelofs & Dayan, 2022; Rösler & Gamer, 2019) and commonly induced by unexpected and surprising events (Schmidt & Berke, 2017; Wessel, 2018; Wessel & Aron, 2017). Notably, freezing seems to occur automatically and outside voluntary control, corroborating its likely “Pavlovian” nature. However, so far, there has been little causal evidence for such “emergency”, high-arousal situations exacerbating Pavlovian biases.

In this study, we aimed to test the causal effect of arousal on the size of Pavlovian biases. While many ways of inducing arousal in experimental lab settings exist, most of them can lead to deliberate shifts in response strategy and potentially induce demand characteristics. A promising way circumventing such strategy shifts is the subliminal presentation of arousing stimuli that cannot be consciously perceived by participants. For example, a study that used subliminally presented disgusted faces found these cues to exacerbate biases in a perceptual decision-making task (Allen et al., 2016). As a validation of their subliminal arousal manipulation, they found disgusted faces to also affect pupil diameter and heart rate— physiological indices of autonomic arousal. Using emotional faces might be a promising way of modulating Pavlovian biases as suggested that found supraliminally presented angry faces to promote freezing (Ly, Huys, Stins, Roelofs, & Cools, 2014). Motivated by these findings, we used subliminally presented angry vs. neutral face stimuli to manipulate arousal and test its effect on the size of Pavlovian biases in behavior. As a manipulation check, we measured pupil diameter, which is commonly interpreted as an index of arousal (Strauch, Wang, Einhäuser, Van der Stigchel, & Naber, 2022) and has in the past been used as a manipulation check in studies using subliminally presented faces (Allen et al., 2016).

To test whether arousal induced by a subliminal presentation of angry/ neutral faces amplified Pavlovian biases, we combined the orthogonalized Motivational Go/NoGo Task, a task measuring Pavlovian biases in humans, with a subliminal arousal induction while measuring participants’ pupil diameter. We expected Win/ Avoid cues featured in the Motivational Go/NoGo Task to induce Pavlovian biases. Furthermore, we hypothesized that subliminally presented angry (compared to neutral) faces would induce heightened arousal, reflected in stronger pupil dilation, which should amplify these biases. We thus expected an interaction between cue valence and the arousal manipulation, with a stronger valence effect, i.e., more Go responses to Win than Avoid cues (reflecting Pavlovian biases), in states of high induced arousal (angry face prime).

Neither responses, reaction times, nor pupil dilation reflected our subliminal arousal manipulation. These null results suggest that the manipulation was ineffective, which meant we could not assess our preregistered hypotheses. We proceeded with exploratory analyses testing whether features of the task itself were reflected in pupil diameter. A feature that might induce heightened arousal, reflected in pupil diameter, is response conflict between the response required to obtain rewards/ avoid punishments and the response triggered by Pavlovian biases (Cavanagh, Eisenberg, Guitart-Masip, Huys, & Frank, 2013; Swart et al., 2018). Participants likely need to recruit additional cognitive and/ or physical effort to resolve this conflict (Frank, 2006; Shenhav et al., 2017). We tested whether pupil dilation reflects the *cognitive effort* associated with the suppressing Pavlovian biases, more globally, or the *physical effort* specifically required for invigorating Go responses, which is particularly warranted when overcoming aversive inhibition induced by Avoid cues.

Several recent studies have suggested that pupil diameter reflects cognitive effort associated with the suppression of prepotent responses, e.g., in cognitive conflict tasks such as the Stroop, Simon, or Flanker task, or in task switching paradigms (da Silva Castanheira, LoParco, & Otto, 2020; D’Ascenzo, Iani, Guidotti, Laeng, & Rubichi, 2016; Rondeel, Van Steenbergen, Holland, & van Knippenberg, 2015; van der Wel & van Steenbergen, 2018; van Steenbergen & Band, 2013). The very same conflict detection and resolution mechanisms might be required for inhibiting Pavlovian biases, i.e., when suppressing Go responses to Win cues or invigorating Go responses in presence of Avoid cues. Previous research has observed increases in midfrontal EEG theta power when participants successfully inhibited Pavlovian biases (Cavanagh et al., 2013; Swart et al., 2018), and other research has found theta power to be correlated with pupil diameter (Dippel, Mückschel, Ziemssen, & Beste, 2017; Lin, Saunders, Hutcherson, & Inzlicht, 2018). In line with these studies, one might expect higher pupil dilations when participants perform a bias-incongruent response, i.e., they successfully inhibit their Pavlovian biases. This pattern would be reflected in an interaction effect between cue valence and response, with stronger dilations for bias-incongruent responses, i.e., Go responses to Avoid cues and NoGo responses to Win cues (Fig. 1E).

**Figure 1.**
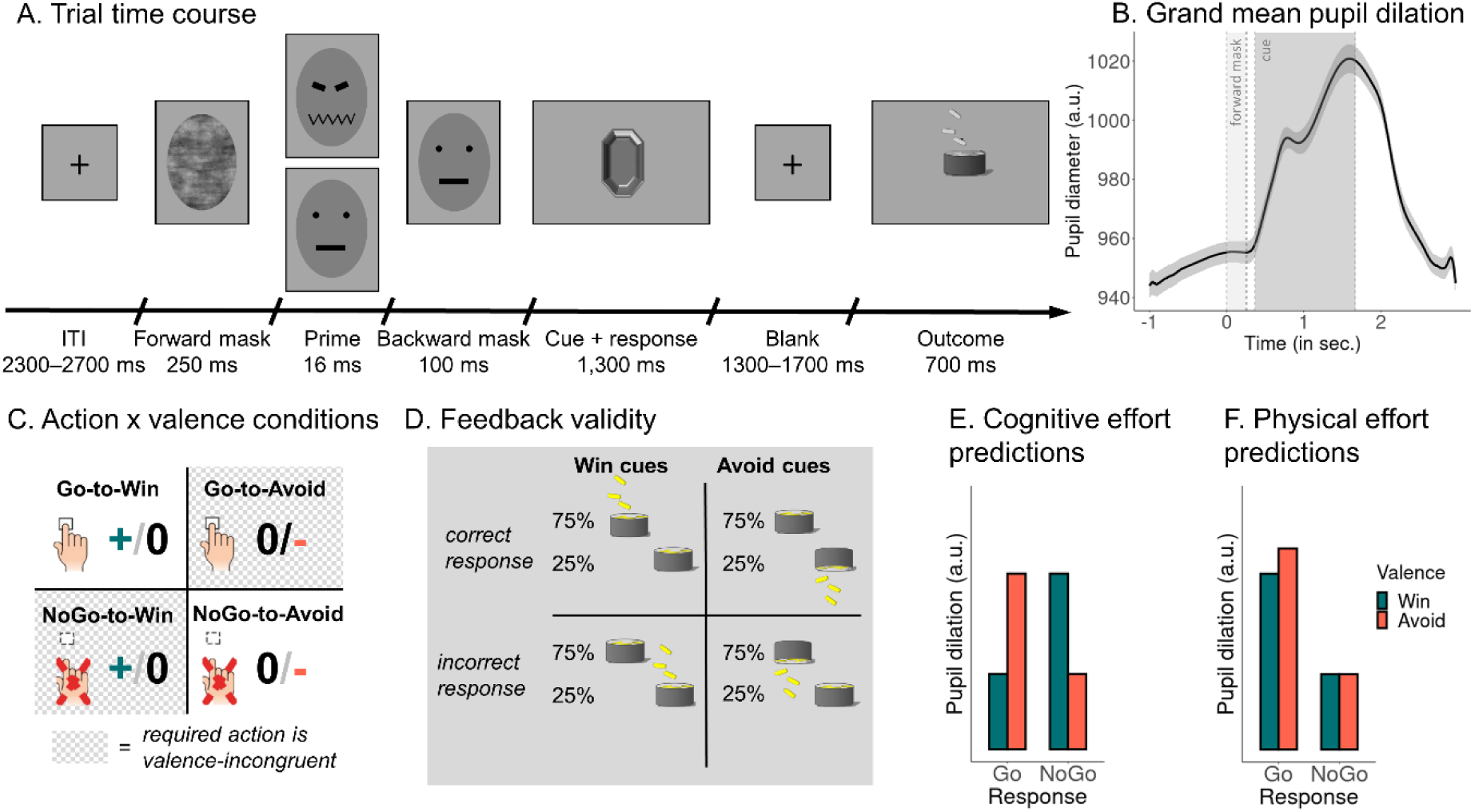
Task design. *Note.* **A**. *Trial time course*. Each trial starts with a forward mask presented for 250 ms (pixel-permuted version of the neutral prime), a prime stimulus (angry or neutral face) for 16 ms, and a backwards mask (another neutral face) for 100 ms. Participants then see one of four cues and have to decide whether to respond with a button press (“Go”) or not (“NoGo”). After a variable interval, the outcome (gain, neutral, loss of points) is shown. Face stimuli from the Karolinska Directed Emotional Faces set (Lundqvist et al., 1998). **B.** *Grand mean average of the pupil dilation for all trials of all participants*. Vertical dashed lines indicate the onset of the forward mask (at 0 ms), the prime (at 250 ms), the backwards mask (at 266 ms), the cue onset (at 366 ms), and the cue offset (at 1666 ms). **C.** *Task conditions*. Half of the cues are “Win” cues for which participants can gain points, while the other half are “Avoid” cues for which participants can lose points. Orthogonal to the cue valence, one half of the cues requires a Go response (“Go” cues) while the other half requires a NoGo response (“NoGo” cues). **D.** *Feedback given cue valence and accura*cy. For half of the cues (“Win cues”), participants receive mostly gains in points (money falling into a can) for correct responses, but no change in point score (a can) for incorrect responses. For the other half (“Avoid” cues), they receive no change in point score (a can) for correct responses, but a loss of points (money falling out of a can) for incorrect responses. **E.** Predictions of per-condition pupil dilation according to the hypothesis that pupil dilation reflects cognitive effort. Dilations are higher for bias-incongruent responses, i.e., Go responses to Avoid cues and NoGo responses to Win cues, irrespective of whether a Go or NoGo response has to be performed. **F.** Prediction of per-condition pupil dilation according to the hypothesis that pupil dilation reflects physical effort. Dilations are higher for Go than NoGo responses and particularly high for Go responses to Avoid cues, for which aversive inhibition must be overcome.

Besides cognitive effort, an alternative hypothesis could be that pupil diameter reflects physical effort associated with Go responses. At first glance, “physical effort” might sound like a hyperbole in the context of the Motivational Go/NoGo Task, which requires only single button presses. Still, we choose this term as the decision to act, at all, and the decision to recruit physical effort are likely driven by the same mechanisms (Berke, 2018; Hamid, 2021).

Furthermore, animal research has used Go/ NoGo tasks akin to our design to study physical effort (Hamid et al., 2016; Syed et al., 2016). Finally, also past research in humans has used relatively minor actions, e.g., the speed of single saccades (Manohar et al., 2015) or choice options that require several keyboard presses (Treadway, Buckholtz, Schwartzman, Lambert, & Zald, 2009), as indices of physical effort.

An association between pupil diameter and movement is long established as pupil diameter reliably increases during movement preparation (Richer & Beatty, 1985; Richer, Silverman, & Beatty, 1983; Schacht, Dimigen, & Sommer, 2010). In classic Go/ NoGo tasks, dilations are generally higher for Go responses than NoGo responses (Schacht et al., 2010; Van der Molen, Boomsma, Jennings, & Nieuwboer, 1989). Going beyond previous research, in the context of the Motivational Go/NoGo Task, it should be particular high for Go responses to Avoid cues (compared to Go responses to Win cues) since participants have to recruit additional physical effort to overcome aversive inhibition induced by Avoid cues. Thus, there should be an effect of cue valence on pupil dilation for Go responses. Conversely, there is no comparable physical effort requirement for inhibiting prepotent Go responses to Win cues, and thus one should expect no effect of cue valence on pupil dilation for NoGo responses. According to this hypothesis, one would expect a main effect of response on pupil dilation, with stronger dilations during Go than NoGo responses, qualified by an interaction between response and cue valence, with particularly strong dilations for Go responses to Avoid cues (Fig. 1F). As the crucial condition of comparison, NoGo responses to Win cues require high cognitive effort (for suppressing Pavlovian biases), but low physical effort (since no movement is prepared), allowing us to dissociate cognitive effort and physical effort accounts of pupil diameter.

## Methods

### Participants and Exclusion Criteria

Sample size (M_age_ = 22.37, SD_age_ = 2.68, range 18–30; 18 women, 17 men; 27 right-handed, 8 left-handed; 18 with right eye dominant; 17 with left eye dominant). The study design, hypotheses, and analysis plan was pre-registered on OSF under https://osf.io/ue397.

English-speaking participants in the age range of 18–35 years old were recruited via the SONA Radboud Research Participation System of Radboud University. Only participants with unimpaired vision or contact lenses were admitted. Exclusion criteria comprised previous neurological treatment, cerebral concussion, brain surgery, or epilepsy. Participants were excluded from all analyses for three (pre-registered) reasons: (a) guessing the hypothesis in the debriefing, (b) performance not significantly above chance (tested by using required action to predict performed action with a logistic regression; only participants with p < .05 were maintained); and (c) no pupil data on more than 128 trials (50% of trials). None of these criteria applied to any of the participants. Hence, the final sample size for all analyses comprised *N* = 35. This reported research was approved by the local ethics committee of the Faculty of Social Sciences at Radboud University (proposal no. ECSW-2018-171 and ECSW-2019-055) in accordance with the Declaration of Helsinki.

The sample size was not based on a power analysis, but on lab availability for this project (four weeks, April 16 till May 17, 2019) as this study was conducted as around several thesis projects. The sample size of *N* = 35 was comparable to previous studies investigating Pavlovian biases with the same task (Algermissen, Swart, Scheeringa, Cools, & den Ouden, 2022; Swart et al., 2018) and slightly larger than the study which inspired the subliminal arousal priming manipulation (Allen et al., 2016). A post-hoc sensitivity power analysis yielded that, given 35 participants providing 256 trials (thus 8,960 trials in total), and assuming intra-cluster coefficients of 0.04 for responses, 0.14 for RTs, and 0.17 for dilations (all estimated from the data), the effective sample size was *n* = 4,090 for responses, *n* = 1,558 for RTs, and *n* = 1,329 for dilations, respectively, which allows to detect effects of β > 0.04 for responses, β > 0.07 for RTs, and β > 0.08 for dilations (standardized regression coefficients) with 80% power (Aarts, Verhage, Veenvliet, Dolan, & van der Sluis, 2014).

### Procedure

Participants completed a single experimental session that lasted about 45 minutes. They provided informed consent, underwent an 9-point eye-tracker calibration, read computerized instructions and performed four practice trials for each of the four cue conditions. Afterwards, they completed 256 trials of the Motivational Go/NoGo Task. After the task, participants completed measures of trait anxiety (STAI, Form Y-2, 20 items) (Spielberger, Gorssuch, Lushene, Vagg, & Jacobs, 1983) and impulsivity (UPPS-P short version, five sub scales, 20 items) (Cyders, Littlefield, Coffey, & Karyadi, 2014), which were part of final year theses written on this data set. At the end, participants went through a funnel debriefing asking them what they thought the hypothesis investigated in the experiment was, if they used any strategies not contained in the task instructions (and, if yes, describe them), whether they noticed anything special about the task not mentioned in the instructions (and, if yes, describe it), if they noticed anything special about the face at the beginning of each trial (and, if yes, describe it), whether they recognized the emotions of the face presented very briefly (and, if yes, describe them), and finally, given that there was an angry and a neutral face presented, what they thought the hypothesis investigated in the experiment was. After the completion of the experiment, participants received course credit in compensation plus a performance-dependent candy bar for task accuracy > 75%.

### Apparatus

Reporting follows recently suggested guidelines for eye-tracking studies (Fiedler, Schulte-Mecklenbeck, Renkewitz, & Orquin, 2020). The experiment was performed in a dimly lit, sound-attenuated room, with participants’ head stabilized with a chin rest. The experimental task was coded in PsychoPy 1.90.3 on Python 2.7, presented on a BenQ XL2420Z screen (1920 x 1080 pixels resolution, refresh rate 144 Hz). People’s dominant eye was recorded with an EyeLink 1000 tracker (SR Research, Mississauga, Ontario, Canada; sampling rate of 1,000 Hz; spatial resolution of 0.01° of visual angle, monocular recording). The chinrest was placed about 90 cm in front of the screen and 70 cm in front of the eye-tracker. Before the task, participants underwent the standard 9-point calibration and validation procedure provided by SR Research, which was repeated until error for all nine points was below 1°. The screen background during the task was of the same gray (RGB [166, 166, 166]) as during the calibration. Participants were instructed to focus on the fixation cross/ center of the screen throughout the task. Manual responses (Go) were performed via the space bar of the keyboard.

### Task

Participants performed 256 trials (split in four blocks of 64 trials each) of an orthogonalized Motivational Go/ NoGo learning task (Swart et al., 2018). Unlike classic Go/ NoGo tasks in which responses are instructed, in this task, required responses for different cues had to be learned from probabilistic feedback. An equal number of Go and NoGo responses was required. Hence, Go responses were not as prepotent as in classic Go/ NoGo tasks. Compared to previous studies using this task, the trial time line of the task was slowed down to reliably measure pupil fluctuations. Each trial started with a series of rapidly presented images used to subliminally induce arousal, followed by a cue indicating the required response and potential outcome of the trial, and finished with the outcome.

The arousal priming manipulation closely followed a procedure previously found effective (Allen et al., 2016). It consisted of a “prime” image presented for 16 ms (two frames), which was either an angry face (image ID AM29ANS; high arousal) or a neutral face (ID AM29NES; low arousal) from the Karolinska Directed Emotional Faces data set (Lundqvist, Flykt, & Öhman, 1998). Hair and background were removed from the face stimulus by cropping it to an elliptical shape (size 281 x 381 pixels; 5.0° x 6.7° visual angle; Fig. 1A). To prevent conscious recognition of the prime stimulus, it was flanked by a forward mask, which was a version of the neutral prime with pixels randomly permuted, presented for 250 ms before the prime, and a backward mask, which was another neutral face taken from the same face data set (ID AM10NES), presented for 100 ms after the prime (Allen et al., 2016). Due to the “subliminal” presentation of the face primes, participants were supposedly unaware of these primes and could not respond to them. Participants were instructed that the presentation of the backward mask served to keep their attention focused on the task.

Next, participants saw one of four cues for 1,300 ms. Unlike the face primes, cues were supraliminally presented gem-shaped stimuli (Fig. 1A) that either required a Go or a NoGo response and either offered the chance to win points (Win cues) or to avoid losing points (Avoid cues). The task was a fully orthogonalized 2 x 2 x 2 design with the factors arousal manipulation via face primes (angry/ neutral face), cue valence (Win/ Avoid), and required action (Go/ NoGo). Participants had to learn from experience whether a cue offered the chance to win points for correct responses (and no change in points for incorrect responses; “Win” cues) or the chance to lose points for incorrect responses (and no change in points for correct responses; “Avoid” cues; Fig. 1C). Also, they needed to learn from trial-and-error whether the cue required a Go response (space bar press) or NoGo response (no press). Cues were of size 300 x 300 pixels (5.3° x 5.3°), presented centrally, set to grayscale and matched for average luminance and local statistical properties using the SHINE toolbox (Willenbockel et al., 2010). Cue assignment to task conditions was counterbalanced across participants. Each cue was presented 16 times in total (eight times with the high arousal and eight times with the low arousal prime), with cue presentation interleaved in a pseudo-randomized way (not more than one consecutive cue repetition). Each of the four blocks featured a new set of four cues to prevent ceiling effects in performance and to maximize the time during which participants were (at least partially) unsure about the correct response.

After a variable inter-stimulus interval (uniform distribution between 1,300–1,700 ms in steps of 100 ms), the outcome was presented for 700 ms. Outcomes consisted in either money falling into a can (positive feedback for Win cues), money falling out of a can (negative feedback for Avoid cues), or simply a can (negative feedback for Win cues/ positive feedback for Avoid cues). Feedback validity was 75%, i.e., correct responses were followed by positive feedback and incorrect responses followed by negative feedback on 75% of trials, with the reverse being the case on the remaining 25% of trials (Fig. 1C). Trials finished with a variable inter-trial interval (uniform distribution between 2,300–2,700ms in steps of 100 ms).

### Data Preprocessing

#### Behavior

For analyses using RTs, we excluded trials with RTs < 300 ms (in total 36 trials out of 8,960 trials; per participant: *M* = 1.01, *SD* = 3.06, range 0–14) since it is implausible that these very fast responses incorporated knowledge about the cue. Note that this step was not pre-registered, but the same procedure was used in previous studies in which we used the same task (Algermissen et al., 2022; Swart et al., 2017). Analyses including all RTs lead to identical conclusions.

#### Pupil preprocessing

Pupil data were preprocessed in R following previously published pipelines (de Gee et al., 2017; Urai, Braun, & Donner, 2017). First, pupil data was epoched into trials from 1,000 ms before until 2,966 ms after forward mask onset (i.e., until the earliest possible end of the ISI/ before possible outcome onset). Note that the pre-registration specifies a different time range (1,000 ms before until 1,666 ms after forward mask onset; i.e. exactly until task cue offset) under the assumption of a peak of the pupil response around 1,000 ms (Hoeks & Levelt, 1993). However, in fact, the grand average pupil response in this data peaked at 1,584 ms (Fig. 1B), i.e., close to the end of the pre-registered time window, with per-trial dilations peaking outside the pre-registered window on almost half of the trials (assuming symmetric noise on the peak latency). The grand average pupil time course only returned to baseline levels around 3,000 ms after forward mask onset (Fig. 1B). We thus decided to extend the time window until 2,966 ms, i.e., until the earliest possible onset of an outcome (Fig. 1A). After epoching, the timing of blinks and saccades (as automatically detected by the EyeLink software) was extracted. These gaps of missing data were zero-padded by deleting 150 ms (for blinks, 20 ms for saccades) of samples before and after them (as recommended by the EyeLink manufacturer). In addition, we computed the first derivative of the pupil time course and marked abnormally fast pupil changes (absolute values of the z-standardized first derivative higher than 2). If two such marks occurred less than 10 samples away from each other, we deleted all samples in-between. Finally, we interpolated missing or deleted samples with linear interpolation and low-pass filtered the data at 6 Hz with a 3-order Butterworth filter. We deleted the first and last 250 ms of each trial to remove edge artifacts caused by the filter. We converted pupil diameter to units of modulation (percent signal change) around the mean of the pupil time series of each block using the grand-mean pupil diameter per block (i.e., 64 trials forming one block). Trials with more than 50% of missing/ interpolated data were excluded (in total 166 trials out of 8,960 trials; per participant: *M* = 4.74, *SD* = 9.10, range 0–43). Finally, we computed the trial-by-trial pupil baseline as mean pupil diameter in the 500 ms before the onset of the forward mask and the maximal pupil dilation as the maximal value during the 2,966 ms after onset of the forward mask (i.e. until the earliest possible end of the ISI). We then computed the trial-by-trial pupil dilation by subtracting the trial-by-trial pupil baseline from the trial-by-trial maximal dilation.

#### Gaze preprocessing

We analyzed the gaze data similar to the pupil data. After epoching, the timing of blinks and saccades (as automatically detected by the Eyelink software) was extracted. These gaps of missing data were zero-padded by deleting 150 ms (for blinks, 20 ms for saccades) of samples before and after them (as recommended by the Eyelink manufacturer). In addition, we computed the first derivative of the pupil time course and marked abnormally fast pupil changes (absolute values of the z-standardized first derivative higher than 2). If two such marks occurred less than 10 samples away from each other, we deleted all samples in-between. We did not apply interpolation for missing gaze data.

### Data Analysis

#### Mixed-effects regression models

For regression analyses, we used mixed-effects linear regression (function lmer) and logistic regression (function glmer) as implemented in the package lme4 in R (Bates, Mächler, Bolker, & Walker, 2015). We used generalized linear models with a binomial link function (i.e., logistic regression) for binary dependent variables (Go vs. NoGo responses) and linear models for continuous variables such as RTs, pupil baseline, and pupil dilation. We used zero-sum coding for categorical independent variables. All continuous dependent and independent variables were standardized such that regression weights can be interpreted as standardized regression coefficients. All regression models contained a fixed intercept. We added all possible random intercepts, slopes, and correlations to achieve a maximal random effects structure (Barr, Levy, Scheepers, & Tily, 2013). *P*-values were computed using likelihood ratio tests with the package afex (Singmann, Bolker, Westfall, & Aust, 2018). We considered *p*-values smaller than α = 0.05 as statistically significant.

As an initial manipulation check, we fit a mixed-effects logistic regression model with responses (Go/NoGo) as dependent variable and required action (Go/ NoGo), cue valence (Win/ Avoid), and their interaction as independent variables. Furthermore, we fit an equivalent linear regression model with RTs as dependent variable. For the confirmatory models involving the arousal priming manipulation as independent variable, see Supplementary Material S05. For the confirmatory models involving trial-by-trial pupil dilation as independent variable, see Supplementary Material S06. In further exploratory analyses, we fit a mixed-effects linear regression model with trial-by-trial pupil dilation as dependent variable and response, cue valence, and their interaction as independent variables.

#### Cluster-based permutation tests on pupil data

In order to test whether the millisecond-by-millisecond pupil time course during a trial differed between conditions, we used cluster-based permutation tests (Maris & Oostenveld, 2007). For cue-locked time courses, we epoched the pre-processed data into trials from -1,000 ms before until 2,966 ms after mask onset, sorted trials into task conditions, and computed the average time course per condition per participant. For response-locked time courses, on trials with Go responses, we epoched the data relative to the trial-specific RT. On trials with NoGo responses, we epoched the data relative to pseudo-RTs computed as the mean RT to Win cues/ Avoid cues for a given participant.

We then computed a permutation null distribution by, for 10,000 iterations, randomly exchanging the labels of conditions, computing the mean difference between conditions per participant, computing the overall mean difference between conditions across participants, thresholding this difference at |*t*| > 2, computing the sum of *t*-values for each cluster of adjacent samples above threshold (cluster mass), and retaining the largest cluster mass detected for each iteration. We then compared the empirical cluster mass obtained from the actual data to the permutation null distribution and computed the permutation *p*-value as the number of iterations with a larger cluster mass than the empirical cluster mass divided by the total number of iterations. To correct for pre-trial baseline differences, for each condition, we subtracted the value at time point 0 (also for each iteration in the permutation distribution).

#### Cluster-based permutation tests on gaze data

In line with previous studies reporting freezing of gaze (Rösler & Gamer, 2019), we used the mean gaze position (x-and y-coordinates) in the 500 ms before mask onset (while the fixation cross from the inter-trial interval was on the screen) as a trial-by-trial baseline and compute the absolute deviation (Euclidean distance in pixels) from that baseline for each trial (“gaze dispersion”). This procedure corrects for any drift in the eye-tracking calibration over time. We then computed the mean distance from the pre-trial baseline at any time point during cue presentation separately for Win and Avoid cues for every participant. We performed a cluster-based permutation test with 10,000 iterations and a cluster-forming threshold of |t| > 2 to test for any difference in the distance from the center between Win and Avoid cues.

#### Generalized additive mixed-effects models

Additive models use smooth functions of a set of predictors (i.e., thin plate regression splines) in order to model a time series. They allow for testing whether the modeled time series differs between conditions. The shape of a smooth function is fitted to the data and can be linear or non-linear, allowing more flexibility in capturing non-linear trends over time compared to linear models, which makes them particularly suited for analyzing pupillometry data (Algermissen, Bijleveld, Jostmann, & Holland, 2019; Baayen, Vasishth, Kliegl, & Bates, 2017; van Rij, Hendriks, van Rijn, Baayen, & Wood, 2019). A smooth function regularizes the raw time courses and in this way suppresses high-frequency (trial-by-trial) noise. It also accounts for non-zero auto-correlation between residuals, which is assumed to be zero in linear models.

In order to test whether the effect of task conditions changed over time, we fit generalized additive mixed-effects models with the trial-by-trial pupil dilation as dependent variable and separate effects of cue repetition (1–16) for each response condition (Go-to-Win, Go-to-Avoid, NoGo-to-Win, NoGo-to-Avoid) as independent variables. We modeled the time course of cue repetition as a factor smooth (which has a similar, but potentially non-linear effect as adding a random intercept and a random slope) for each participant for each block, allowing for the possibility that condition differences were different in different participants in different blocks (equivalent to a full random-effects structure). We used a scaled *t*-distribution instead of a Gaussian distribution for the response variable in case it led to lower fREML values (which was the case for RTs, pupil baselines, and dilations). In case of significant residual auto-correlation at lag 1 (which was the case for baselines), we added an AR(1) auto-regressive model, with the proportionality constant set to the lag 1-correlation of the residuals from the model without the AR(1). For all fitted models, we visually checked that residuals were approximately normally distributed using quantile-quantile plots and whether auto-correlation was near zero using auto-correlation plots (van Rij et al., 2019).

### Transparency and openness

We report how we determined our sample size, all data exclusions, all manipulations, and all measures in the study. All data, analysis code, and research materials will be shared upon publication. The study design, hypotheses, and confirmatory analysis plan were pre-registered under https://osf.io/57zjh and updated under https://osf.io/azqjt (extending data collection by one week).

We deviated from the pre-registration in the definition of the time window in which pupil dilation was defined. This time window was pre-registered as spanning 1,000 ms before until 1,666 ms after forward mask onset (i.e., exactly until task cue offset). Given the unexpectedly late peak in the grand average pupil time course, we extended this time window until 2,966 ms after mask onset (i.e., until the earliest possible onset of an outcome). The pre-registration also comprised plans for computational models and a deconvolution GLM approach. Since the arousal priming manipulation did not work, these plans were eventually not carried. Finally, we performed exploratory analyses of pupil dilation as a function of task factors (testing physical effort and cognitive effort accounts of pupil diameter), which were not pre-registered.

Data were analyzed using R, version 4.1.3 (R Core Team, 2022). Models were fitted with the package lme4, version 1.1.31 (Bates et al., 2015). Plots were generated with ggplot, version 3.4.2 (Wickham, 2016).

## Results

### Manipulation checks: Learning and Pavlovian bias

First, in line with the pre-registration, we performed manipulation checks to replicate effects typically found with this task (Algermissen et al., 2022; Swart et al., 2018). We fit a mixed-effects logistic regression with responses (Go/ NoGo) as dependent variable and required action (Go/ NoGo) and cue valence (Win/ Avoid) as well as their interaction as independent variables (see Supplementary Material S01 for an overview of all regression results; see Supplementary Material S02 for means and standard deviations per condition). Participants performed significantly more (correct) Go responses to Go cues than (incorrect) Go responses to NoGo cues (required action), *b* = 1.367, 95%-CI [1.178, 1.556], χ^2^(1) = 66.523, *p* < .001, indicating that participants successfully learned the task (Fig. 2A-C). Overall, participants showed 82% accuracy for Go cues (SD = 10%, range 63–95%) and 69% accuracy for NoGo cues (SD = 15%, range 37–96%; see also Supplementary Material S02 for per condition mean response rates). Also, participants performed more Go responses to Win than to Avoid cues (cue valence), *b* = 0.538, 95%-CI [0.341, 0.734], χ^2^(1) = 20.986, *p* < .001, reflecting the Pavlovian bias (Fig. 2A-C). The interaction between required action and valence was not significant, *b* = 0.068, 95%-CI [-0.044, 0.181], χ^2^(1) = 1.348, *p* = .246, providing no evidence that the Pavlovian bias was stronger for Go or for NoGo cues.

**Figure 2.**
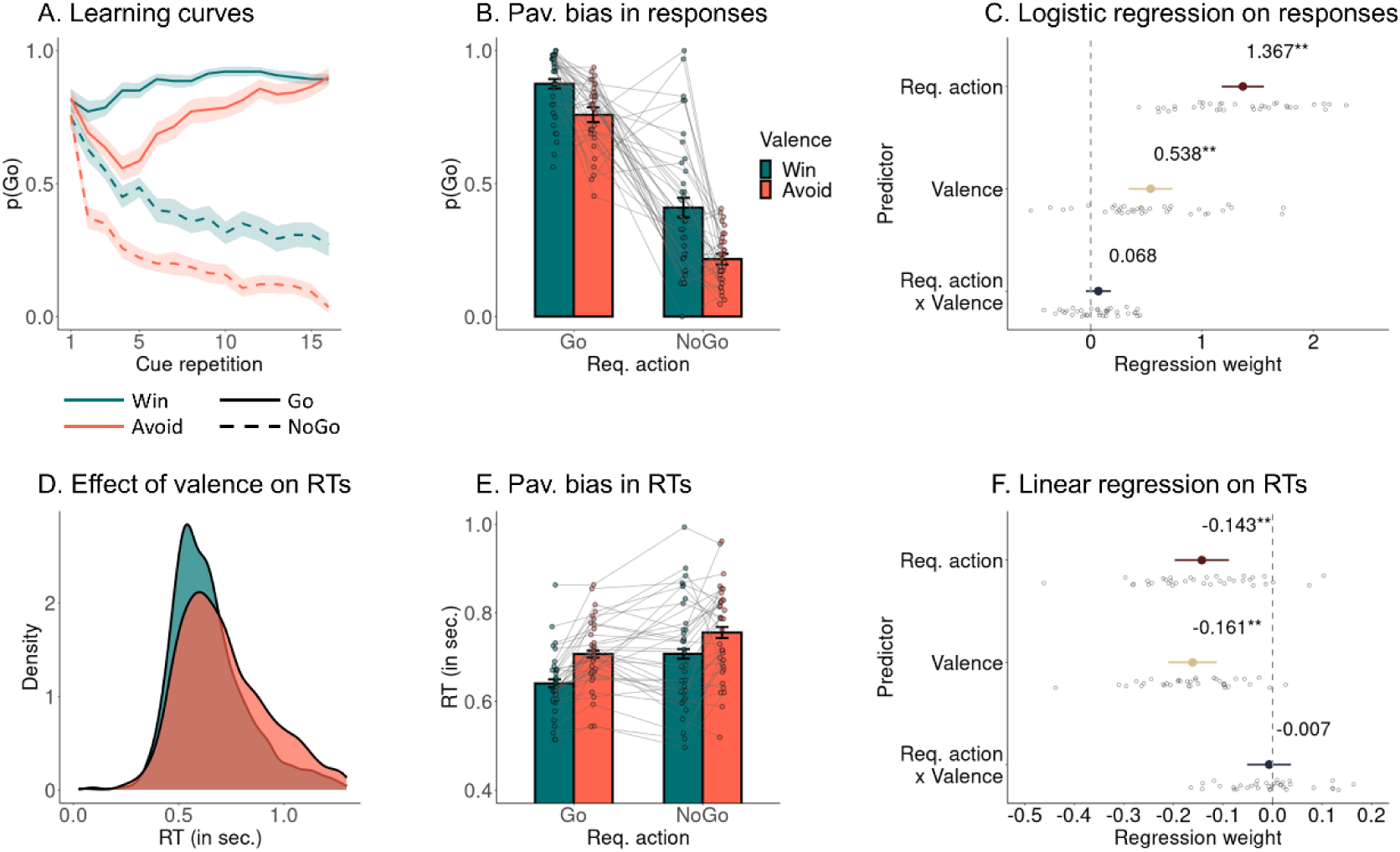
Effect of required action and cue valence on responses and RTs. *Note*. **A.** Trial-by-trial proportion of Go responses per cue condition. Participants learn to perform a Go response or not, with significantly more Go responses to Go cues than NoGo cues. Also, they perform significantly more Go responses to Win cues than to Avoid cues, reflecting the Pavlovian bias. Note that participants are initially unaware of the cue valence and have to infer it from (non-neutral) feedback, which explains why the bias only emerges after the first few trials. For the Go-to-Avoid conditions, the bias initially suppresses responding, and participants have to subsequently learn to overcome the bias and perform a Go response. This is reflected in the dip in Go responses for Go-to-Avoid cues for trials 1–5 when the negative valence of this cue is learned, and a subsequent rise in Go responding as the correct response to this cue is learned. Error bands are ± SEM across participants. **B.** Proportion of Go responses per cue condition (whiskers are ± SEM across participants, dots indicate individual participants). Participants show significantly more Go responses to Go than NoGo cues (reflecting learning) and significantly more Go responses to Win cues than Avoid cues (indicative of Pavlovian biases). **C.** Group-level (colored dot, 95%-CI) and individual-participant (grey dots) regression coefficients from a mixed-effects logistic regression of responses on required action, cue valence, and their interaction. **D**. Distribution of raw RTs separately per cue valence. **E.** Mean RTs per cue condition. Participants show significantly faster (correct) Go responses to Go than (incorrect) Go responses to NoGo cues and significantly faster responses to Win cues than Avoid cues (indicative of Pavlovian biases). **F.** Group-level and individual-participant regression coefficients from a mixed-effects linear regression of RTs on required action, cue valence, and their interaction.

Furthermore, we fit a mixed-effects linear regression with RTs as dependent variable and again required action, cue valence, and their interaction as independent variables. This analysis was omitted in the pre-registration, but in line with previous studies (Algermissen et al., 2022). RTs were only available for Go responses. Participants were faster at correct responses (to Go cues) than incorrect responses (to NoGo cues; required action), *b* = -0.143, 95%-CI [-0.197, -0.088], χ^2^(1) = 20.446, *p* < .001 (Fig. 2D-F). Also, they were faster at performing responses to Win than to Avoid cues (cue valence), *b* = -0.143, 95%-CI [-0.197, - 0.088], χ^2^(1) = 27.329, *p* < .001, again reflecting the Pavlovian biases (Fig. 2D-F). For the time courses of both effects over trials within a block, see Supplementary Material S03. The interaction between required action and valence was not significant, *b* = -0.007, 95%-CI [-0.051, 0.037], χ^2^(1) = 0.083, *p* = .773, providing no evidence that the Pavlovian bias was stronger for correct responses (to Go cues) or incorrect responses (to NoGo cues). Pavlovian biases in responses and RTs were not correlated with participants’ trait anxiety or impulsivity scores, providing no evidence for stronger or weaker biases in more impulsive/ more anxious individuals (see Supplementary Material S04). Taken together, these results corroborate that participants learned the task and exhibited Pavlovian biases.

### Exploratory analyses: Freezing of gaze induced by Avoid cues

Previous research on humans and animals has investigated the phenomenon of “freezing”, i.e., temporarily reduced body motion in presence of a thread (Blanchard, 2017; Roelofs, 2017). Freezing in humans is typically measured via reductions in heart rate (Hashemi et al., 2019; Klaassen et al., 2021) and bodily mobility (Ly et al., 2014) tracked with a stabilometric force-platform that records spontaneous fluctuations in body sway. Recently, it has been suggested that freezing might also affect gaze, with a stronger center bias and less visual exploration while participants prepare a response to avoid an electric shock (Merscher & Gamer, 2024; Merscher, Tovote, Pauli, & Gamer, 2022; Rösler & Gamer, 2019). Here, we tested whether a similar freezing of gaze pattern occurred during the presentation of Avoid compared to Win cues in the context of the Motivational Go/NoGo Task, testing for a difference in the absolute distance from the center of the screen (“gaze dispersion”) between trials with Win and Avoid cues.

A cluster-based permutation test in the time range of 0–500 ms after cue onset was significant (*p* = .024, two-sided; driven by a cluster above threshold 202–278 ms after cue onset; Fig. 3A, B). Distance from the center was lower on trials with Avoid cues than on trials with Win cues, in line with the idea of “freezing of gaze” induced by Avoid cues. Computing the maximal distance from the center in this time window for every trial, averaging distances for Win and Avoid cues per participant, and then averaging across participants confirmed this difference (Fig. 3C). Importantly, there was no difference in gaze dispersion between Win and Avoid cues on the first five repetitions of a cue, i.e., when participants were not fully aware of cue valence yet (Fig. 3D; no cluster above threshold), but this difference only emerged on cue repetitions 6–10 (Fig. 3E; *p* = .009, cluster above threshold from 242–281 ms after cue onset) and became stronger for cue repetitions 11–15 (Fig. 3F; multiple disconnected clusters above threshold between 55–353 ms after cue onset; largest cluster above threshold from 245–261 ms, *p* = .023).

**Figure 3.**
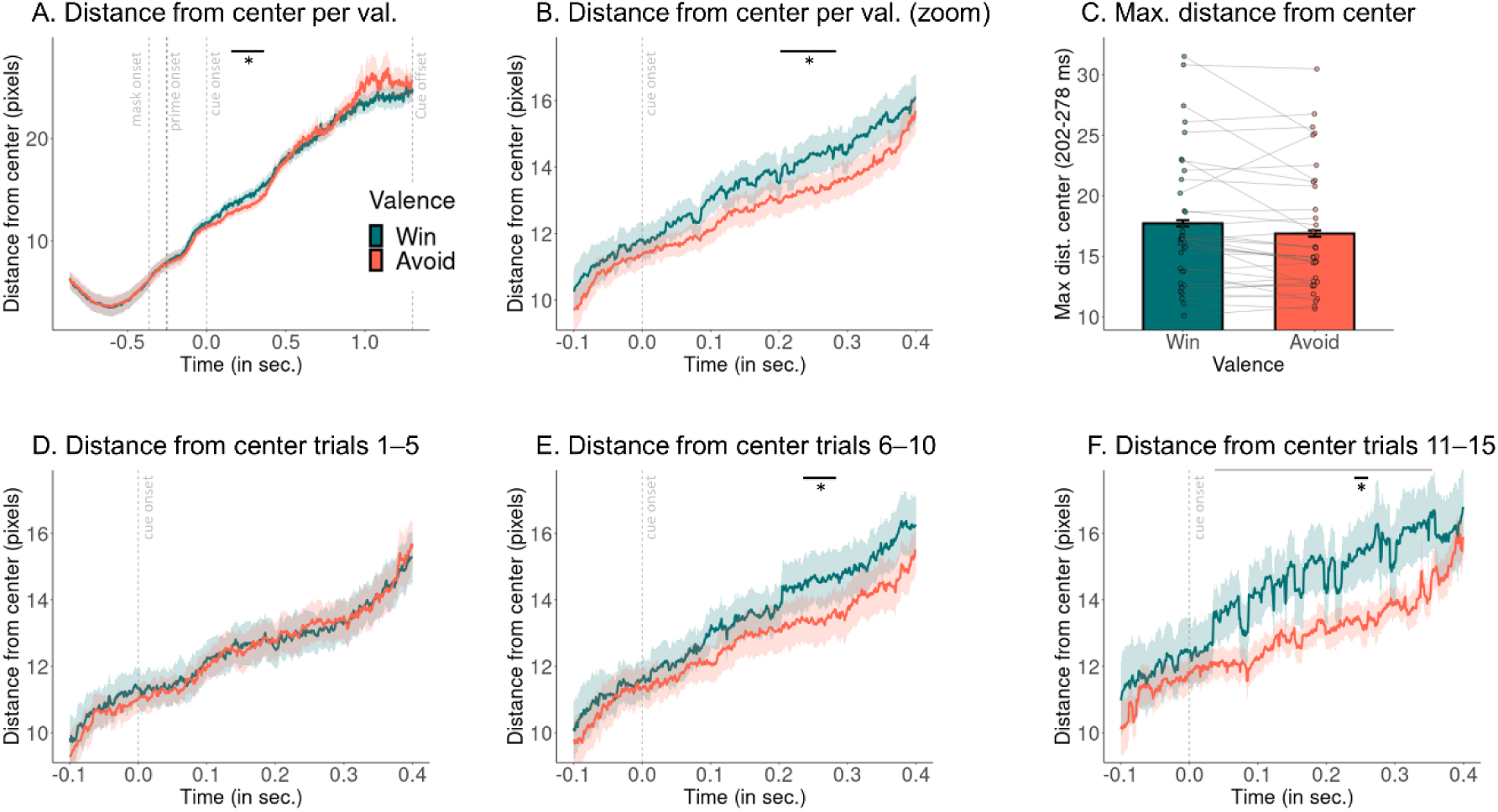
Freezing of gaze induced by Avoid cues. *Note.* **A.** Mean distance from the gaze position during the trial-by-trial baseline (“center”). Vertical dashed lines indicate the onset of the forward mask (at -366 ms), the prime (at -266 ms), the backwards mask (at -250 ms), the cue onset (at 0 ms), and the cue offset (at 1300 ms). Distance increases with time. Around 202–278 ms after cue onset, distance from the center is lower on trials with Avoid cues compared to trials with Win cues. **B.** Same as panel A, but zoomed into the time range of -100–400 ms after cue onset. **C.** Maximum distance from the pre-trial baseline (whiskers are ± SEM across participants, dots indicate individual participants) averaged for Win and Avoid cues for each participant. Distance is lower on trials with Avoid cues compared to trials with Win cues. **D-F.** Same as panel B, but computed for subsets of trials. While freezing of gaze is absent on the first five cue repetitions when participants are not yet fully aware of the cue valence (see learning curves in Fig. 2A), with no cluster above threshold (**D**), the freezing of gaze bias emerges on cue repetitions 6–10 (**E**; p = .009; cluster above threshold 242–281 ms after cue onset) and becomes even stronger on cue repetitions 11–15 (**F**; multiple disconnected clusters above threshold between 55–353 ms after cue onset, grey horizontal line; largest cluster above threshold from 245–261 ms, p = .023, black horizontal line).

In sum, we found evidence for freezing of gaze induced by Avoid cues, with lower gaze dispersion on trials with Avoid cues compared to trials with Win cues. This difference in gaze dispersion only emerged with learning the aversive nature of cues.

### Confirmatory analyses: No effect of the arousal manipulation on responses, RTs, or pupil dilation

In pre-registered confirmatory analyses, we tested whether the arousal manipulation (angry/ neutral face primes) affected the proportion of Go/NoGo responses, RTs, or pupil dilation, either as a main effect or in interaction with cue valence, required action, or the trial-by-trial pupil dilation. In brief, the arousal manipulation had no effect on any dependent measure, also not in interaction with other task factors. These findings suggest that the manipulation was ineffective and participants did not process the face primes, neither consciously nor subconsciously. We report the full results of all pre-registered confirmatory as well as additional exploratory analyses in Supplementary Material S05, in which we also discuss reasons for why the manipulation might have been ineffective.

### Exploratory analyses: Stronger trial-by-trial pupil dilations for Go responses, especially to Avoid cues

We measured trial-by-trial fluctuations in pupil dilations for two reasons. Firstly, pupil dilation is commonly interpreted as an index of arousal, which could reveal whether the arousal manipulation via angry/ neutral face primes was effective or not. Secondly, pupil dilation can index other arousal-related processes such as cognitive and physical effort, which might be required on incongruent trials in order to suppress Pavlovian bias. Specifically, cognitive effort is required for conflict detection and resolution on all incongruent trials, i.e., both when inhibiting Go responses to Win cues and when invigoration Go responses to Avoid cues. Higher pupil dilations on incongruent trials should be reflected in an interaction effect between cue valence and required action, with stronger dilations for NoGo than Go responses to Win cues, but stronger dilations for Go than NoGo response to Avoid cues (Fig. 1E). In contrast, physical effort is only required for making Go responses, and particular effort might be required for Go responses to Avoid cues when aversive inhibition induced by Avoid cues has to be overcome. Hence, if pupil dilation specifically reflected physical effort, one would expect a main effect of response, with stronger dilations for Go than NoGo responses, qualified by an interaction effect between response and cue valence, with stronger dilations for Go responses to Avoid than to Win cues, but no difference between Win and Avoid cues for NoGo responses (Fig. 1F). To test these two accounts, we performed exploratory, non-pre-registered analyses of trial-by-trial fluctuations in pupil dilation, operationalized as the maximum dilation between forward mask onset and minimal possible ITI offset (i.e., 2,966 ms after the forward mask) minus the pre-mask baseline (−500 till 0 ms before the mask), as a function of the executed response (Go/ NoGo) and cue valence (Win/ Avoid).

We observed a significant main effect of Go/NoGo responses on trial-by-trial pupil dilation, *b* = 0.112, 95%-CI [0.084, 0.140], χ^2^(1) = 38.769, *p* < .001, with much stronger pupil dilations on trials with Go responses compared to trials with NoGo responses (Fig. 4A, B). Furthermore, there was a significant main effect of valence, *b* = -0.020, 95%-CI [-0.040, - 0.001], χ^2^(1) = 4.007, *p* = .045, with stronger dilation for Avoid than Win cues (Fig. 4A, B). The interaction between performed action and valence was not significant, *b* -0.006, 95%-CI [-0.026, 0.014], χ^2^(1) = 0.356, *p* = .551. However, the pattern displayed in Fig. 4A was suggestive of an interaction effect, with higher dilations for Avoid than Win cues only for Go responses, with this pattern reversing for NoGo responses. This observation was confirmed when using post-hoc *z*-tests, which yielded a significant effect of valence only for Go responses, *z* = 1.974, *p* = .048, but not for NoGo responses, *z* = 0.915, *p* = .360. Complementary analyses with dilations as independent variable and responses as dependent variable reproduced these results (see Supplementary Material S06).

**Figure 4.**
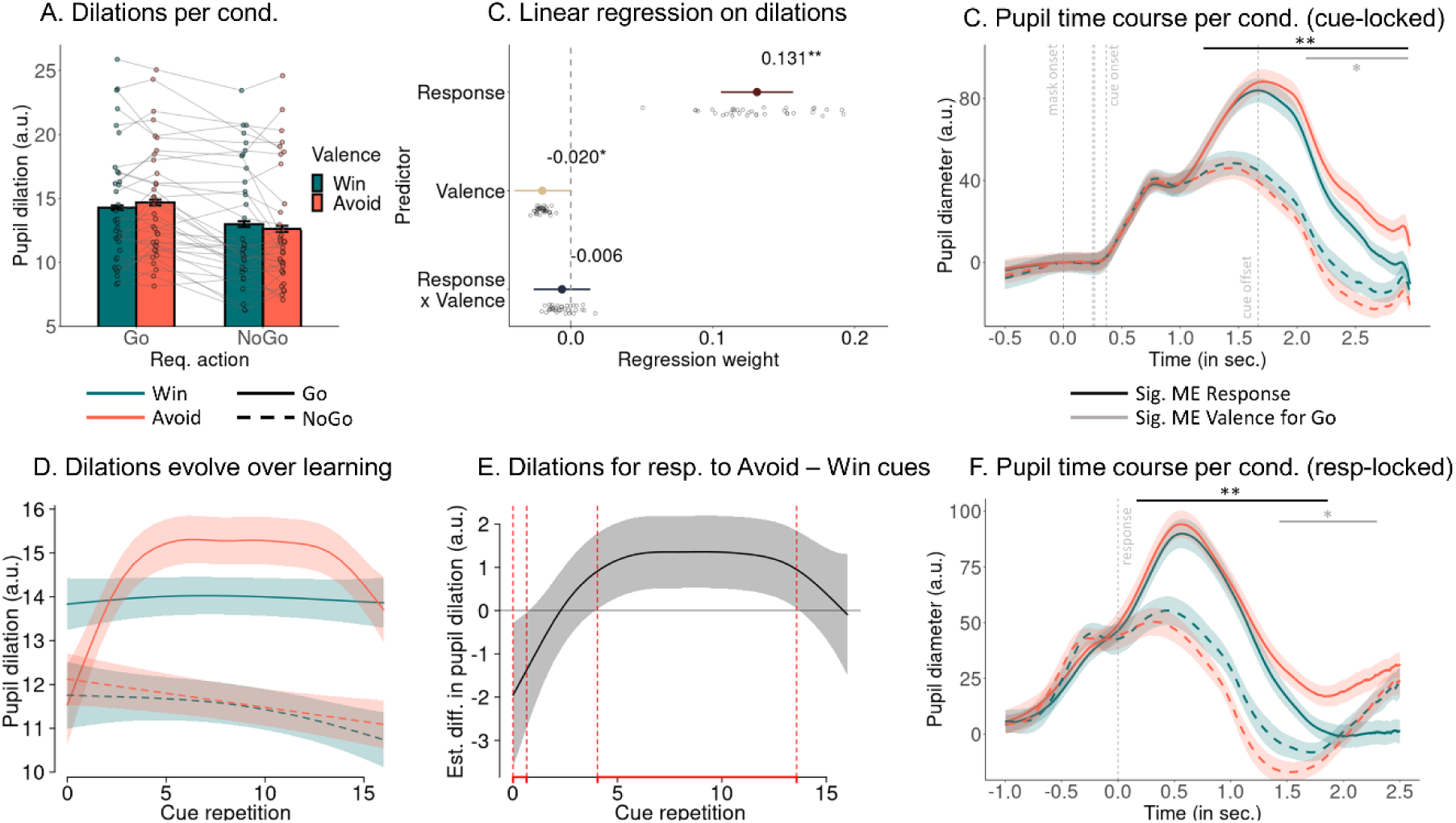
Pupil dilations as a function of the response and cue valence. *Note.* **A.** Mean pupil dilation per response and cue valence. Dilations are significantly higher for Go than NoGo responses and significantly higher for Go responses to Avoid cues than responses to Win cues. **B.** Group-level (colored dot, 95%-CI) and individual-participant (grey dots) regression coefficients from a mixed-effects linear regression of dilations on response, cue valence, and their interaction. There are significant main effects of response and cue valence, but the interaction is not significant. **C**. Pupil time course within a trial locked to forward mask onset per response per cue valence (mean ± SEM across participants; baseline-corrected). Vertical dashed lines indicate the onset of the forward mask (at 0 ms), the prime (at 250 ms), the backwards mask (at 266 ms), the cue onset (at 366 ms), and the cue offset (at 1666 ms). The pupil dilates significantly more on trials with Go responses than on trials with NoGo responses starting 1,190 ms after forward mask onset (black horizontal line). Furthermore, the pupil dilates significantly more sustainedly for responses to Avoid than to Win cues, starting 2,157 ms after forward mask onset (grey horizontal line). See Supplementary Material S08 for a version without baseline correction. **D.** Time course of dilations over cue repetitions (mean ± SE) as predicted from a generalized additive mixed-effects model (GAMM), separated by response and cue valence. Dilations are significantly stronger on trials with Go responses than on trials with NoGo responses through blocks. Furthermore, dilations are significantly stronger for responses to Avoid cues than to Win cues from cue repetition 3 to 13, putatively reflecting heightened effort recruitment on trials with Avoid cues in order to overcome aversive inhibition. **E.** Difference line between dilations on trials with responses to Avoid cues minus Win cues. Areas highlighted in red indicate time windows with significant differences. **F**. Pupil time course within a trial locked to RTs (for NoGo responses: locked to mean RT for responses to Win/ Avoid cues per participant) split by response and cue valence (mean ± SEM across participants; baseline-corrected). Vertical dashed lines indicate the RT (at 0 ms). Note that neither cue nor outcome are systematically locked to RTs and thus not represented in this figure. The pupil dilates significantly more on trials with Go responses than on trials with NoGo responses starting 144 ms after the RT (black horizontal line). Furthermore, the pupil dilates significantly more sustainedly for responses to Avoid than to Win cues, starting 1,516 ms after the RT (grey horizontal line). This latter difference is continues up until the outcome-induced pupil dilation (right end of figure).

In sum these results, are in line with a physical effort account of pupil dilation, with stronger dilations for Go responses and particularly so for responses to Avoid cues which require overcoming aversive inhibition. We followed up on this inconsistency between regression results (Fig. 4B) and the pattern observed when plotting the data (Fig. 4A) with more fine-grained analyses of the pupil time courses during trials.

### Exploratory analyses: More sustained pupil dilations for Go responses to Avoid cues

The previous analyses focused on the trial-by-trial peak of the pupil time course, which is a frequently used summary statistic of the pupil time course. However, it does not capture any variation beyond the peak height, such as condition differences in peak timing or in how sustained the peaks are. Given the above-reported inconsistency between regression results and patterns observed when plotting the data, as a more sensitive measure of condition differences, we tested for such differences in the millisecond-by-millisecond pupil time course using cluster-based permutation tests (Strauch et al., 2022). We corrected for any pre-onset baseline differences (for results without baseline correction, see Supplementary Material S08).

The pupil was significantly wider on trials with Go compared to trials with NoGo responses, *p* < .001, driven by a cluster above threshold from 1,190–2,966 ms after mask onset (i.e., until the end of the testing window, Fig. 4C). Within this time window, the pupil was significantly wider for Go responses to Avoid cues than Go responses to Win cues, *p* = .035, driven by a cluster above threshold from 2,157–2,966 ms (i.e., until the end of the testing window; Fig. 4C). There was no significant difference between NoGo responses to Win and to Avoid cues, p = 1, with no cluster above threshold.

The difference between Go responses to Avoid and Win cues occurred rather late (2,157–2,966 ms), i.e., after the peak of the grand mean pupil response (at 1,591 ms) and after the task cue had already disappeared (i.e., after 1,666 ms). Responses to Avoid cues were most prominently associated with more sustained, rather than stronger pupil dilations. Despite the late time point of this condition difference, due to the sluggishness of the pupil response, it might reflect differences in cognitive processing occurring much earlier, i.e., during cue processing and response selection. Note that this difference occurred much later than differences in gaze dispersion between Avoid and Win cues (i.e., 202–278 ms after cue onset); freezing of gaze and difference and pupil dilation are thus unlikely to confound each other. The difference between responses to Avoid and Win cues also occurred when locking the pupil time course to the response (RT) instead of the mask onset, with stronger dilations during Go than NoGo responses (144–1,934 ms relative to response, *p* < .001) and, within this window, stronger dilations during Go responses to Avoid cues compared to Win cues (1,516–1,934 ms, *p* = .015; Fig. 4F). The latter difference in fact continued up until the outcome-induced dilation as visible in outcome-locked analyses (Supplementary Material S09). For results without baseline-correction, see Supplementary Material S08. For associations between task factors and outcome-locked pupil dilations, see Supplementary Material S09.

Taken together, these results confirmed the pattern of Fig. 4A, with a strong main effect of the executed response on pupil dilation, qualified by particularly strong and sustained pupil dilations for Go responses to Avoid cues. They are line with a physical effort account of pupil dilation (Fig. 1F), with particularly high physical effort recruitment when overcoming aversive inhibition induced by Avoid cues. However, they are not in line with a cognitive effort account of pupil dilation because NoGo responses to Win cues, which require response inhibition and thus cognitive effort, were not associated with strong pupil dilations. Notably, effort requirements should vary over time: they should start once participants have recognized the cue valence and the response required for a given cue, subsequently rise while participants try to counteract their Pavlovian bias, but eventually diminish again once the task is well learned. Hence, next, we tested for condition differences in the dilation time course within task blocks and how these changed with learning.

### Exploratory analyses: Stronger dilations for Go responses to Avoid cues arise and vanish again with learning

Pavlovian biases and their effective suppression depends on participants learning the cue valence and required response, recognizing the demand for physical effort recruitment in order to suppress aversive inhibition. Participants are initially unaware of the correct response or cue valence and thus do not recruit particular physical effort to invigorate Go responses to Avoid cues (see learning curve per cue in Fig. 2A). As they become more certain about which response to perform, physical effort recruitment might increase, particular for the cues they have learned to be Avoid cues. With further learning, response selection becomes more certain and the instrumental system dominates the Pavlovian system, requiring less effort with increasing practice. As a result of these two antagonistic trends, an inverted-U shape, with maximal effort recruitment at intermediate stages of learning, could be expected. To test this hypothesis, we fit generalized additive mixed-effects models to participants’ trial-by-trial pupil dilations, testing whether the time course of pupil dilations (modeled via the cue repetition, 1– 16) differed between conditions.

The model suggested significantly higher pupil dilations for Go than NoGo responses throughout learning (repetitions 1–16), parametric term *t*(5.54, 7.45) = 14.585, *p* < .001, smooth term *F*(1.32, 1.56) = 2.340, *p* = .165. Furthermore, pupil dilations were significantly stronger for Go responses to Avoid cues than to Win cues between cue repetitions 3 till 13 (and lower around cue repetition 1), parametric term *t*(5.75, 7.67) = 3.039, *p* = .002, smooth term *F*(3.39, 4.16) = 3.483, *p* = .007 (Fig. 4D, E). Note how this time course is mirroring the learning curve for Go-to-Aoid cues (Fig. 2A). See Supplementary Material S07 for results showing that this pattern held independently of other factors affecting pupil dilations for Go responses, such as accuracy, response speed, and response repetition.

In sum, these results indicate that stronger dilations for Go responses to Avoid compared to Win cues occurred specifically at intermediate stages of learning, when overcoming aversive inhibition has become driven by past experiences, but not sufficiently practiced yet.

## Discussion

In this study, we tested whether subliminally induced arousal modulated Pavlovian biases in an orthogonalized Motivational Go/NoGo Task, and whether measured fluctuations in arousal indexed via pupil dilation reflected processes involved in inhibiting those biases. Win vs. Avoid cues induced strong Pavlovian biases in both responses and RTs, with faster and more Go responses to Win compared to Avoid cues. The aversive nature of the Avoid cues even elicited a brief “freezing” of gaze, with less gaze dispersion from the center for Avoid compared to Win cues. However, neither responses, nor RTs, nor pupil dilations showed any effect of the arousal priming manipulation, suggesting that this manipulation—though successfully used in past research (Allen et al., 2016)—was ineffective (for a discussion of these findings, see Supplementary Material S05).

Exploratory analyses showed that measured fluctuations in trial-by-trial pupil dilation reflected participants’ responses, specifically the physical effort they recruited to exert a Go response: Stronger dilations occurred on trials with Go responses, with particularly strong and sustained dilations for responses to Avoid cues that were performed against the hindrance induced by Pavlovian biases. There were no comparable pupil responses on trials on which participants inhibited responses to Win cues, which also required the suppression of Pavlovian biases. Thus, pupil dilations do not reflect response conflict or cognitive effort associated with resolving such conflict on “incompatible” trials, but selectively the physical effort required for overcoming the aversive inhibition induced by Avoid cues. Notably, stronger pupil dilations for Go responses to Avoid cues only emerged with learning, indicative that they do not reflect motor processes per se, but the specific physical effort demands required. Taken together, these results are in line with an account of pupil dilation reflecting physical (but not cognitive) effort investment. Beyond previous literature on conflict detection and response suppression in the context of Pavlovian biases (Cavanagh et al., 2013; Swart et al., 2018), these results highlight another cognitive capacity required to manage Pavlovian biases, namely response invigoration against adversities.

### Freezing of gaze by Avoid cues

Avoid cues robustly reduced response rates and slowed reaction times. Note that strong aversive Pavlovian biases are usually absent in variants of the Motivational Go/NoGo Task that separate Pavlovian cues and the response window in time (Guitart-Masip et al., 2012; Queirazza, Steele, Krishnadas, Cavanagh, & Philiastides, 2023). Hence, the instruction to respond immediately to the appearance to the cue seems necessary for observing these biases in behavior. Only in such a variant, it becomes possible to study the mechanisms by which participants overcome an aversive bias.

Beyond Pavlovian biases in responses and RTs, we also found cue valence to affect gaze position: During the cue presentation, participants’ gaze showed less dispersion from the center of the screen for Avoid cues compared to Win cues in a time range around 200–280 ms after cue onset, with differences becoming stronger with learning. It is notable that, compared to previous studies reporting freezing of gaze (Merscher & Gamer, 2024; Merscher et al., 2022; Rösler & Gamer, 2019), the reduction of gaze dispersion observed in this study was temporally and spatially very constrained. This difference likely arises from differences in the experimental set-up. Previous studies encouraged participants to visually explore photos of natural scenes while they prepared for a button press in order to prevent an electric shock. In contrast, in our task, participants were instructed to maintain fixation at the center of the screen while an aversive cue signaling the chance of losing points was presented. This might explain why the freezing of gaze effect in our study was much smaller than in previous studies, reflecting differences in only a few pixels instead of hundreds of pixels, with a duration of less than 100 ms.

We also considered the possibility that reduced gaze dispersion for Avoid cues did not reflect automatic, “Pavlovian” effects, but rather deliberate, strategic response adjustments for such cues. Notably, the freezing effect on gaze occurred within less than 300 ms after cue onset, which is faster than typical EEG correlates of task engagement (e.g., the P300 event-related potential). Differences in pupil dilation, which we interpret as reflecting differences in physical effort exertion, occurred more than 2,000 ms later, suggesting that effort-related processes were separate from this early freezing of gaze. Taken together, freezing of gaze likely reflects early, automatic Pavlovian processes, which might be followed by later deliberate changes in task engagement to counteract the freezing, rather than vice versa. Beyond high-level differences in task engagement, differences between cue valence conditions are unlikely to be driven by low-level visual properties, since we matched all cues for average luminance and counterbalanced the assignment of cues to task conditions across participants. Lastly, we observed that this freezing of gaze phenomenon was not yet present on the first five occurrences of a cue when cue valence had not been learned, but emerged only in the middle of blocks when participants had become aware of the cue valence. In sum, these considerations suggest that the observe freezing of gaze is specifically related to the learned valence of the cues and reflects early automatic processes rather than later strategic processes that aim to overcome aversive inhibition.

Our results corroborate recent evidence that freezing does not merely affect limb movements, but also the oculomotor system. Past research has shown that the chance to gain rewards speeds up saccades (Manohar et al., 2015; Shadmehr, Reppert, Summerside, Yoon, & Ahmed, 2019; Tachibana & Hikosaka, 2012), a process sensitive to dopamine and likely implemented by the direct pathway of the basal ganglia (Grogan, Sandhu, Hu, & Manohar, 2020; Kawagoe, Takikawa, & Hikosaka, 1998). Conversely, the indirect pathway in the basal ganglia seems responsible for the suppression of eye movements in presence of low-value objects (Amita & Hikosaka, 2019; Kim, Amita, & Hikosaka, 2017), a role it might also play for negative events such as aversive cues and threats of punishment. Overall, these findings suggest a more principled role of the basal ganglia in modulating the vigor of eye movements as a function of incentives (Park, Coddington, & Dudman, 2020; Turner & Desmurget, 2010). Our results contribute to this literature by showing how the oculomotor system can give insights in reward- and punishment processing not only in animals, but also in humans (Shadmehr et al., 2019).

### Pupil dilation reflects physical effort expenditure in a graded fashion

Apart from gaze, also pupil dilations reflected aspects of the Motivational Go/NoGo Task. The biggest effect on pupil dilations was caused by responses, with much stronger pupil dilations for Go than for NoGo responses. This finding concords with a large body of literature reporting stronger pupil dilations under movement preparation, movement execution, and effort exertion (Beatty, 1982; Bijleveld, Custers, & Aarts, 2009; da Silva Castanheira et al., 2020; Kurniawan, Grueschow, & Ruff, 2021; van der Wel & van Steenbergen, 2018; Zénon, Sidibé, & Olivier, 2014). However, it is still an open question which specific processes drive these previously observed response-related pupil dilations. Some studies have argued that such response-related pupil dilations constitute an epiphenomenon of motor movements, i.e. an signal that qualitatively reflects whether a movement is executed or not in an all-or-nothing fashion (Richer & Beatty, 1985; Richer et al., 1983). Alternatively, pupil dilations have been suggested to reflect the effort that is required to execute a response in a more graded, continuous fashion (da Silva Castanheira et al., 2020; van der Wel & van Steenbergen, 2018). Our results concur with the latter interpretation given that we found particularly strong and sustained dilations for responses to Avoid cues, which is plausible since Avoid cues induce aversive inhibition, which might require particular effort to overcome. This effect was not constant as expectable for a motor artifact, but changed systematically with learning. Further supplementary analyses revealed particularly strong dilations also for slow responses, which might reflect conflict and effort recruitment, as well. We discuss these findings in the following. Lastly, we found no comparable increase in pupil dilations for NoGo responses to Win cues, arguing against a cognitive effort account of pupil dilation.

Higher pupil dilations during responses to Avoid than to Win cues specifically reflect effort demands, which dynamically change as a function of learning. Differences between Avoid and Win cues occurred specifically in the middle of each block, i.e., after participants were made aware of the cue valence, but before they had fully learned the correct response. At the beginning of each block, new cues were introduced, and until participants had experienced a win or loss of points, they could not know the cue valence. Thus, until the aversive nature of Avoid cues had been experienced, these cues did not induce aversive inhibition nor did they motivate additional effort recruitment. Similarly, little effort was required at the end of blocks when the instrumental learning system had acquired reliable action values that were unlikely to be “swayed” by Pavlovian biases (Dorfman & Gershman, 2019). Additionally, at the end of each block, the experienced rate of punishments had become lower due to increased accuracy, which in turn might have lowered the aversive value of the cues and reduced aversive inhibition. In summary, effort was recruited only after the aversive nature of cues had become clear, but only until responses to them became well-learned, concurring with the interpretation of pupil dilation as reflecting effort recruited to overcome aversive inhibition.

Another piece of evidence suggesting that pupil dilations reflect effort recruitment in a continuous fashion is the finding that dilations were stronger for slower compared to faster responses (see Supplementary Materials S06 and S07). Slow responses are often interpreted as reflecting action selection against difficulties, involving effortful cognitive control to resolve conflict (Frank, 2006). The link between dilations and responses was particularly strong for incorrect Go responses (to NoGo cues), which were slower than correct responses (to Go cues), implying that these do not reflect “impulsive” errors, but rather deliberate choices made in spite of previous feedback providing evidence against Go responses. Such slow, incorrect responses might have required particularly high levels of physical effort to trigger a Go response against competing instrumental processes suggesting a NoGo response. Notably, this type of effort was not associated with relatively faster, but slower responses, reflecting situations where eventual Go responses result sequentially from conflict detection and subsequent effort recruitment. Hence, in the context of this task, the recruitment of “physical effort” or “vigor” was not in the service of speeding up responses, but instead of executing responses in the first place.

One might wonder how our findings relate to the literature that links pupil dilation to cognitive effort associated with response conflict or task switching (van der Wel & van Steenbergen, 2018). Studies of these phenomena have usually employed paradigms that feature choices between several “Go” responses. Boosting a slow, more controlled response over an automatic, prepotent response might require inhibiting the latter, but could also be implemented by invigorating of the former, which likely involves some form of physical effort. In contrast, tasks requiring pure response inhibition, e.g., classic Go/ NoGo tasks, are experienced as cognitively effortful (Dixon & Christoff, 2012), but arguably require little physical effort. In such tasks, pupil dilations are smaller for effortful, controlled NoGo responses compared to prepotent Go responses, which suggests that response conflict and cognitive effort associated with the inhibition of prepotent responses are not sufficient to drive pupil dilations (Schacht et al., 2010; Van der Molen et al., 1989). Together with our results, these findings suggest that pupil dilation is more tightly linked to the invigoration of slow, controlled, deliberate responses, rather than response conflict or cognitive effort per se. This link is only visible in specific paradigms such as the Motivational Go/NoGo Task that create response conflict, but dissociate physical effort from cognitive effort requirements. Note however that some paradigms requiring cognitive effort, but no particular physical effort, such as mental arithmetic or problem solving tasks, did observe increased pupil dilations (Hess & Polt, 1964; Kahneman, 1973). Future research will have to identify which specific task features induced these dilations. What we conclude from this study is that response conflict and inhibition alone, such as NoGo responses to Win cues in this task, are not sufficient to drive strong pupil dilations.

### Putative neural mechanisms of aversive biases and their suppression

The link between pupil dilation and physical effort is also corroborated by direct recordings from neurons in the locus coeruleus, the major source of noradrenaline in the brain, which strongly correlates with pupil dilations (Joshi, Li, Kalwani, & Gold, 2016; Strauch et al., 2022). Such direct recordings in monkeys have linked noradrenaline levels to physical effort expenditure (Bornert & Bouret, 2021; Varazzani, San-Galli, Gilardeau, & Bouret, 2015). Specifically, one study recorded activity from the substantia nigra and locus coeruleus, the primary sources of dopamine and noradrenaline, while monkeys performed a reward/ effort trade-off task involving a grip forcer (Varazzani et al., 2015). Dopamine reflected expected value and required effort before response onset, while noradrenaline reflected the grip force actually exerted during responses, which was also reflected in pupil diameter. These findings suggest a link between noradrenaline, pupil dilation, and physical effort expenditure that is likely shared across species.

While several studies have reported a correlation between pupil diameter and activity of the locus coeruleus (Joshi & Gold, 2019; Joshi et al., 2016; Murphy, O’Connell, O’Sullivan, Robertson, & Balsters, 2014), this link has recently come under debate (Megemont, McBurney-Lin, & Yang, 2022). Pupil size also correlates with the trial-by-trial BOLD signal activity in other brain stem nuclei, specifically the dopaminergic ventral tegmental area and substantia nigra, at least during rest (Lloyd, de Voogd, Mäki-Marttunen, & Nieuwenhuis, 2023). It might be interesting to consider the possibility that the response-induced modulation of pupil dilation in this study in fact reflects dopaminergic activity (Varazzani et al., 2015; Walton & Bouret, 2018). In line with this hypothesis, one of our past studies (Algermissen et al., 2022) found the same pattern observed in pupil dilations in this study—a strong main effect of responses, with a particular strong signal for responses to Avoid cues—in the dorsal striatal BOLD signal, which replicated previous patterns of VTA and striatal BOLD signal (Guitart-Masip et al., 2012) and was recently replicated again (Queirazza et al., 2023). The same study found striatal BOLD to be correlated with midfrontal theta power. Other studies have found pupil diameter to be related to midfrontal theta power (Dippel et al., 2017; Lin et al., 2018) and the P300, an evoked potential likely generated by stimulus-locked oscillations in the theta range (de Gee, Correa, Weaver, Donner, & van Gaal, 2021; Murphy, Robertson, Balsters, & O’Connell, 2011; Nieuwenhuis, Aston-Jones, & Cohen, 2005). In sum, striatal BOLD, midfrontal theta power, and pupil diameter might all reflect the same underlying signal, which however is not noradrenergic, but dopaminergic in nature.

If pupil dilation indexes dopaminergic processes in the striatum, these might be directly related to the “unfreezing” of gaze. Striatal dopamine has been suggested to enhance the contrast of cortical action representations against background noise, facilitating their selection (Nicola, Woodward Hopf, & Hjelmstad, 2004). The early freezing evoked by activation of the indirect pathway, visible in reduced gaze dispersion, might be directly counteracted by the dopaminergic enhancement of action representations in the direct pathway, visible in pupil dilation, which would bridge the two main findings of this study. Interestingly, while freezing of gaze became stronger over time within a task block, potentially reflecting more automatic retrieval of the valence of Avoid cues, the incremental pupil dilation associated with Go responses to Avoid cues showed an inverted-U shaped time course, diminishing towards the end of task blocks. This observation tentatively suggests that counteracting the Pavlovian biases also becomes more automatized over time, requiring less physical effort with practice. Future research will have to use brain imaging methods to directly investigate the time course of such effort-related processes in the striatum.

The hypothesis that pupil dilation (at least partially) reflects dopaminergic processes in the striatum is in line with recent accounts of the role of dopamine in motivating action. Specifically, it has been proposed that the striatum evaluates whether recruiting additional effort to invigorate a candidate response option will lead to increases in expected reward, i.e., it computes the “value of work” (Hamid et al., 2016; Mohebi et al., 2019; Syed et al., 2016; Westbrook, Frank, & Cools, 2021). Further work by the same authors has suggested that dopamine reflects the control or “agency” an individual experiences over its environment, reflecting whether it is worth investing effort to try to increase reward rates or not (Hamid, 2021; Hamid, Frank, & Moore, 2021). The value of work is particularly high under response conflict, when boosting slow, controlled response options over fast, automatic, prepotent response options can make a difference for whether a correct response is executed (and a reward obtained) or not. Hence, pupil dilation might give insights into this underexplored facet of cognitive control.

### Limitations

The present study features a number of limitations and points at new directions for future research. Firstly, the unsuccessful subliminal manipulation motivates the question whether a supraliminal manipulation might be more successful. However, for supraliminally presented stimuli, even more care must be taken in matching their visual properties, and condition differences could reflect differences in low-level stimulus processing. Furthermore, consciously perceived emotional stimuli can induce high-level changes in response strategy, i.e., demand characteristics (Mahlberg et al., 2021), which necessitates the use of an elaborate and effective cover story. Lastly, the presence of strong response-related transients in the pupil data might potentially camouflage more subtle stimulus-induced effects. Other physiological measures of arousal such as heart rate and skin conductance might be more suitable to measure the effects of supraliminally presented arousing stimuli (Hashemi et al., 2019; Klaassen et al., 2021). However, these measures need much longer measurement periods, requiring a slower trial structure.

In the present data, pupil diameter peaked around 1,600 ms after stimulus onset and returned to baseline around 3,000 ms, showing a slower time course than previous studies on pupil dilation (Hoeks & Levelt, 1993) and warranting care when pre-registering analysis windows. The time course of the pupil dilation might vary considerably as a function of the task structure and should be measured in pilot data before pre-registering a definite analysis window.

### Summary

In summary, our results shed new light on the effects of aversive cues on motor behavior (eye and hand movements) and on the effortful counter-mechanisms recruited to overcome aversive inhibition. Aversive cues reduced response rates, slowed responses and reduced gaze dispersion (“freezing of gaze”). Over time, participants learned to counteract this aversive Pavlovian bias and make Go responses even to aversive cues. These responses were associated with particularly strong and sustained pupil dilations, which we interpret as reflecting additional physical effort recruitment in order to overcome aversive inhibition. While previous literature has primarily focused on how impulsive responding to Win cues can be suppressed (Cavanagh et al., 2013; Swart et al., 2018), this study sheds light on the opposite end of Pavlovian biases, namely how humans can invigorate responding against factors holding them back. Future studies could use pupillometry in the context of aversive inhibition to further probe this underexplored facet of cognitive control.

## Acknowledgements

We thank Pim Klee, Gert Proper, David Renjaän, and Karlijn Tummers for assistance with data collection. We thank Micah Allen for advice on stimulus preparation and the subliminal priming procedure.

## Supplementary Material S01: Overview of results from all mixed-effects regression models

Here, we report an overview over all major statistical results reported in the main text and the Supplementary Materials S05 and S06. For details on how mixed-effects regression were performed, see the Methods section of the main text.

**Table S01.**
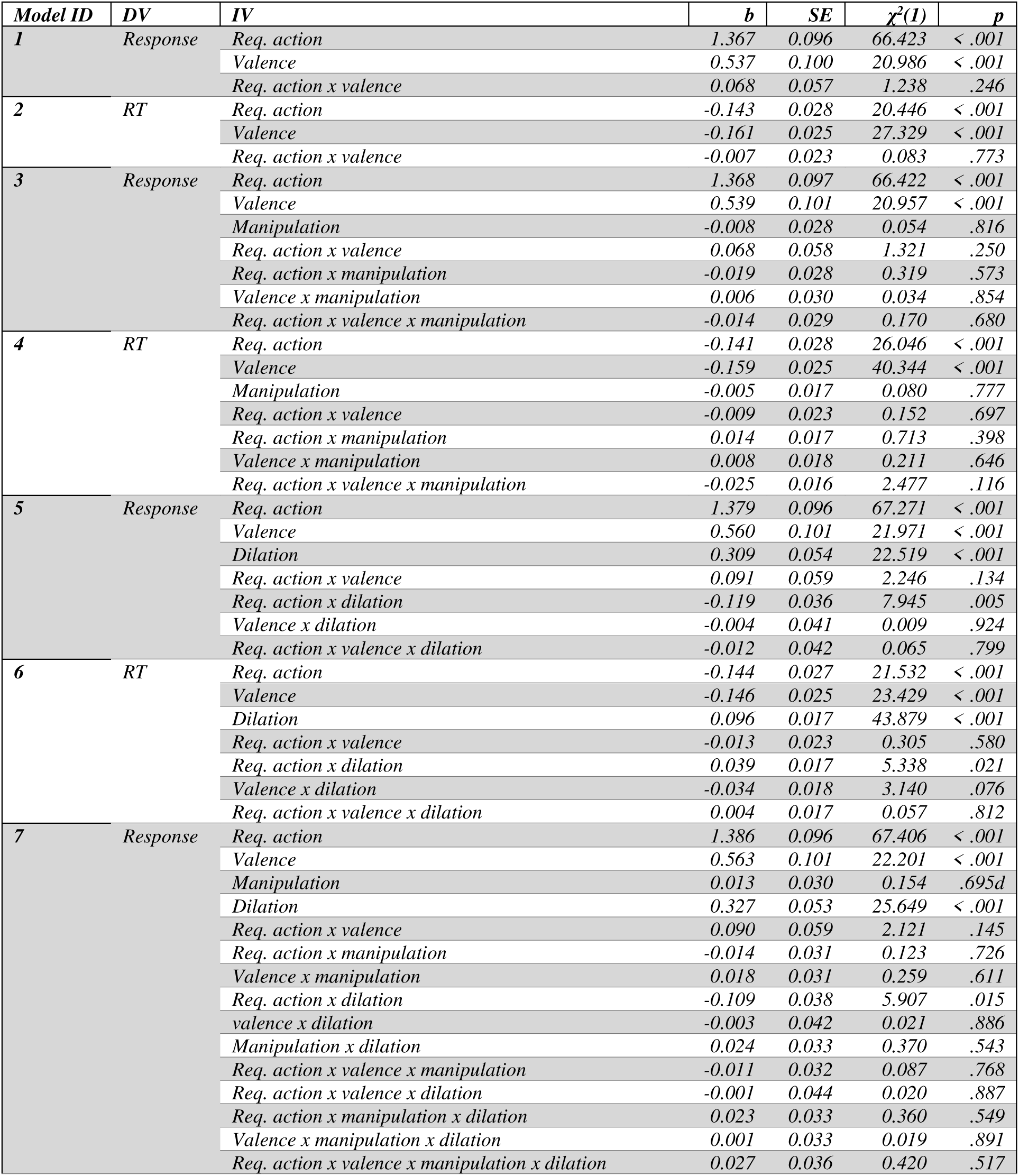
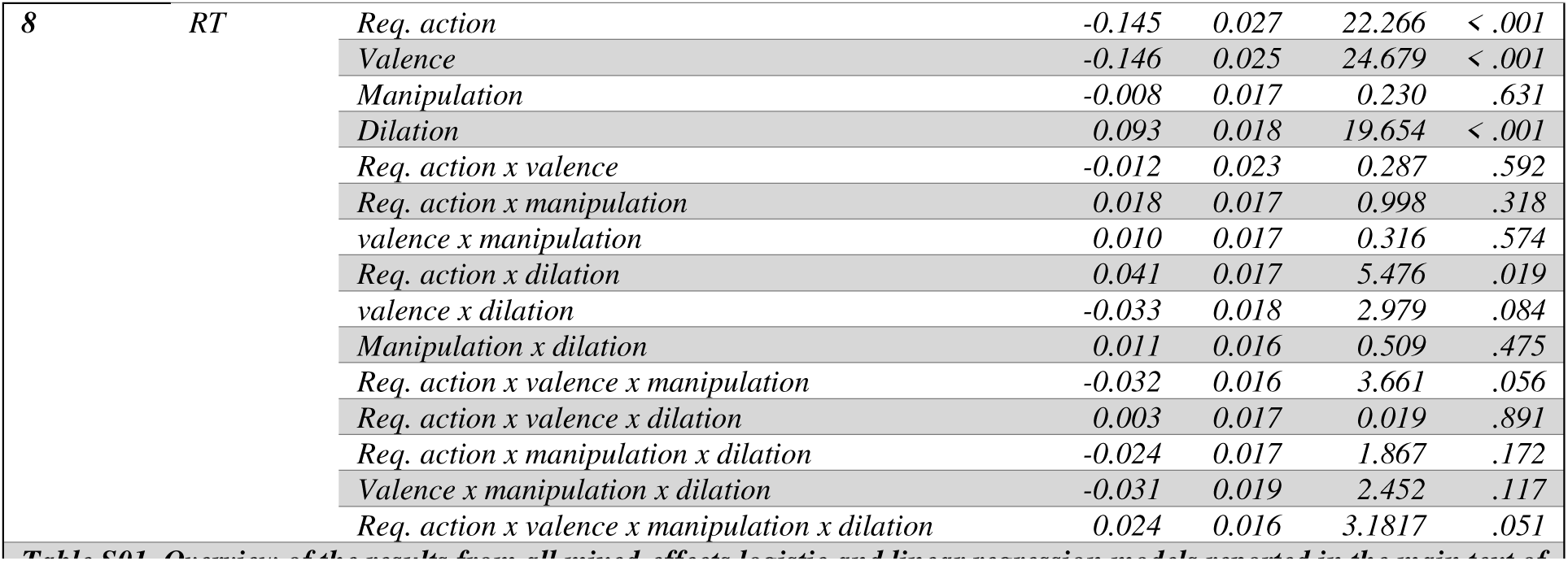
Overview of the results from all mixed-effects logistic and linear regression models reported in the main text of the manuscript.

| *Model ID* | *DV* | *IV* | *b* | *SE* | $\chi^2(I)$ | *p* |
| --- | --- | --- | --- | --- | --- | --- |
| **1** | *Response* | *Req. action* | 1.367 | 0.096 | 66.423 | < .001 |
|  |  | *Valence* | 0.537 | 0.100 | 20.986 | < .001 |
|  |  | *Req. action x valence* | 0.068 | 0.057 | 1.238 | .246 |
| **2** | *RT* | *Req. action* | -0.143 | 0.028 | 20.446 | < .001 |
|  |  | *Valence* | -0.161 | 0.025 | 27.329 | < .001 |
|  |  | *Req. action x valence* | -0.007 | 0.023 | 0.083 | .773 |
| **3** | *Response* | *Req. action* | 1.368 | 0.097 | 66.422 | < .001 |
|  |  | *Valence* | 0.539 | 0.101 | 20.957 | < .001 |
|  |  | *Manipulation* | -0.008 | 0.028 | 0.054 | .816 |
|  |  | *Req. action x valence* | 0.068 | 0.058 | 1.321 | .250 |
|  |  | *Req. action x manipulation* | -0.019 | 0.028 | 0.319 | .573 |
|  |  | *Valence x manipulation* | 0.006 | 0.030 | 0.034 | .854 |
|  |  | *Req. action x valence x manipulation* | -0.014 | 0.029 | 0.170 | .680 |
| **4** | *RT* | *Req. action* | -0.141 | 0.028 | 26.046 | < .001 |
|  |  | *Valence* | -0.159 | 0.025 | 40.344 | < .001 |
|  |  | *Manipulation* | -0.005 | 0.017 | 0.080 | .777 |
|  |  | *Req. action x valence* | -0.009 | 0.023 | 0.152 | .697 |
|  |  | *Req. action x manipulation* | 0.014 | 0.017 | 0.713 | .398 |
|  |  | *Valence x manipulation* | 0.008 | 0.018 | 0.211 | .646 |
|  |  | *Req. action x valence x manipulation* | -0.025 | 0.016 | 2.477 | .116 |
| **5** | *Response* | *Req. action* | 1.379 | 0.096 | 67.271 | < .001 |
|  |  | *Valence* | 0.560 | 0.101 | 21.971 | < .001 |
|  |  | *Dilation* | 0.309 | 0.054 | 22.519 | < .001 |
|  |  | *Req. action x valence* | 0.091 | 0.059 | 2.246 | .134 |
|  |  | *Req. action x dilation* | -0.119 | 0.036 | 7.945 | .005 |
|  |  | *Valence x dilation* | -0.004 | 0.041 | 0.009 | .924 |
|  |  | *Req. action x valence x dilation* | -0.012 | 0.042 | 0.065 | .799 |
| **6** | *RT* | *Req. action* | -0.144 | 0.027 | 21.532 | < .001 |
|  |  | *Valence* | -0.146 | 0.025 | 23.429 | < .001 |
|  |  | *Dilation* | 0.096 | 0.017 | 43.879 | < .001 |
|  |  | *Req. action x valence* | -0.013 | 0.023 | 0.305 | .580 |
|  |  | *Req. action x dilation* | 0.039 | 0.017 | 5.338 | .021 |
|  |  | *Valence x dilation* | -0.034 | 0.018 | 3.140 | .076 |
|  |  | *Req. action x valence x dilation* | 0.004 | 0.017 | 0.057 | .812 |
| **7** | *Response* | *Req. action* | 1.386 | 0.096 | 67.406 | < .001 |
|  |  | *Valence* | 0.563 | 0.101 | 22.201 | < .001 |
|  |  | *Manipulation* | 0.013 | 0.030 | 0.154 | .695d |
|  |  | *Dilation* | 0.327 | 0.053 | 25.649 | < .001 |
|  |  | *Req. action x valence* | 0.090 | 0.059 | 2.121 | .145 |
|  |  | *Req. action x manipulation* | -0.014 | 0.031 | 0.123 | .726 |
|  |  | *Valence x manipulation* | 0.018 | 0.031 | 0.259 | .611 |
|  |  | *Req. action x dilation* | -0.109 | 0.038 | 5.907 | .015 |
|  |  | *valence x dilation* | -0.003 | 0.042 | 0.021 | .886 |
|  |  | *Manipulation x dilation* | 0.024 | 0.033 | 0.370 | .543 |
|  |  | *Req. action x valence x manipulation* | -0.011 | 0.032 | 0.087 | .768 |
|  |  | *Req. action x valence x dilation* | -0.001 | 0.044 | 0.020 | .887 |
|  |  | *Req. action x manipulation x dilation* | 0.023 | 0.033 | 0.360 | .549 |
|  |  | *Valence x manipulation x dilation* | 0.001 | 0.033 | 0.019 | .891 |
|  |  | *Req. action x valence x manipulation x dilation* | 0.027 | 0.036 | 0.420 | .517 |
| 8 | RT | Req. action | -0.145 | 0.027 | 22.266 | < .001 |
|  |  | Valence | -0.146 | 0.025 | 24.679 | < .001 |
|  |  | Manipulation | -0.008 | 0.017 | 0.230 | .631 |
|  |  | Dilation | 0.093 | 0.018 | 19.654 | < .001 |
|  |  | Req. action x valence | -0.012 | 0.023 | 0.287 | .592 |
|  |  | Req. action x manipulation | 0.018 | 0.017 | 0.998 | .318 |
|  |  | valence x manipulation | 0.010 | 0.017 | 0.316 | .574 |
|  |  | Req. action x dilation | 0.041 | 0.017 | 5.476 | .019 |
|  |  | valence x dilation | -0.033 | 0.018 | 2.979 | .084 |
|  |  | Manipulation x dilation | 0.011 | 0.016 | 0.509 | .475 |
|  |  | Req. action x valence x manipulation | -0.032 | 0.016 | 3.661 | .056 |
|  |  | Req. action x valence x dilation | 0.003 | 0.017 | 0.019 | .891 |
|  |  | Req. action x manipulation x dilation | -0.024 | 0.017 | 1.867 | .172 |
|  |  | Valence x manipulation x dilation | -0.031 | 0.019 | 2.452 | .117 |
|  |  | Req. action x valence x manipulation x dilation | 0.024 | 0.016 | 3.1817 | .051 |

## Supplementary Material S02: Overview of means and standard deviations of responses and RTs per task condition

**Table S02.** Means and standard deviations of Go/NoGo responses across participants per required action x valence condition.

| Req. Act. | Responses |  |  |  |
| --- | --- | --- | --- | --- |
|  | Go | Go | NoGo | NoGo |
| Valence | Win | Avoid | Win | Avoid |
| Mean | 0.875 | 0.759 | 0.410 | 0.216 |
| SD | 0.124 | 0.122 | 0.258 | 0.096 |

**Table S03.** Means and standard deviations of Go/NoGo responses across participants per required action x valence x prime condition.

| Req. Act. | Responses |  |  |  |  |  |  |  |
| --- | --- | --- | --- | --- | --- | --- | --- | --- |
|  | Go | Go | Go | Go | NoGo | NoGo | NoGo | NoGo |
| Valence | Win | Win | Avoid | Avoid | Win | Win | Avoid | Avoid |
| Prime | High | Low | High | Low | High | Low | High | Low |
| Mean | 0.871 | 0.880 | 0.754 | 0.763 | 0.414 | 0.405 | 0.215 | 0.217 |
| SD | 0.131 | 0.124 | 0.138 | 0.124 | 0.258 | 0.269 | 0.106 | 0.102 |

**Table S04.** Means and standard deviations of reaction times across participants per required action x valence condition.

| Req. Act. | RTs |  |  |  |
| --- | --- | --- | --- | --- |
|  | Go | Go | NoGo | NoGo |
| Valence | Win | Avoid | Win | Avoid |
| Mean | 0.641 | 0.707 | 0.707 | 0.756 |
| SD | 0.071 | 0.076 | 0.122 | 0.103 |

**Table S05.** Means and standard deviations of reaction times across participants per required action x valence x prime condition.

| Req. Act. | RTs |  |  |  |  |  |  |  |
| --- | --- | --- | --- | --- | --- | --- | --- | --- |
|  | Go | Go | Go | Go | NoGo | NoGo | NoGo | NoGo |
| Valence | Win | Win | Avoid | Avoid | Win | Win | Avoid | Avoid |
| Prime | High | Low | High | Low | High | Low | High | Low |
| Mean | 0.641 | 0.641 | 0.713 | 0.702 | 0.711 | 0.704 | 0.738 | 0.771 |
| SD | 0.081 | 0.067 | 0.078 | 0.083 | 0.131 | 0.123 | 0.131 | 0.116 |

## Supplementary Material S03: Effect of response accuracy and cue valence on RTs over time

In the main text, we report main effects of accuracy/ required action on RTs, with slower RTs for incorrect responses (to NoGo cues) than for correct responses (to Go cues), and of cue valence on RTs, with slower responses to Avoid cues than to Win cues. Here, we fitted additional generalized additive mixed-effects models to test whether these effects changed over trials (i.e., cue repetitions).

For the time course of RTs across trials per condition, see Fig. S01A. Again, we found significantly slower RTs for incorrect responses (to NoGo cues) than for correct responses (to Go cues) for cue repetitions 3–13, parametric term *t*(3.19, 0.11) = 3.726, *p* < .001, smooth term *F*(3.03, 3.73) = 2.894, *p* = .023 (Fig. S01B). The fact that incorrect responses were significantly slower already on the third repetition of a cue reveals that participants had some (partial) awareness of the correct response already after the first few trials. Differences disappeared towards the end of blocks. Note that accuracy increased over time, with fewer and fewer trials contributing to the time courses of incorrect responses. Changes in the number of trials contributing to conditions likely explain why the disappearance of this effect seems to be at odds with FigS01A, in which the time course for incorrect responses to Avoid cues (red dashed line) continues to become slower. Accuracy for NoGo-to-Avoid trials is very high, especially late in the blocks (see Fig. 2A in the main text), with mostly NoGo responses in this condition and very few incorrect Go responses that contribute to the RT time course. These few incorrect NoGo-to-Avoid trials are down-weighted relative to the more frequent incorrect NoGo-to-Win trials.

Furthermore, we found significantly slower RTs for responses to Avoid cues than for responses to Win cues for cue repetitions 2–16, parametric term *t*(3.36, 0.11) = 12.851, *p* < .001, smooth term *F*(3.31, 4.06) = 8.790, *p* < .001(Fig. S01C). Pavlovian biases in RTs emerged already on the 2^nd^ repetition of a cue and continued until the end of blocks.

In sum, these results reflect that effects of required action/ accuracy and cue valence on RTs emerged after the first few trial repetitions. While the accuracy effect disappeared towards the end of blocks (with few incorrect Go responses to NoGo cues), the cue valence effect persisted till the end.

**Figure S01.**
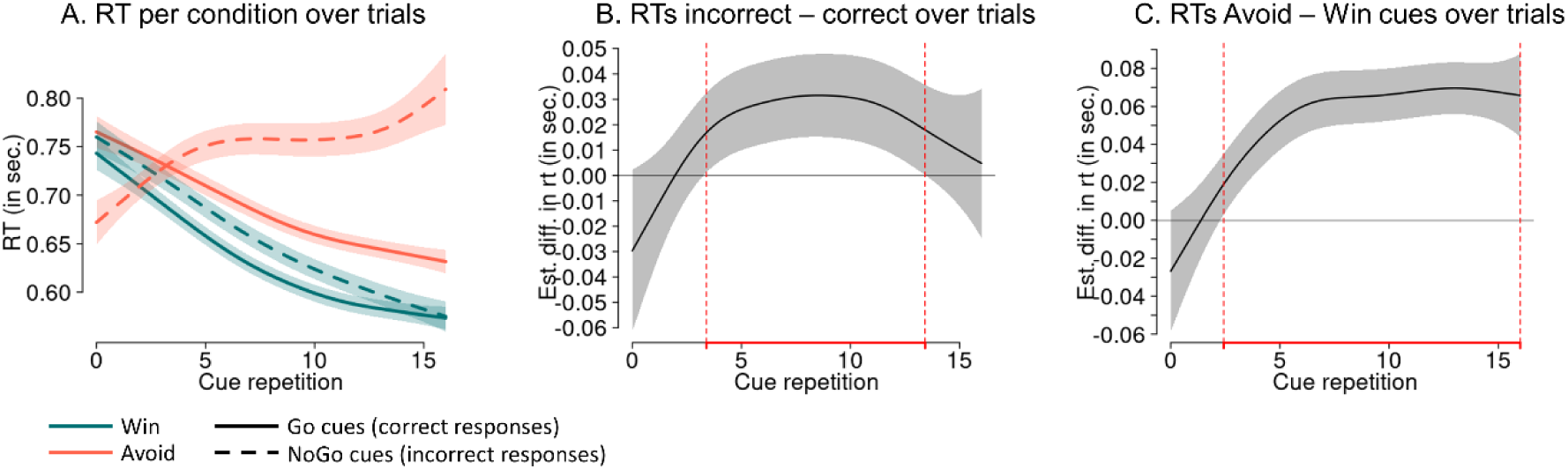
Effect of accuracy and cue valence on RTs over cue repetitions. **A.** Time course of dilations over cue repetitions (mean ± SE) as predicted from a generalized additive mixed-effects model (GAMM), separated by accuracy/ required action and cue valence. RTs are significantly slower for incorrect than correct responses from cue repetition 3 to 13 and significantly slower for responses to Avoid than Win cues from cue repetition 2 to 16 (i.e., the end of blocks). **B.** Difference line between RTs for incorrect responses minus correct responses, with significant differences from cue repetition 3 to 13. Areas highlighted in red indicate time windows with significant differences. **C.** Difference line between RTs for responses Avoid cues minus Win cues, with significant differences from cue repetition 2 to 16.

## Supplementary Material S04: Correlations of the effects of cue valence on responses and RTs with questionnaires

In line with the exploratory analysis plans in mentioned in our pre-registration, we extracted the per-participant coefficients (fixed plus random effects) for (a) the effect of cue valence on responses and RTs (Pavlovian bias), (b) the effect of the arousal manipulation on responses and RTs, and (c) the effect of pupil dilation on responses and RTs. We then computed correlations of these coefficients with trait anxiety (STAI, Form Y-2, 20 items) (Spielberger, Gorssuch, Lushene, Vagg, & Jacobs, 1983) and the five sub-scales negative urgency, lack of perseveration, lack of premeditation, sensation seeking, and positive urgency of the UPPS-P Impulsive Behavior Scale (short version, 20 items) (Cyders, Littlefield, Coffey, & Karyadi, 2014). One might plausibly hypothesize that impulsivity is related to the Pavlovian bias since many impulsive behaviors can be conceptualized as automatic, cue-triggered behaviors.

See Fig. S02 for scatterplots of all bivariate associations. None of these correlations were significant at α = .05 (uncorrected), providing no evidence for the strength of the Pavlovian bias in responses or RTs being related to either trait anxiety or sub-facets of impulsivity. Note that these analysis are underpowered to detect correlations of small-to-moderate size: With *N* = 35, we have 80% power to detect correlations of |*r|* > 0.45, and only correlations of |*r|* > 0.33 (50% power) will become significant.

**Figure S02.**
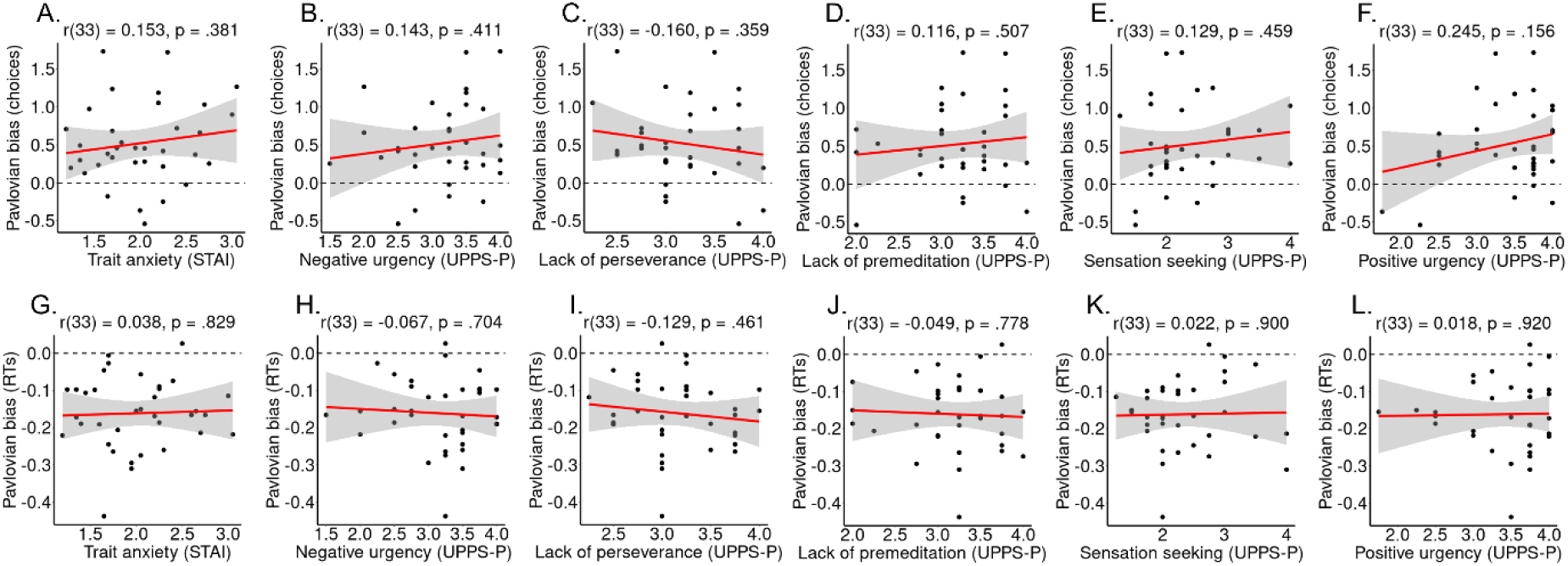
Association of trait anxiety and various sub-facets of trait impulsivity with the effect of valence on responses and RTs. Correlations between the effect of valence on responses (**A–F**) and on RTs (**G-L**), reflecting Pavlovian biases, and trait anxiety (**A**, **G**), negative urgency (**B, H**), lack of perseverance (**C, I**), lack of premeditation (**D, J**), sensation seeking (**E, K**), and positive urgency (**F, L**). Black dots represent per-participant scores, the red line the best-fitting regression line, the grey shade the 95%-confidence interval. None of the displayed correlations is significant at α = .05.

## Supplementary Material S05: No effect of arousal manipulation on the responses, RTs, and pupil dilation

Here, we report the full results from confirmatory pre-registered and additional exploratory analyses of a responses, RTs, and trial-by-trial pupil dilation as function of the subliminal arousal manipulation (angry vs. neutral face primes). In brief, there was no effect of the arousal manipulation on responses, RTs, or pupil dilation, neither as a main effect nor in interaction with other task factors or measured pupil dilation. In pre-registered, confirmatory models, we fit a mixed-effects logistic regression model with responses (Go/NoGo) as dependent variable and required action (Go/ NoGo), cue valence (Win/ Avoid), arousal priming manipulation (angry/ neutral face), as well as all possible interactions between them as independent variables. Furthermore, we fit an equivalent exploratory linear regression model with RTs as dependent variable. As a initial, exploratory manipulation check, we fit an equivalent linear regression model with trial-by-trial pupil dilation as dependent variable.

As a initial manipulation check, we performed exploratory analyses testing for an effect of the subliminal arousal manipulation (angry vs. neural face primes) on trial-by-trial-pupil dilation. We expected higher pupil dilations for angry compared to neural faces, reflecting heightened arousal induced by angry faces. In a mixed-effects model regression trial-by-trial pupil dilations onto the conditions of the arousal manipulation, there was no effect of the manipulation, *b =* -0.003, 95%-CI [-

0.022, 0.017], χ^2^(1) = 0.071, *p* = .790 (Fig. S03A), providing no evidence for the subliminal arousal manipulation affecting arousal as index by pupil diameter. In addition to this regression model, we performed two more exploratory manipulation checks, testing for differences between subliminally presented angry vs. neutral faces at any time point during a trial as well as at any time point during learning.

To test for any effect of the arousal manipulation on pupil dilation at any time point within a trial, we computed the raw pupil time course per condition (high vs. low arousal) for every participant and then the average per condition across participants. A cluster-based permutation test yielded no significant difference at any time point (no cluster above the cluster-forming threshold of |t| > 2), suggesting again no effect of the arousal manipulation on pupil dilation (Fig. S03B).

Furthermore, we tested whether the arousal manipulation affected pupil dilations at any time point within a block using generalized additive mixed-effects models. There was no difference in the trial-by-trial time course of pupil dilations between high-arousal and low-arousal trials, linear term *t*(5.75, 7.61) = 0.252, p = .801, smooth term *F*(2.42, 2.98) = 1.757, *p* = .170, suggesting again no effect of the arousal manipulation on pupil dilation (Fig. S03C). In sum, none of the performed manipulation checks suggested any evidence for the subliminal arousal manipulation affecting arousal as indexed by trial-by-trial fluctuations in pupil dilations.

**Figure S03.**
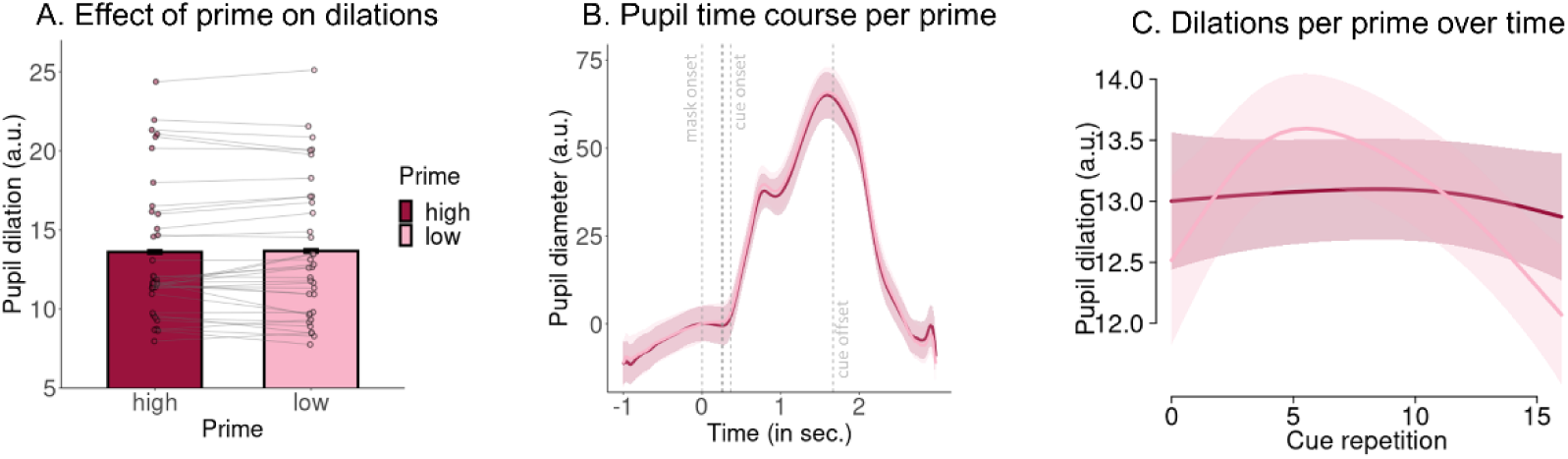
Effect of arousal manipulation on pupil dilation. **A.** Mean pupil dilation per level of the arousal priming manipulation (whiskers are ±SEM across participants, dots indicate individual participants). There is no effect of the arousal priming manipulation on pupil dilations. **B.** Pupil time course within a trial (mean ± SE; baseline-corrected) separately for high vs. low arousal condition. Vertical dashed lines indicate the onset of the forward mask (at 0 ms), the prime (at 250 ms), the backwards mask (at 266 ms), the cue onset (at 366 ms), and the cue offset (at 1666 ms). There is no significant difference (no cluster above cluster-forming threshold). **C.** Time course of dilations over cue repetitions (mean ± SE) as predicted from a generalized additive mixed-effects model (GAMM), separated by arousal condition. There is no significant difference in pupil dilation between conditions at any time point.

Next, we proceed with our main confirmatory, pre-registered analyses. As a first set of confirmatory, pre-registered analyses, we extended the regression model in the main text fitting responses as a function of required action and cue valence by adding the arousal priming manipulation (high/ low, i.e., angry/ neutral face stimulus) and all higher-order interactions possible. Neither the main effect of the arousal priming manipulation, *b* = -0.008, 95%-CI [-0.063, 0.047], χ^2^(1) = 0.054, *p* = .816, nor the 2-way interaction between the priming manipulation and cue valence, *b* = 0.006, 95%-CI [-0.052, 0.065], χ^2^(1) = 0.034, *p* = .854, nor the 3-way interaction between the priming manipulation, cue valence, and required action, *b* = -0.014, 95%-CI [-0.071, 0.043], χ^2^(1) = 0.170, *p* = .680, was significant, providing no evidence for any effect of the priming manipulation on responses (Fig. S04A-C).

Fitting an equivalent model to RTs, neither the main effect of the arousal priming manipulation, *b* = -0.005, 95%-CI [-0.038, 0.028], χ^2^(1) = 0.073, *p* = .787, nor the 2-way interaction between the priming manipulation and cue valence, *b* = 0.008, 95%-CI [-0.026, 0.043], χ^2^(1) = 0.197, *p* = .657, nor the 3-way interaction between the priming manipulation, cue valence, and required action, b = -0.025, 95%-CI [-0.055, 0.006], χ^2^(1) = 2.354, *p* = .125, was significant, providing no evidence for any effect of the arousal priming manipulation on responses (Fig. S04D-F). Taken together, none of the confirmatory analyses provided any evidence for the arousal priming manipulation affecting behavior.

As a third set of confirmatory analyses, we fit a regression model with required action, cue valence, the arousal priming manipulation, trial-by-trial pupil dilation, and all higher-order interactions possible. There was no significant 4-way interaction on either responses, *b* = 0.027, 95%-CI [-0.044, 0.098], χ^2^(1) = 0.420, *p* = .517, nor RTs, b = 0.024, 95%-CI [-0.006, 0.055], χ^2^(1) = 3.817, *p* = .051, again providing no evidence for an effect of the arousal priming manipulation, also not as a function of the trial-by-trial pupil dilation.

**Figure S04.**
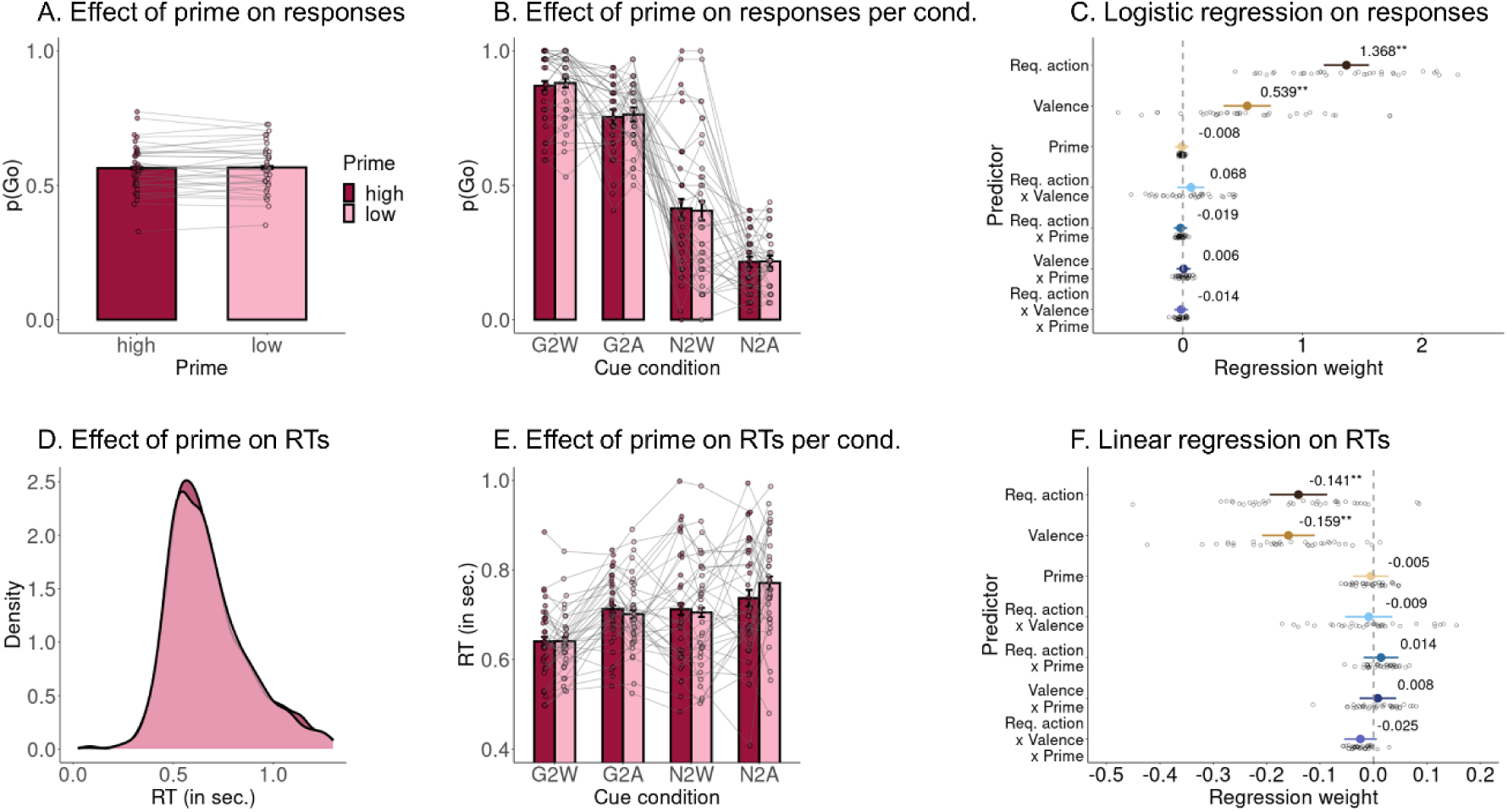
Effect of the arousal priming manipulation on responses and RTs. **A.** Proportion of Go responses for high (angry face) and low arousal (neutral face) priming manipulation (whiskers are ± SEM across participants, dots indicate individual participants). There is no effect of the manipulation on responses. **B**. Proportion of Go responses for high and low arousal priming manipulation separately per cue condition. There is no effect of the manipulation on responses for any condition. **C.** Group-level (colored dot, 95%-CI) and individual-participant (grey dots) regression coefficients from a mixed-effects logistic regression of responses on required action, cue valence, the arousal priming manipulation, and all higher-order interactions. None of the terms involving the arousal priming manipulation is significant. **D**. Distribution of raw RTs separately per arousal priming manipulation level. There is no difference between both levels. **E**. Mean RTs for high and low arousal priming manipulation separately per cue condition. There is no effect of the manipulation on RTs for any condition. **F.** Group-level and individual-participant regression coefficients from a mixed-effects linear regression of RTs on required action, cue valence, the arousal priming manipulation, and all higher-order interactions. None of the terms involving the arousal priming manipulation is significant.

As an exploratory check, we tested whether individual differences in the effects of the arousal manipulation on responses, RTs, and pupil dilation were correlated, i.e., whether only those participants who showed an effect on pupil dilation also showed an effect on behavior. For this purpose, we fit regression models with the manipulation as sole independent variable and responses, RTs, and dilations and dependent variables, extracted the per-participants coefficients (fixed + random effects), and correlated them. Neither the per-participants effects of the manipulation on dilations and responses, *r*(33) = - 0.202, *p* = .243 (Fig. S05A), nor the effects on dilations and RTs, *r*(33) = 0.121, *p* = .487 (Fig. S05B), were significantly correlated, providing no evidence for systematic individual differences in the effect of the arousal manipulation on behavior and physiology.

**Figure S05.**
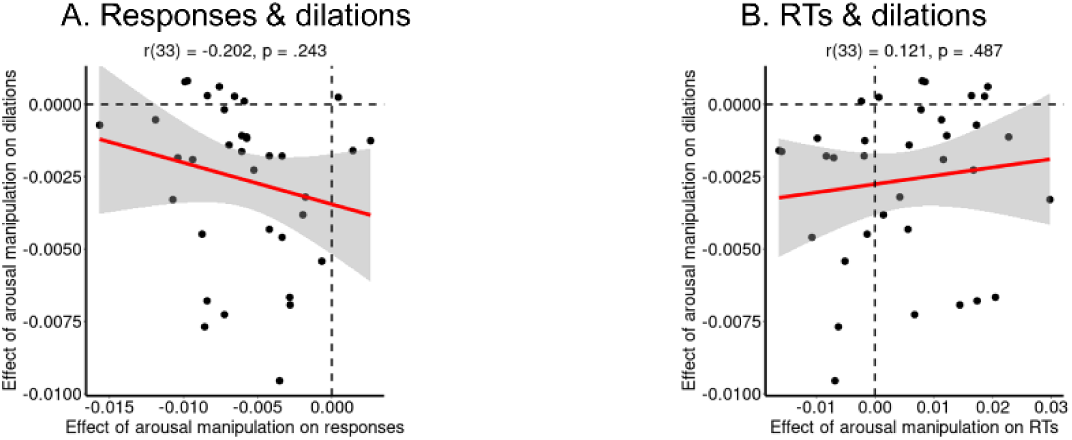
Correlations between the effects of the arousal manipulation on response, RTs, and pupil dilations. **A.** Correlation between the effect of the arousal manipulation on responses and on trial-by-trial pupil dilation. Black dots represent per-participant scores, the red line the best-fitting regression line, the grey shade the 95%-confidence interval. The correlation is not significant. **B**. Correlation between the effect of the arousal manipulation on RTs and on trial-by-trial pupil dilation. The correlation is not significant.

As a final set of exploratory analyses, we tested whether individual differences in the effect of the arousal priming manipulation on responses or RTs were correlated with individual differences in self-reported anxiety and/or impulsivity. One might plausibly hypothesize that trait anxiety would be associated with a stronger effect of the exogenously induced arousal on responses and RTs. See Fig. S06 for scatterplots of all bivariate associations. Neither the effect of the arousal priming manipulation on responses nor on RTs was correlated with trait anxiety or impulsivity across participants, providing no evidence for the strength of the effect of induced arousal on responses and RTs being related to either trait anxiety or sub-facets of impulsivity. Note that these analysis are underpowered to detect correlations of small-to-moderate size: With *N* = 35, we have 80% power to detect correlations of |*r|* > 0.45, and only correlations of |*r|* > 0.33 (50% power) will become significant.

**Figure S06.**
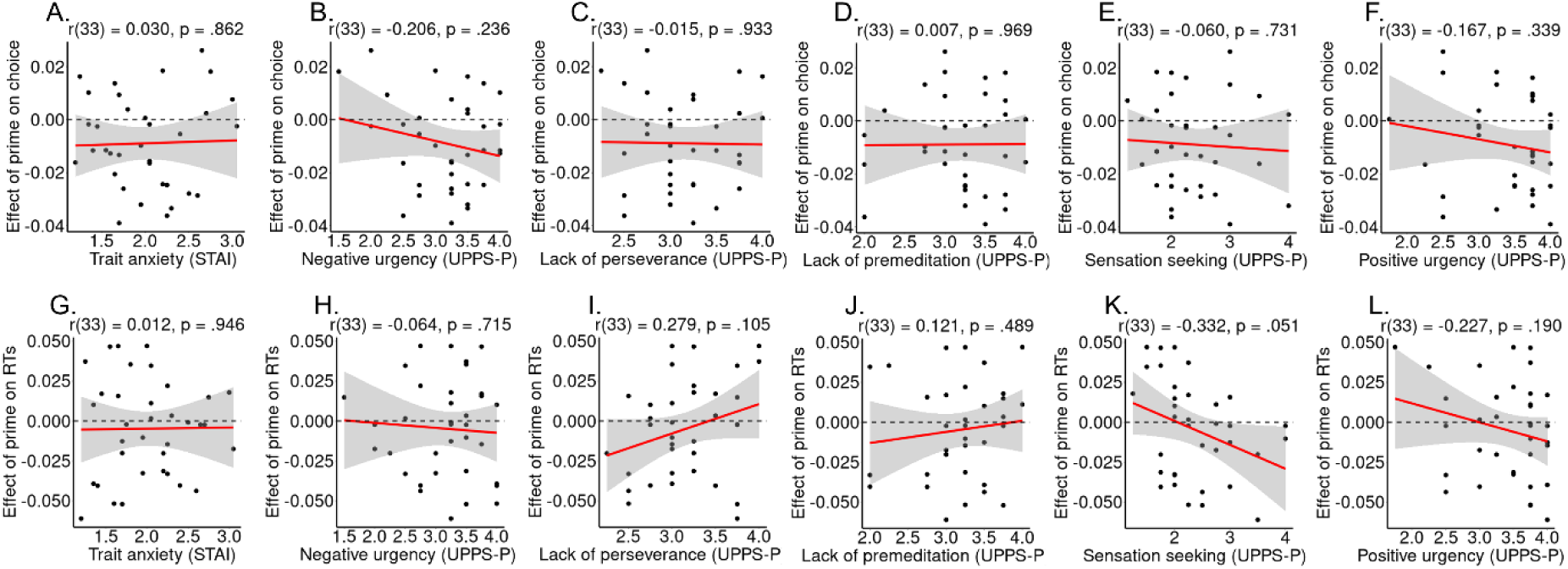
Association of trait anxiety and various sub-facets of trait impulsivity with the effect of the arousal manipulation on responses on RTs. Correlations between the effect of the subliminal arousal manipulation on responses (**A–F**) and on RTs (**G-L**), and trait anxiety (**A**, **G**), negative urgency (**B, H**), lack of perseverance (**C, I**), lack of premeditation (**D, J**), sensation seeking (**E, K**), and positive urgency (**F, L**). Black dots represent per-participant scores, the red line the best-fitting regression line, the grey shade the 95%-confidence interval. None of the displayed correlations is significant at α = .05.

In this study, we used a previously established manipulation that subliminally presented faces with angry or neutral faces to induce high vs. low arousal (Allen et al., 2016). We did not observe any effects on responses, RTs, or pupil dilation. Confidence intervals and raw data plots indicated that the effect of the manipulation on all dependent measures was close to zero (Fig. S04), with little variation across participants, providing strong evidence for a null effect. Hence, although this procedure has been used successfully in the past (and proven seemingly effective in data from four pilot participants we had collected initially), it was unsuccessful in this study. Likely, the presentation duration was too short for participants to (even subliminally) process the emotional faces. The pupillometry data in particular provides strong evidence that no processing of the emotional faces occurred. This failure to use a subliminal manipulation to induce arousal aligns with other recent reports calling into question the effectiveness of subliminal manipulations reported in the literature (Mudrik & Deouell, 2022). Several cognitive processes previously reported to occur without awareness, including emotional face processing, might in fact require awareness (Mudrik & Deouell, 2022; Skora, Livermore, Dienes, Seth, & Scott, 2023; Vadillo, Malejka, Lee, Dienes, & Shanks, 2022). It is possible that subsets of participants who perceived stimuli supraliminally did in fact drive seemingly subliminal effects in past studies (Skora et al., 2023).

## Supplementary Material S06: Association of pupil dilation with responses and RTs

In the main text of the manuscript, we report exploratory regression models with pupil dilations as dependent variable and task factors such as the performed response or cue valence as independent variables. However, a complementary approach, which we in fact mentioned in our pre-registration, are regression models with dependent and independent variables inverted, i.e., using responses and RTs as dependent variables and trial-by-trial fluctuations in pupil dilation as independent variable, including interactions between pupil dilation and the task factors cue valence and required action.

Note that these analyses are harder to interpret with regard to cognitive effort and physical effort accounts of pupil dilation since they do not directly contrast the relevant task conditions. Nonetheless, the results from these analyses are largely in line with the results reported in main text, with high pupil dilations being associated with Go responses and with particularly with slower RTs.

As confirmatory models, we fit a mixed-effects logistic regression model with responses (Go/NoGo) as dependent variable and required action (Go/ NoGo), cue valence (Win/ Avoid), trial-by-trial pupil dilations, as well as all possible interactions between them as independent variables. Furthermore, we checked whether including RTs as an additional independent variable or including the interaction between RTs and trial-by-trial pupil dilation as an additional independent variable led to different results. Adding those additional independent variables lead to identical conclusions. These models are thus not separately reported here. In case of interactions between dilations and task conditions, in order to better understand these effects, we combined required action and cue valence into a single “cue condition” variable and fit a model with dilation, cue condition, and their interaction. We then tested for differences between conditions in the slope of the dilation effect using z-tests provided by the emmeans package in R, which corrects for multiple comparisons using the Tukey method.

In these analyses, we extended the regression model reported in the main text fitting responses as a function of required action and cue valence by adding the trial-by-trial pupil dilation and all possible higher-order interactions. There was a significant main effect of dilation, *b* = 0.309, 95%-CI [0.203, 0.414], χ^2^(1) = 22.519, *p* < .001, with overall stronger dilations for Go responses (Fig. S07A, C). Furthermore, there was a significant interaction between dilations and required action, *b* = -0.119, 95%-CI [-0.19, -0.049], with a stronger relationship for incorrect responses (to NoGo cues) than correct responses (to Go cues) (Fig. S07B, C). In contrast, neither the 2-way interaction between dilations and cue valence *b* = -0.004, 95%-CI [-0.085, 0.077], χ^2^(1) = 0.009, *p* = .923, nor the 3-way interaction between dilations, cue valence, and required action was significant, *b* = -0.012, 95%-CI [-0.095, 0.071], χ^2^(1) = 0.065, *p* = .799, providing no evidence for pupil dilation modulating the effect of Pavlovian biases on responses (Fig. S07C). To better understand the 2-way interaction between dilations and required action, we fit a follow-up model combining required action and cue valence into a single “cue condition” variable with 4 levels (Go-to-Win, Go-to-Avoid, NoGo-to-Win, NoGo-to-Avoid). The 2-way interaction between dilations and conditions was significant, χ^2^(1) = 8.977, *p* = .030. The association between dilation and the probability of making a Go response was positive in all conditions, with a marginally significant tendency for a stronger link for NoGo-to-Win cues than for Go-to-Win cues (*z* = 2.409, *p* = 0.076) and for Go-to-Avoid cues (*z* = 2.406, *p* = .076), but overall no significant difference between pairs of conditions. See Supplementary Material S07 for evidence that the stronger dilations for incorrect responses (to NoGo cues) than correct responses (to Go cues) occurred due to the former being overall slower, with the association between pupil dilations and accuracy vanishing when controlling for RT differences. In sum, Go responses were associated with stronger pupil dilation, which was especially the case for responses to NoGo cues (i.e., when those responses were incorrect and slow), but there was no evidence for dilations modulating the Pavlovian bias in responses.

An equivalent model fit to RTs yielded a significant main effect pupil dilation, *b* = 0.096, 95%-CI [0.062, 0.129], χ^2^(1) = 43.879, *p* < .001, with stronger dilations being associated with slower RTs, and a significant 2-way interaction between dilations and required action, *b* = 0.039, 95%-CI [0.007, 0.072], χ^2^(1) = 5.338, *p* = .021, with a stronger link between dilations and RTs for Go cues compared to NoGo cues (Fig. S07E, F). The 2-way interaction between dilations and cue valence was only marginally significant, *b =* -0.034, 95%-CI [-0.070, 0.002], χ^2^(1) = 3.140, *p* = .076, tending towards a stronger link between dilations and RTs for Avoid compared to Win cues (Fig. S07D, F). The 3-way interaction between dilations, cue valence, and required action was no significant, *b* = 0.004, 95%-CI [-0.03, 0.038], χ^2^(1) = 0.057, *p* = .812 (Fig. S07F).

To better understand the (marginally) significant 2-way interactions, i.e., test explicitly whether effects were driven by only one of the four cue conditions, we again fit a follow-up model combining required action and cue valence into a single “cue condition” variable with four levels. The 2-way interaction between dilation and cue condition was significant, χ^2^(1) = 9.603, *p* = .023. The association between dilations and RTs was positive in all conditions, strongest in the Go-to-Avoid condition, and weakest in the NoGo-to-Win condition, with this difference being significant, *z* = 3.339, *p* = .005, but none of the other comparisons being significant *p* > .108. See Supplementary Material S07 for evidence that the association between strong pupil dilations and slow RTs also explains the association between pupil dilations and incorrect responses. In sum, stronger dilations were associated with slower RTs, especially so for Go cues and Avoid cues, exacerbating the Pavlovian bias in RTs.

In sum, pupil dilations were stronger for Go responses, particularly for slow and for incorrect responses. The link between pupil dilation and RTs was stronger for Avoid compared to Win cues. However, this effect was only marginally significant and appeared to be driven by responses to Go-to-Avoid (rather than NoGo-to-Avoid) cues (though note that, for the latter, Go responses were incorrect, and thus only few trials with RTs were available). Taken together, these results are in line with a physical effort account of pupil dilation, with stronger dilations for Go than NoGo responses, overall, and particularly strong dilations in situations that require particular physical effort, such as responses to Avoid cues, (rare) incorrect responses, and particularly slow responses.

**Figure S07.**
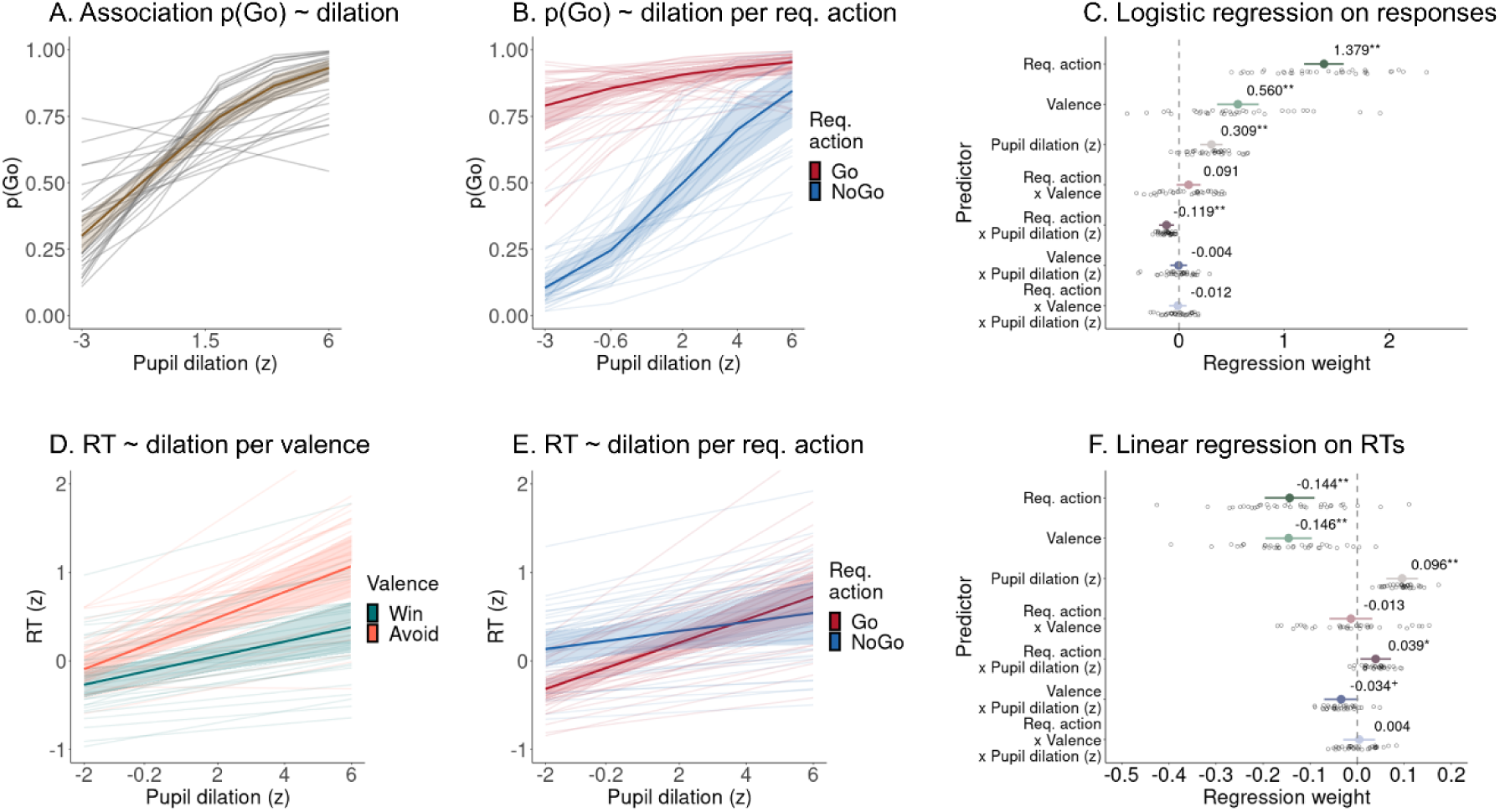
Effect of the trial-by-trial pupil dilation on responses and RTs. **A.** Proportion of Go responses as a function of trial-by-trial pupil dilation as predicted from a mixed-effects logistic regression model (colored line and shades are the group-level association + 95%-CIs; individual lines are the predictions for each individual participant). Go responses are associated with stronger pupil dilations. **B**. Predictions from panel A split per required action. The association between responses and pupil dilations is significantly stronger for (incorrect) responses to NoGo cues than for (correct) responses to Go cues. This difference between incorrect and correct responses is likely due to the former being slower than the latter (see Supplementary Material S07). **C.** Group-level (colored dot, 95%-CI) and individual-participant (grey dots) regression coefficients from a mixed-effects logistic regression of responses on required action, cue valence, pupil dilation, and all higher-order interactions. The main effect of pupil dilation and its interaction with required action are significant. **D**. Predictions of RTs from a mixed-effects logistic regression model based on trial-by-trial pupil dilation separately for Win and Avoid cues. Stronger pupil dilations are associated with slower responses. This relationship is marginally significantly stronger for Avoid than for Win cues. **E**. Predictions of RTs from a mixed-effects logistic regression model based on trial-by-trial pupil dilation separately for Go and NoGo cues. The association between pupil dilation and RTs is significantly stronger for (correct) responses to Go cues than (incorrect) responses to NoGo cues. **F.** Group-level and individual-participant regression coefficients from a mixed-effects linear regression of RTs on required action, cue valence, pupil dilation, and all higher-order interactions. The main effect of pupil dilation as well as its interaction with required action is significant; its interaction with cue valence is marginally significant.

As a final set of exploratory analyses, we tested whether individual differences in the link between pupil dilation and responses or RTs were correlated with individual differences in self-reported anxiety and/or impulsivity. One might plausibly hypothesize that trait anxiety would be associated with a stronger effect of endogenous arousal fluctuations as reflected in trial-by-trial pupil diameter on responses and RTs. See Fig. S08 for scatterplots of all bivariate associations. The only correlation significant at a level of α = .05 (uncorrected) was between trait anxiety and the effect of dilations on RTs, with more anxious individuals showing a weaker link between trial-by-trial pupil dilation (supposedly reflecting fluctuations in endogenous arousal) and RTs. None of the other correlations were significant, providing no evidence for the strength of the effect of endogenous arousal on responses and RTs being related to either trait anxiety or sub-facets of impulsivity. Note that these analysis are underpowered to detect correlations of small-to-moderate size: With *N* = 35, we have 80% power to detect correlations of |*r|* > 0.45, and only correlations of |*r|* > 0.33 (50% power) will become significant.

**Figure S08.**
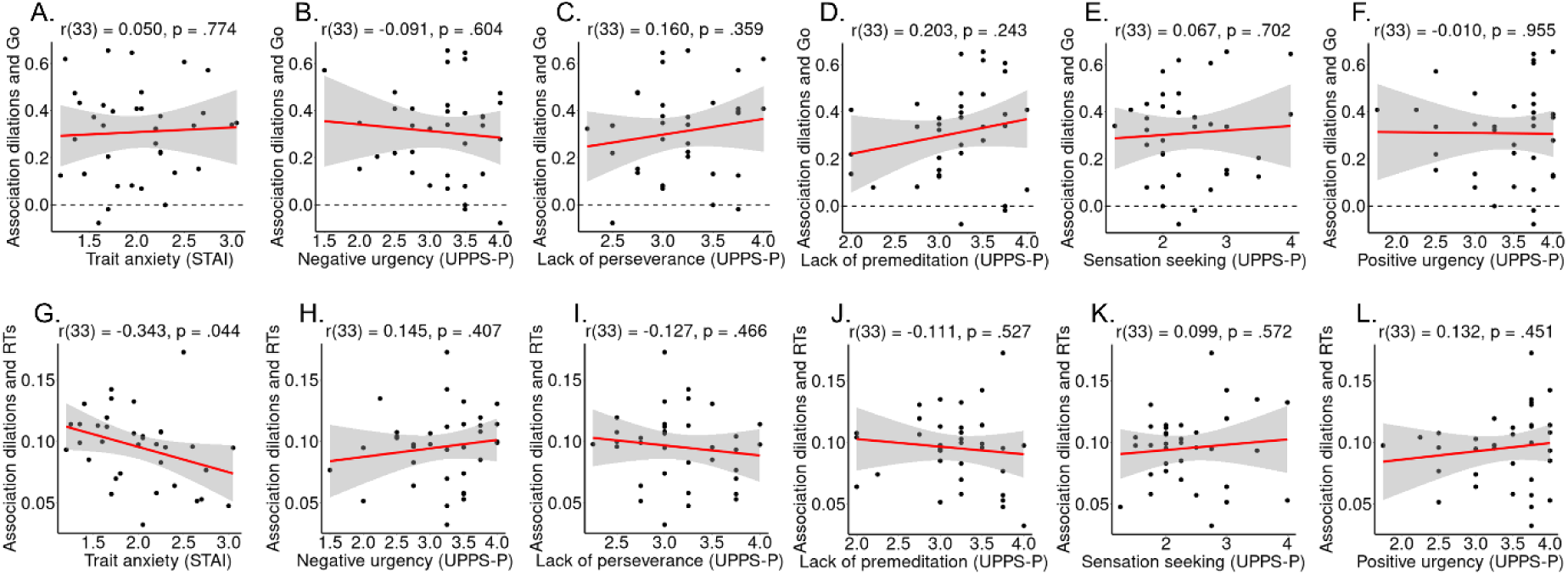
Association of trait anxiety and various sub-facets of trait impulsivity with the effect of trial-by-trial pupil dilation on responses on RTs. Correlations between the effect of trial-by-trial pupil dilation on responses (**A–F**) and on RTs (**G-L**), and trait anxiety (**A**, **G**), negative urgency (**B, H**), lack of perseverance (**C, I**), lack of premeditation (**D, J**), sensation seeking (**E, K**), and positive urgency (**F, L**). Black dots represent per-participant scores, the red line the best-fitting regression line, the grey shade the 95%-confidence interval. The only correlation significant at a level of α = .05 (uncorrected) is between trait anxiety and the effect of dilations on RTs, with more anxious individuals showing a weaker link between trial-by-trial pupil dilation (supposedly reflecting fluctuations in endogenous arousal) and RTs.

## Supplementary Material S07: Higher pupil dilations for responses to Avoid cues than to Win cues while controlling for accuracy, RTs, and response repetition over time

We performed control analyses testing whether the difference in pupil dilation between Go responses to Avoid compared to Win cues could be due to other factors associated with increased pupil dilations, specifically (a) correct vs. incorrect responses, (b) fast vs. slow responses (median split), and (c) response repetitions vs. switches to the alternative response option (with respect to the last encounter of the same cue).

See Table S06 for inferential statistics from mixed-effects linear regression models regressing trial-by-trial pupil dilations onto accuracy, response speed, and response repetition, separately and in interaction with the performed response (Go vs. NoGo). See Table S07 for inferential statistics from generalized additive models testing whether condition differences occurred selectively at particular time points within blocks. Incorrect responses were associated with significantly larger dilations compared to correct responses, an effect that was marginally stronger for NoGo responses (Fig. S09A). Over the time course of blocks, dilations were higher for incorrect NoGo responses than correct NoGo responses on cue repetitions 4 until 13, with no difference between incorrect and correct Go responses (Fig. S09D). Furthermore, slow responses were associated significantly with higher dilations compared to fast responses (Fig. S09B; note that on NoGo trials, no RTs can be observed) throughout blocks (Fig. S09E). Lastly, trials on which participants switched their response with respect to the last encounter of the same cue were associated with significantly higher pupil dilations (Fig. S09C) throughout a block (Fig. S09F), with no interaction with the performed response. In sum, incorrect responses, slower responses, and response switches were associated with stronger pupil dilations.

Both incorrect and slower responses were associated with significantly increased pupil dilations, but also with each other: incorrect responses (to NoGo cues) tended to be slower than correct responses (to Go cues; see Fig. 2E, F in the main text). We thus split trials with Go responses by both accuracy (correct/ incorrect) and response speed (fast/ slow; median split performed separately for correct and incorrect responses for each participant) and tested whether both factors contributed independently to pupil dilations. Slower responses were associated with stronger dilations than faster responses irrespective of accuracy, while accuracy alone had no effect on dilations when controlling for response speed (Fig. S10 and inferential statistics in Tables S06 and S07). Hence, stronger pupil dilations on incorrect compared to correct responses follow from the former being slower than the latter. Note that GAMMs control for any changes in overall response speed or accuracy over time; the difference between fast and slow responses cannot be accounted for by increases in speed and accuracy over time.

Next, we investigated whether higher pupil dilations for Go responses to Avoid cues compared to Win cues were still observed for separate levels of accuracy, response speed (fast/ slow; median split performed separately for Win and Avoid cues for each participant), and response repetition. Dilations were still marginally significantly higher for response to Avoid cues than to Win cues irrespective of accuracy (Fig. S11A, Table S06). Additive models suggested significantly higher dilations for correct Go responses to Avoid than to Win cues on cue repetitions 4–13 as well as higher dilations for incorrect Go responses to Avoid than to Win cues on cue repetitions 6–16 (Fig. S11D, Table S07). Furthermore, while linear regression models suggested significantly higher dilations for slow than fast responses (median split performed separately for Win and Avoid cues), with no significant difference between Avoid and Win cues (Fig. S11B, Table S06), additive models suggested significantly higher dilations for slow responses to Avoid cues than slow responses to Win cues on cue repetitions 4–14, with no such difference for fast responses (Fig. S11E, Table S07). Lastly, while linear regression models indicated significantly higher dilations for response switches than response repetitions, with no differences between Avoid and Win cues (Fig. S11C, Table S06), additive models indicated that significantly higher dilations for response repetitions to Avoid than to Win cues on cue repetitions 3–13 (Fig. S11F, Table S07). For response switches, the pattern of differences was more complicated, with higher dilations for response switches for Avoid cues than for Win cues on the first three repetitions, but the reverse pattern on cue repetitions 6–13.

Taken together, these results suggest that dilations were indeed higher for Go responses to Avoid cues (for which participants had to overcome aversive inhibition) than Go responses to Win cues irrespective of accuracy, suggesting that the observed increase in pupil dilations cannot be attributed to error processing. In fact, seemingly higher dilations to incorrect compared to correct responses are probably attributable to incorrect responses being relatively slower. Moreover, dilations were higher for Go responses to Avoid than to Win cues, but only for slow responses, with no such difference for fast responses. This pattern is in line with our interpretation of pupil dilation reflecting cognitive conflict and heightened physical effort recruitment in order to overcome aversive inhibition, a pattern that should lead to (and should only be observable on trials with) slow responses. In contrast, for fast responses, no such conflict might have occurred, potentially because these responses were made more “impulsively” and without proper processing of the cue or because responses had started to become well learned. Lastly, dilations on Go response repetitions (the large majority of responses) were higher for Avoid cues than Win cues, suggesting that this pattern was not induced by a different pattern of response switches for Avoid than Win cues. Notably, this pattern reversed for response switches. Note however that response switches towards Go were overall rare, and especially so for Win cues (i.e. the green dashed line in Fig. S11F reflects pupil dilations on those trials on which participants had previously performed a NoGo response to a Win cue and then decided to switch towards a Go response, likely because they deemed the previous response to be incorrect—a pattern that occurred very rarely in this task given that participants performed few NoGo responses to Win cues in the first place). In sum, these results are in line with our interpretation of heightened dilations for response to Avoid cues reflecting heightened physical effort recruitment in order to overcome aversive inhibition, a pattern associated with slow responses.

**Figure S09.**
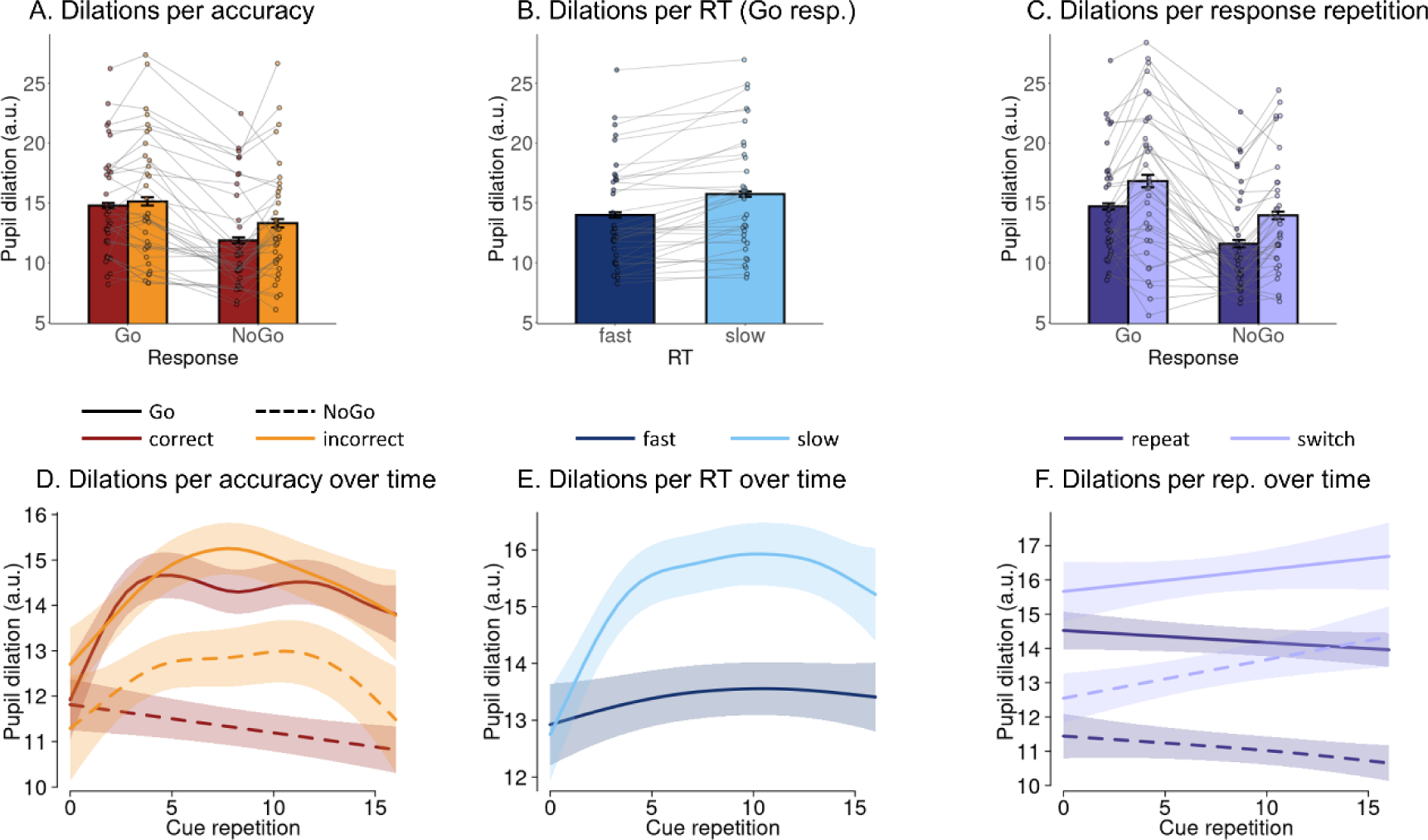
Association of pupil dilation with accuracy, response speed, and response repetition. **A.** Mean pupil dilation per response and accuracy (whiskers are ±SEM across participants, dots indicate individual participants). Dilations are significantly higher for Go than NoGo responses and higher for incorrect than correct responses (an effect that is marginally stronger for NoGo than Go responses). **B**. Mean pupil dilation per response speed (fast/ slow). Dilations are significantly higher for slow compared to fast responses. **C**. Mean pupil dilation per response and response repetition. Dilations are significantly higher for Go than NoGo responses and higher for response switches than response repetitions. **D**. Time course of dilations over cue repetitions (mean ± SE) as predicted from a generalized additive mixed-effects model (GAMM), separated by response and accuracy. Dilations are significantly stronger on trials with Go responses than on trials with NoGo responses throughout blocks. Furthermore, dilations are higher for incorrect than correct NoGo responses on repetitions 4–13. **E**. Time course of dilations over cue repetitions separated by response speed. Dilations are higher for slow compared to fast Go responses throughout blocks. **F**. Time course of dilations over cue repetitions separated by response and response repetition. Dilations are significantly stronger on trials with Go responses than on trials with NoGo responses and for response switches compared to response repetitions throughout blocks.

**Figure S10.**
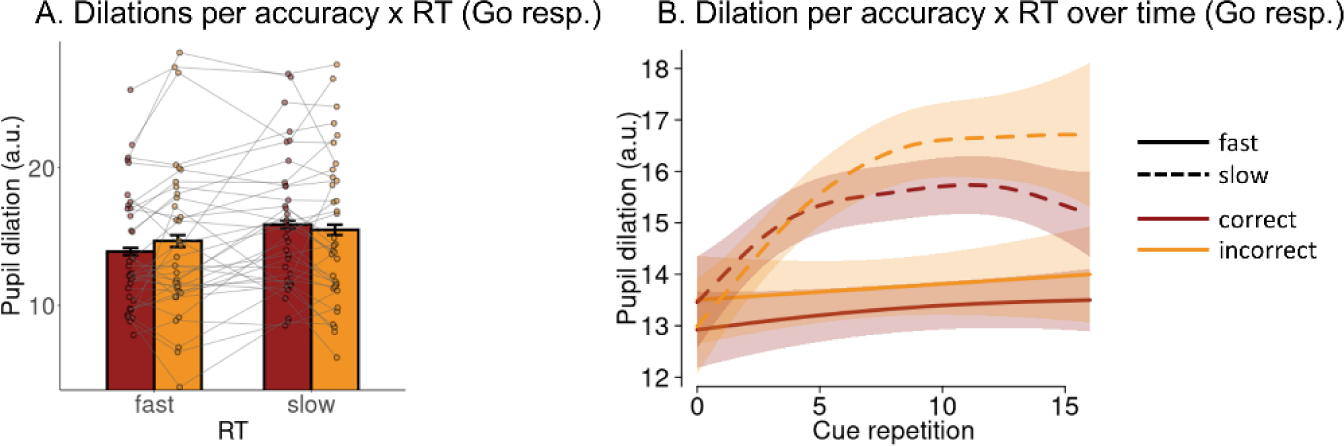
Association of pupil dilation with accuracy and response speed. **A.** Mean pupil dilation split by response speed and accuracy (whiskers are ±SEM across participants, dots indicate individual participants). Dilations are significantly higher on trials with slow responses than on trials with fast responses, with no significant differences between correct and incorrect responses. **B**. Time course of dilations over cue repetitions (mean ± SE) as predicted from a generalized additive mixed-effects model (GAMM), separated by accuracy and response speed. Dilations are significantly higher on trials with slow responses than on trials with fast responses, with no significant differences between correct and incorrect responses.

**Figure S11.**
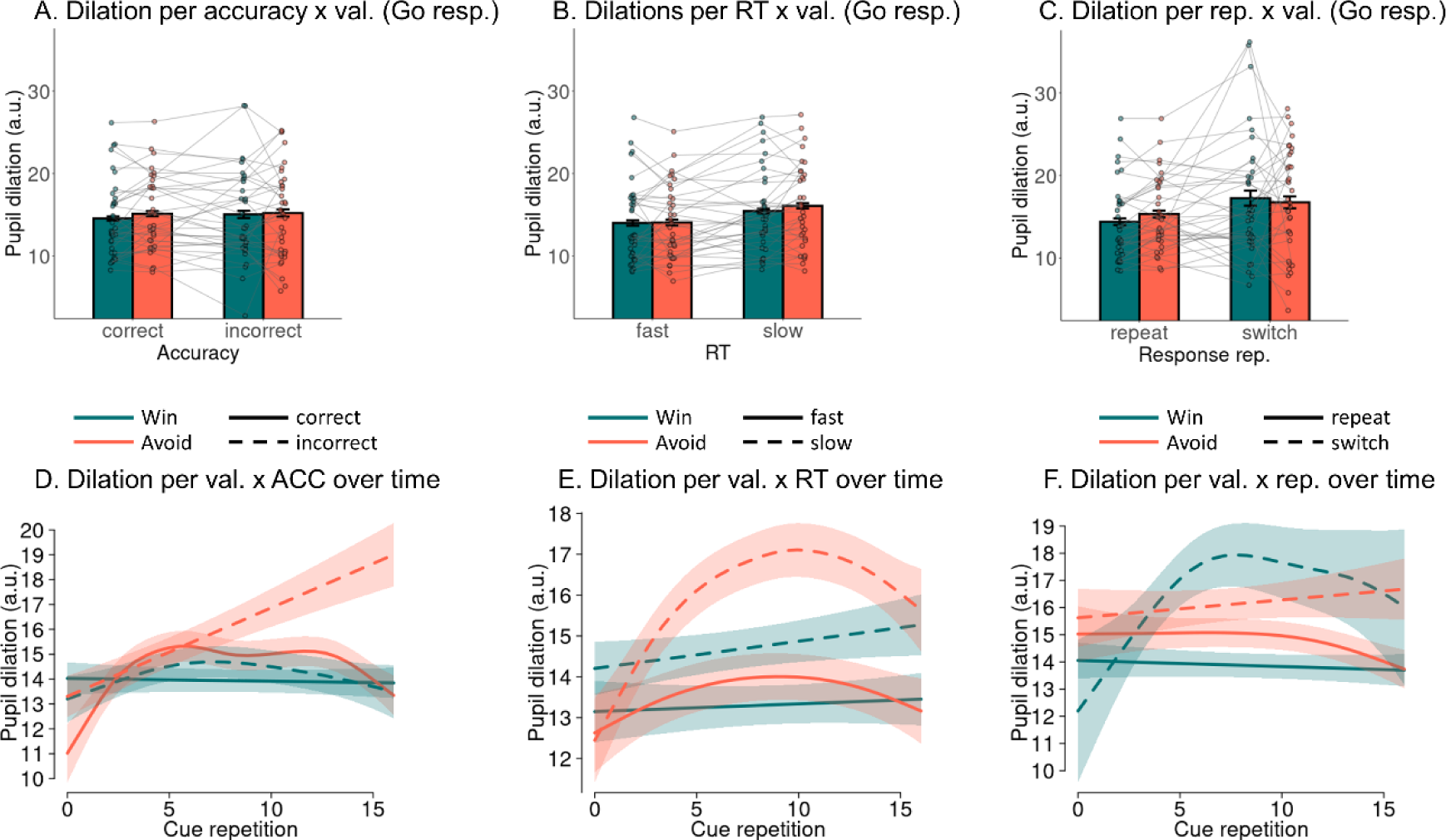
Higher pupil dilation for responses to Win compared to Avoid cues for trials split by accuracy, response speed, and response repetition. **A.** Mean pupil dilation on trials with Go responses per accuracy level per cue valence (whiskers are ±SEM across participants, dots indicate individual participants). Dilations are marginally significantly higher for responses to Avoid than to Win cues. **B**. Mean pupil dilation per response speed (fast/ slow) per cue valence. Dilations are significantly higher for slow compared to fast responses, while the effect of cue valence is not significant. **C**. Mean pupil dilation on trials with Go responses per response repetition per cue valence. Dilations are significantly higher for response repetitions to Avoid than to Win cues, while this effect is reversed for response switches. **D**. Time course of dilations over cue repetitions (mean ± SE) as predicted from a generalized additive mixed-effects model (GAMM), separated by accuracy and cue valence. Dilations are significantly stronger on for correct Go responses to Avoid than to Win cues on cue repetitions 4–13. Moreover, dilations are significantly stronger for incorrect Go responses to Avoid than to Win cues on cue repetitions 6–16. **E**. Time course of dilations over cue repetitions separated by response speed and cue valence. Dilations are significantly higher for slow compared to fast responses throughout blocks. Furthermore, dilations are significantly higher for slow responses to Avoid cues than to Win cues on cue repetitions 4–14, with no such difference for fast responses. **F**. Time course of dilations over cue repetitions separated by response repetition and cue valence. Dilations are significantly higher for response repetitions to Avoid than to Win cues on cue repetitions 3–13. Finally, dilations for response switches for Avoid cues are significantly higher than for Win cues on the first three repetitions, but this pattern reverses later, with stronger dilations for switches for Win cues than for Avoid cues on cue repetitions 6–13.

**Table S06.** Results from mixed-effects linear regression models with trial-by-trial pupil dilation as dependent variable.

| Model ID | Trial subset | DV | IV | b | SE | $\chi^2(1)$ | p |
| --- | --- | --- | --- | --- | --- | --- | --- |
| 1 | All trials | Dilations | Accuracy (correct/ incorrect) | -0.039 | 0.012 | 8.267 | .004 |
|  |  |  | Response (Go/ NoGo) | 0.112 | 0.015 | 33.973 | < .001 |
|  |  |  | Accuracy x Response | 0.026 | 0.012 | 3.532 | .060 |
| 2 | Go responses | Dilations | RTs (fast/ slow) | -0.081 | 0.015 | 21.760 | < .001 |
| 3 | All trials | Dilations | Response repetition (repeat/ switch) | -0.105 | 0.019 | 22.924 | < .001 |
|  |  |  | Response (Go/ NoGo) | 0.139 | 0.019 | 34.249 | < .001 |
|  |  |  | Response repetition x response | 0.008 | 0.015 | 0.320 | .571 |
| 4 | Go responses | Dilations | Accuracy (correct/ incorrect) | -0.007 | 0.016 | 0.224 | .636 |
|  |  |  | RTs (fast/ slow) | -0.073 | 0.017 | 14.429 | < .001 |
|  |  |  | Accuracy x RTs | -0.018 | 0.016 | 1.386 | .239 |
| 5 | Go responses | Dilations | Accuracy (correct/ incorrect) | -0.017 | 0.017 | 1.099 | .338 |
|  |  |  | Valence (Win/ Avoid) | -0.029 | 0.016 | 3.381 | .071 |
|  |  |  | Accuracy x Valence | 0.004 | 0.017 | 0.078 | .730 |
| 6 | Go responses | Dilations | RTs (fast/ slow) | -0.082 | 0.015 | 20.826 | < .001 |
|  |  |  | Valence (Win/ Avoid) | -0.016 | 0.014 | 0.732 | .392 |
|  |  |  | RTs x Valence | 0.016 | 0.014 | 0.812 | .368 |
| 7 | Go responses | Dilations | Response repetition (repeat/ switch) | -0.107 | 0.027 | 12.841 | < .001 |
|  |  |  | Valence (Win/ Avoid) | 0.005 | 0.027 | 0.039 | .844 |
|  |  |  | Response repetition x valence | -0.044 | 0.031 | 2.046 | .153 |

**Table S07.** Results from generalized additive mixed models (GAMMs) with difference smooths between two conditions. The parametric term reflects a linear difference between conditions, while the smooth terms reflects any non-linear difference. Both add up to the total term. The time window of significant condition differences is automatically returned by the model. For the accuracy x RT and RT x valence models, the median split into fast and slow responses is performed separately for correct/ incorrect responses and Win/ Avoid cues for each participant.

| *Model* | **Parametric coefficient**<br>**(Intercept difference)** | **Smooth**<br>**(non-linear differences)** | **Windows of**<br>**significant**<br>**differences** |
| --- | --- | --- | --- |
| **Accuracy (all trials):** |  |  |  |
| *Go correct – Go incorrect* | $t(5.870, 7.707) = 1.657, p = .098$ | $F(1.001, 1.001) = 0.457, p = .499$ | none |
| *NoGo correct – NoGo incorrect* | $t(4.460, 6.596) = 2.671, p = .008$ | $F(3.573, 4.409) = 1.397, p = .198$ | 4 – 13 |
| **RTs (Go responses):** |  |  |  |
| *Fast – slow* | $t(5.710, 7.650) = 7.184, p < .001$ | $F(1.422, 1.702) = 0.751, p = .364$ | 1 – 16 |
| **Repetition (all trials):** |  |  |  |
| *Go repeat – Go switch* | $t(6.054, 7.759) = 5.026, p < .001$ | $F(1.000, 1.000) = 1.792, p = .181$ | 2 – 16 |
| *NoGo repeat – NoGo switch* | $t(4.473, 6.606) = 5.904, p < .001$ | $F(1.000, 1.000) = 1.823, p = .177$ | 1 – 16 |
| **Accuracy x RTs (Go responses):** |  |  |  |
| *Slow Correct – Fast Correct* | $t(5.107, 7.275) = 6.194, p < .001$ | $F(1.000, 1.000) = 0.140, p = .709$ | 0 – 16 |
| *Slow Incorrect – Fast Incorrect* | $t(3.000, 5.191) = 2.879, p = .004$ | $F(1.000, 1.000) = 5.071, p = .025$ | 6 – 16 |
| *Fast Incorrect – Fast Correct* | $t(3.970, 6.536) = 1.616, p = .106$ | $F(1.003, 1.006) = 0.256, p = .617$ | none |
| *Slow Incorrect – Slow Correct* | $t(6.416, 7.818) = 1.304, p = .192$ | $F(1.000, 1.000) = 1.951, p = .163$ | none |
| **Accuracy x Valence (Go responses):** |  |  |  |
| *Correct Avoid – correct Win* | $t(5.182, 7.313) = 2.244, p = .025$ | $F(4.479, 5.456) = 3.839, p = .001$ | 4 – 13 |
| *Incorrect Avoid – incorrect Win* | $t(3.000, 5.253) = 2.159, p = .031$ | $F(1.000, 1.000) = 2.573, p = .109$ | 6 – 16 |
| **RTs x Valence (Go responses):** |  |  |  |
| *Fast Avoid – fast Win* | $t(4.582, 6.825) = 0.958, p = .338$ | $F(1.798, 2.176) = 0.408, p = .758$ | none |
| *Slow Avoid – slow Win* | $t(5.974, 7.799) = 3.222, p = .001$ | $F(2.384, 2.936) = 2.409, p = .065$ | 4 – 14 |
| **Repetition x Valence (Go responses):** |  |  |  |
| *Repeat Avoid – repeat Win* | $t(5.225, 7.400) = 3.246, p = .001$ | $F(1.856, 2.278) = 0.869, p = .353$ | 3 – 13 |
| *Switch Avoid – switch Win* | $t(5.710, 7.650) = 7.184, p < .001$ | $F(1.422, 1.702) = 0.751, p = .364$ | 0 – 2, 6 – 13 |

## Supplementary Material S08: Association of pupil baseline with accuracy, RTs, and response repetition over time

Beyond task-evoked trial-by-trial pupil dilations, past literature has also investigated pre-stimulus baseline pupil diameter as a potential readout of noradrenergic activity (Aston-Jones & Cohen, 2005; Eldar, Cohen, & Niv, 2013; Gilzenrat, Nieuwenhuis, Jepma, & Cohen, 2010). One the one hand, pupil baseline and task-evoked pupil dilation tend to be negatively correlated since high baseline leave less dynamic range for further dilations. In this sense, both measures could potentially capture similar phenomena and are partly redundant. However, on the other hand, pupil dilations are corrected for the immediately preceding pre-stimulus baseline and thus cannot reflect more “tonic” changes in pupil diameter on time scales longer than a single trial. In fact, pupil baseline itself tends to strongly decrease over the time course of an experiment (Muller, Mars, Behrens, & O’Reilly, 2019), likely reflecting decreases in arousal. These slower changes might reflect processes orthogonal to the trial-by-trial pupil dilations. Given that baselines are measured before cue onset, they cannot reflect the (randomized) task conditions (required action, valence, and arousal manipulation). Nonetheless, the process they reflect could still impact (or at least predict) task performance (responses, accuracy, and RTs).

While on the one hand, baseline pupil diameter could lead additional insights into cognitive processes beyond pupil dilation, on the other hand, caution is warranted given that possibility of spurious associations driven by time. When baseline pupil diameter decreases over time, any other variable that also changes on a similar time scale might be spuriously correlated with pupil diameter. Here, we used mixed-effects linear regression and generalized additive mixed effects models to test for effects of the baseline pupil diameter on responses, accuracy, and RTs (fast vs. slow, median split), controlling for potential linear and non-linear effects of time (cue repetition, 1–16).

See Table S08 for inferential statistics from mixed-effects linear regressions. See Fig. S12A-C for baselines per condition averaged over trials. When ignoring time, higher baseline pupil diameter was associated with a significantly higher propensity of Go responses, incorrect responses, and slower responses (see Table S08; Fig. S12A-C). The associations with accuracy and RTs disappeared when controlling for a linear effect of cue repetition (see Table S08). Most notably, additive models suggested that baseline pupil diameter strongly decreased over time (Fig. S12D-F), with no significant difference between Go and NoGo responses, correct and incorrect responses, and only a minor (albeit significant) difference between fast and slow responses (Table S09; Fig. S12D-F) on the first eight cue repetitions, which was in fact of opposite sign (i.e., higher baselines before fast responses) to the results from the mixed-effect linear regression model (Fig. S12C). Thus, indeed, spurious associations between baseline pupil diameter and other variables arise through both changing over time, with participants showing less Go responses, less incorrect responses, and faster responses as they progress through a task block. In sum, there was strong evidence for baseline pupil diameter decreasing over the time course of a block, but no strong evidence for baseline pupil diameter affecting subsequent responses.

See Fig. S13A-C for the pupil dilation time course within a trial split by response and cue-valence when no baseline-correction is applied. Go responses to Avoid cues were associated with considerably stronger pupil dilations than Go responses to Win cues, However, this was partly driven by pre-existing baseline differences between those two trial types. Since baselines decreased with time, higher baselines on trials with Go responses to Avoid cues compared to those with Go responses to Win cues could potentially be explained by the former occurring relatively earlier within blocks (when baselines were still higher) than the latter. However, the opposite was the case: as participants learned the task, they showed more Go responses to Avoid cues with time, and the ratio between Go responses to Win and Avoid cues approached 50:50 with time. Hence, the overall decay in baseline cannot explain baseline differences between these two trial types. In fact, baseline differences were even stronger in the second half of blocks (Fig. S13C) compared to the first half (Fig. 13B), i.e. they prevailed and became even stronger as the ratio of both trial types approached 50:50. A generalized additive model corroborated that pupil baselines were significantly higher on trials with Go responses to Avoid cues compared to trials with Gon responses to Win cues in the second half of blocks (Fig. S13D, E; Table S09). In sum, Go responses to Avoid cues were not only associated with higher pupil dilations, but also higher pupil baselines, suggesting that pre-existing differences arousal before cue onset might have contributed to the mobilization of effort and invigoration of Go responses against aversive Pavlovian biases.

**Figure S12.**
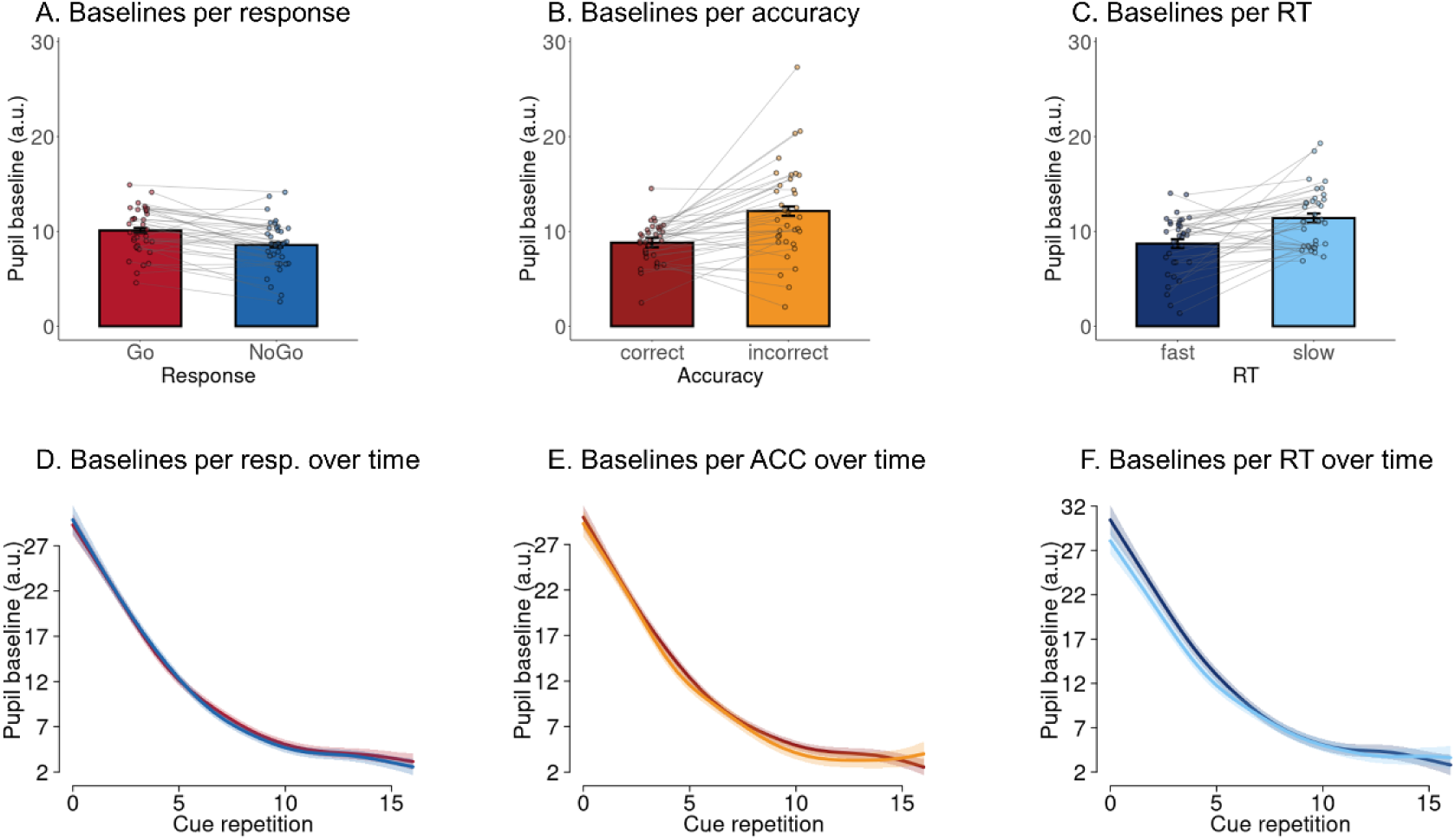
Relationship of pre-trial baseline pupil diameter with responses, accuracy, and RTs. **A.** Pupil pre-trial baseline split by the response made on the trial (whiskers are ± SEM across participants, dots indicate individual participants). Considering trials irrespective of their temporal position within a block, baseline pupil diameter is significantly higher before trials with Go responses than trials with NoGo responses. **B**. Pupil baseline split by the speed of the response made on the following trial (only trials with Go responses). Considering trials irrespective of their temporal position within a block, baseline pupil diameter is significantly higher before trials with incorrect responses than trials with correct responses. **C**. Pupil baseline split by the accuracy of the response made on the following trial. Considering trials irrespective of their temporal position within a block, baseline pupil diameter is significantly higher before trials with slow responses than trials with fast responses. **D**. Time course of baseline pupil diameter over cue repetitions (mean ± SE) as predicted by a generalized additive mixed-effects model (GAMM), separated by responses. There is no significant difference between trials with Go and NoGo responses. **E**. Time course of baseline pupil diameter over cue repetitions as predicted by a generalized additive mixed-effects model (GAMM), separated by accuracy. There is no significant difference between trials with Go and NoGo responses. **F.** Time course of baseline pupil diameter over cue repetitions (mean ± SE) as predicted by a generalized additive mixed-effects model (GAMM), separated by response speed (fast/ slow; median split). For the first eight cue repetitions, baseline pupil diameter is higher before fast compared to slow responses.

**Figure S13.**
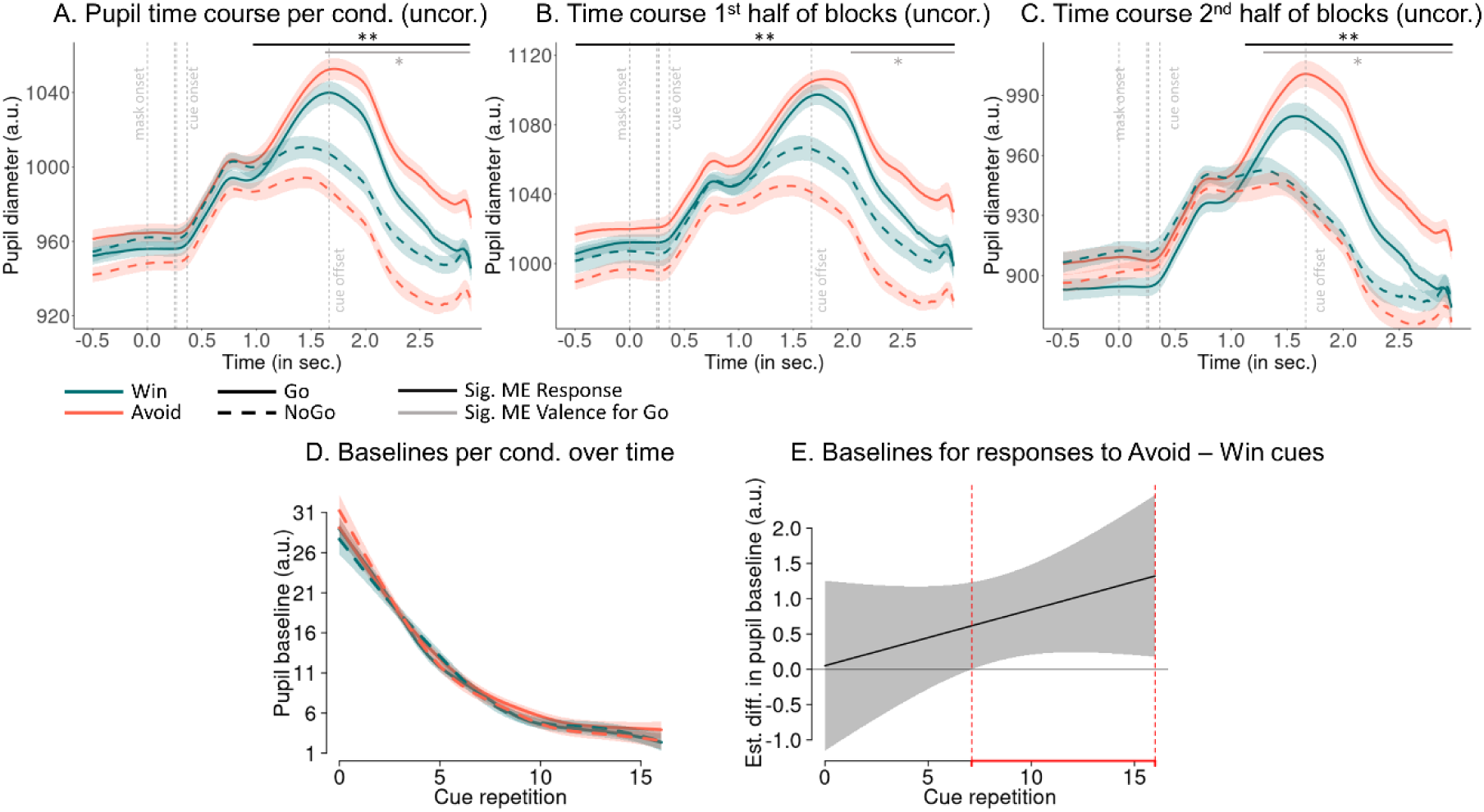
Pupil time course within a trial per response per cue valence without baseline correction (mean ± SEM across participants). **A.** Pupil time course split by cue valence and response made (whiskers are ± SEM across participants, dots indicate individual participants). Vertical dashed lines indicate the onset of the forward mask (at 0 ms), the prime (at 250 ms), the backwards mask (at 266 ms), the cue onset (at 366 ms), and the cue offset (at 1666 ms). The pupil dilates significantly more strongly on trials with Go responses than on trials with NoGo responses (cluster above threshold: 917–2,966 ms; *p* < .001; longer black horizontal line). Furthermore, within this time window, the pupil dilates significantly more strongly and sustainedly for responses to Avoid than to Win cues (cluster above threshold: 1,545–2,966 ms; *p* = .011; shorter black horizontal line). Note however that pre-cue pupil baselines are already higher for Go responses to Avoid cues than Go responses to Win cues. **B.** When repeating this analysis for only the first half of trials within a block, the pupil is wider on trials with Go responses than on trials with NoGo responses throughout the entire time window (cluster above threshold: -1,000–2,966 ms; *p* < .001; longer black horizontal line) and, within this time window, wider for Go responses to Avoid than to Win cues (cluster above threshold: 2,038–2,966 ms; *p* = .049; short black horizontal line). **(C)** In the second half of trials, the pupil is wider on trials with Go responses than on trials with NoGo responses in a more restricted time window (cluster above threshold: 1,137–2,966 ms; *p* < .001) and, within this time window, wider for Go responses to Avoid than to Win cues (cluster above threshold: 1,262–2,966 ms; *p* < .001). The fact that the differences in pupil diameter for Go responses to Avoid cues compared to responses to Win cues gets larger with time suggests that people learn to mobilize effort to invigorate Go responses against the Pavlovian bias (aversive inhibition) present on trials with Avoid cues. **D.** Time course of pupil baselines over cue repetitions (mean ± SE) as predicted from a generalized additive mixed-effects model (GAMM), separated by response and cue valence. Baselines are significantly stronger on trials with Go responses than on trials with Go responses to Avoid cues than trials with Go responses to Win cues from cue repetition 7 to 16, putatively reflecting that pre-cue fluctuations in arousal contribute to the invigoration of Go response against aversive Pavlovian biases. **E**. Difference line between baselines on trials with responses to Avoid cues minus Win cues. Areas highlighted in red indicate time windows with significant differences.

**Table S08.**
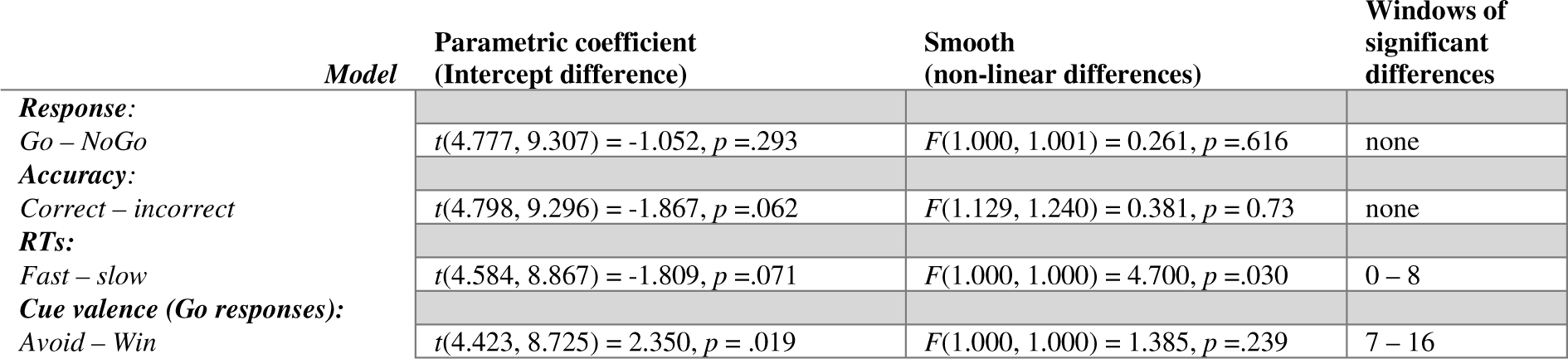
Results from mixed-effects linear regression models with trial-by-trial baseline pupil diameter as dependent variable.

**Table S09.**
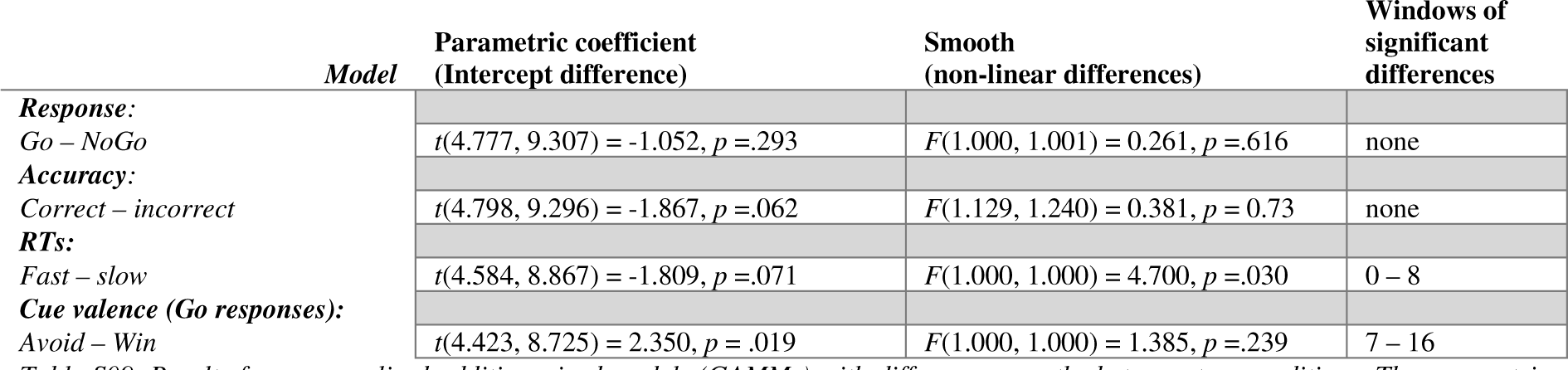
Results from generalized additive mixed models (GAMMs) with difference smooths between two conditions. The parametric term reflects a linear difference between conditions, while the smooth terms reflects any non-linear difference. Both add up to the total term. The time window of significant condition differences is automatically returned by the model.

## Supplementary Material S09: Outcome-locked pupil dilation

Apart from cue- (or masked-) locked pupil dilation, we also investigated outcome-locked pupil dilation (epoched from -1000 ms before until 2000 ms after outcome onset) as a function of the obtained outcome and the previously made response.

See Table S10 and Fig. S14 for results from mixed-effects linear regression models as well as post-hoc *z*-tests contrasting conditions against each other. Pupil dilations were significantly stronger on trials with punishments compared to trials with rewards or neutral outcomes, while trials with rewards and neutral outcomes were not significantly different from each other. Dilations were not different between trials on which neutral outcomes signaled the absence of rewards compared to trials on which they signaled the absence of punishments.

When analyzing dilations as a function of both the obtained outcome and the previously made response, we observed main effects of outcome and response, while the interaction between them was not significant (Table S10). Pupil dilations were higher after NoGo responses compared to Go responses (Fig. S15A). However, inspection of the raw pupil time course within a trial revealed that this difference was an artifact of baseline correction: raw pupil time courses tended to be higher after Go compared to NoGo responses (for trials with punishment and neutral outcomes; Fig. S15C), leaving less dynamic range for further increases on Go compared to NoGo trials and thus leading to lower (baseline-corrected) pupil dilations on Go compared to NoGo trials (Fig. S15B).

In sum, the pupil dilated more strongly in response to punishments compared to rewards or neutral outcomes.

**Table S10.** Results from mixed-effects linear regression models with outcome-locked trial-by-trial pupil dilation as dependent variable. Differences between any conditions were first tested with χ^2^ tests and then followed up with z-tests testing two conditions against each other. P-values for the follow-up z-tests are corrected for multiple comparisons using the Tukey method.

| *Model ID* | *DV* | *IV* | $\chi^2(I)$ | *z* | *p* |
| --- | --- | --- | --- | --- | --- |
| **1** | *Pupil dilation* | *Outcome valence (positive/ negative)* | 13.439 |  | < .001 |
| **2** | *Pupil dilation* | *Outcome displayed (reward/ neutral/ punishment)* | 27.237 |  | < .001 |
|  |  | *Punishment – neutral* |  | 6.351 | < .001 |
|  |  | *Punishment – reward* |  | 5.473 | < .001 |
|  |  | *Neutral – reward* |  | 1.093 | .519 |
| **3** | *Pupil dilation* | *Outcome interpreted (rew./ no rew./ no pun./ pun.)* | 31.251 |  | < .001 |
|  |  | *Punished vs. not punished* |  | 5.591 | < .001 |
|  |  | *Punished vs. not rewarded* |  | 6.996 | < .001 |
|  |  | *Punished vs. rewarded* |  | 5.457 | < .001 |
|  |  | *Not punished vs. not rewarded* |  | 1.586 | .387 |
|  |  | *Not punished vs. rewarded* |  | 0.321 | .989 |
|  |  | *Not rewarded vs. rewarded* |  | 2.021 | .180 |
| **4** | *Pupil dilation* | *Outcome displayed (reward/ neutral/ punishment)* | 25.704 |  | < .001 |
|  |  | *Response (Go/ NoGo)* | 19.116 |  | < .001 |
|  |  | *Outcome displayed x response* | 1.306 |  | .521 |

**Figure S14.**
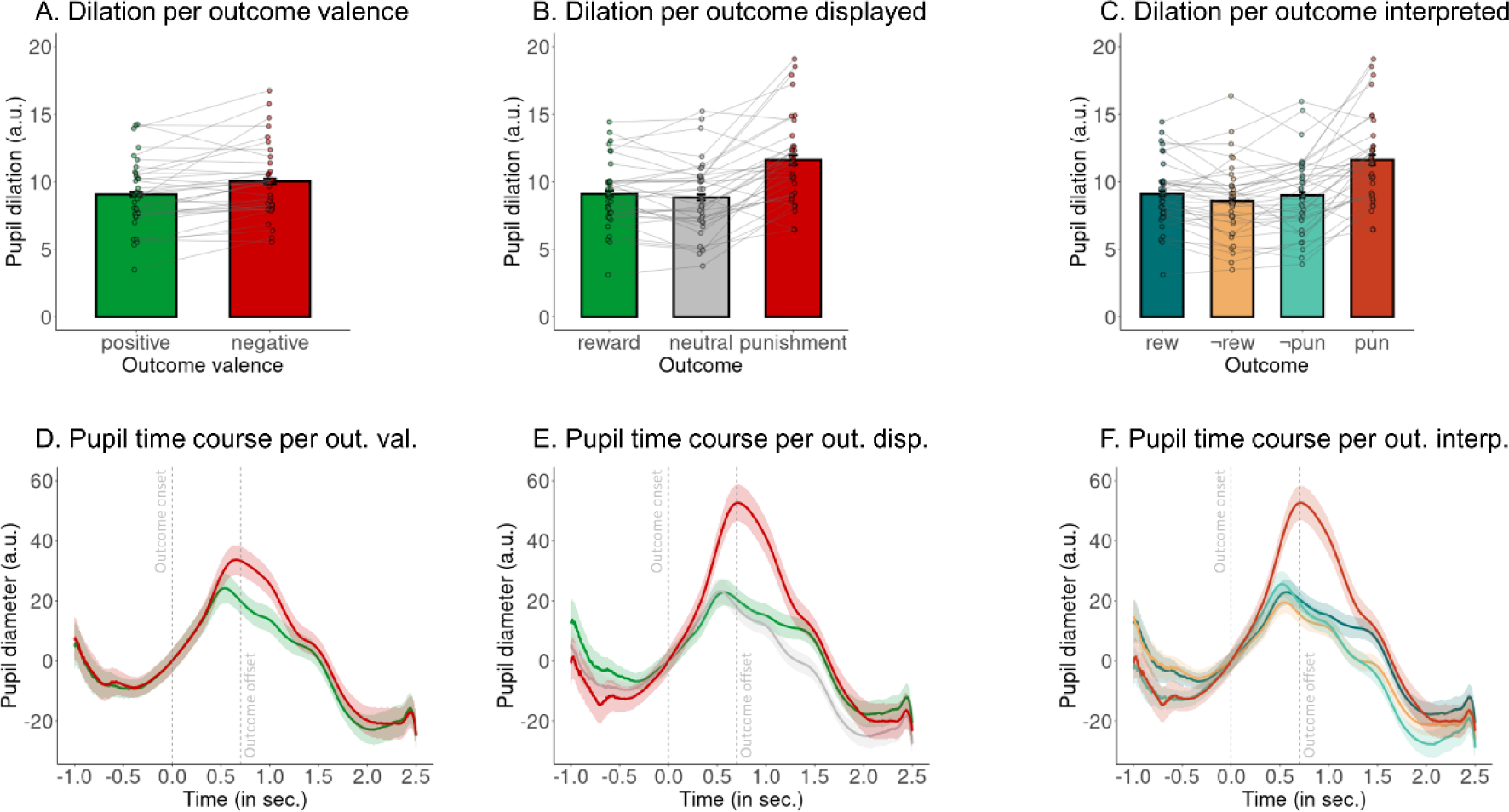
Effect of outcomes on outcome-locked pupil dilation. Pupil dilation as a function of outcome valence (**A**), the displayed outcome (**B**) or the outcome interpreted (with neutral outcomes recognized as signaling the absence of a reward/ punishment, **C**; whiskers are ± SEM across participants, dots indicate individual participants). The pupil dilates more strongly on trials with punishments compared to rewards or neutral outcomes. (**D-F**) Pupil time course within a trial separately for the different outcome conditions (mean ± SEM across participants; baseline-corrected). Vertical dashed line represent the onset (at 0 ms) and offset (at 700 ms) of outcomes.

**Figure S15.**
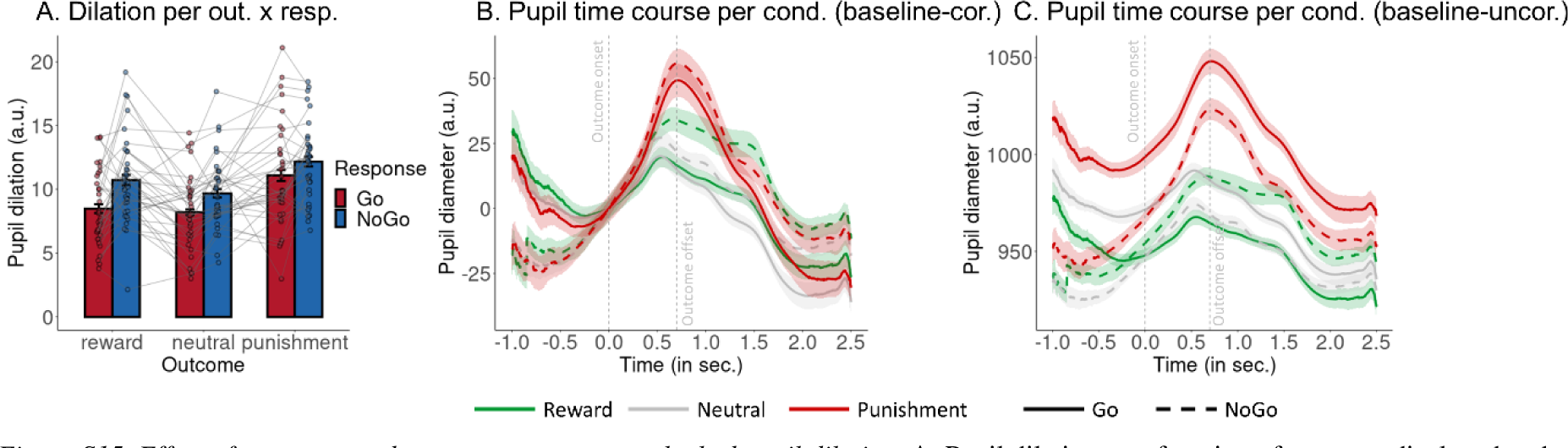
Effect of outcomes and responses on outcome-locked pupil dilation. **A.** Pupil dilation as a function of outcome displayed and the response performed on the same trial manipulation (whiskers are ±SEM across participants, dots indicate individual participants). When applying baseline-correction for differences in the time window of 500 ms before outcome onset, dilations are significantly higher on trials with punishments compared to trials with rewards or neutral outcomes and higher on trials with NoGo than trials with Go responses. **B**. Pupil time course within a trial separately per outcome and response condition (mean ± SEM across participants; baseline-corrected). It appears that for trials with rewards and neutral outcomes, pupil dilations are higher after NoGo than Go responses. Vertical dashed line represent the onset (at 0 ms) and offset (at 700 ms) of outcomes. **C**. Same as panel B, but not baseline corrected. It becomes clear that the pupil time course is higher after Go compared to NoGo responses, leaving less room for further increase on trials with Go compared to NoGo responses, explaining while the baseline-corrected dilations tends to be smaller after Go than NoGo responses.

## References

1. Aarts, E., Verhage, M., Veenvliet, J. V., Dolan, C. V., & van der Sluis, S. (2014). A solution to dependency: Using multilevel analysis to accommodate nested data. Nature Neuroscience, 17(4), 491–496. doi: 10.1038/nn.3648

2. Algermissen, J., Bijleveld, E., Jostmann, N. B., & Holland, R. W. (2019). Explore or reset? Pupil diameter transiently increases in self-chosen switches between cognitive labor and leisure in either direction. Cognitive, Affective, & Behavioral Neuroscience, 379214. doi: 10.3758/s13415-019-00727-x

3. Algermissen, J., Swart, J. C., Scheeringa, R., Cools, R., & den Ouden, H. E. M. (2022). Striatal BOLD and midfrontal theta power express motivation for action. Cerebral Cortex, 32(14), 2924–2942. doi: 10.1093/cercor/bhab391

4. Allen, M., Frank, D., Schwarzkopf, D. S., Fardo, F., Winston, J. S., Hauser, T. U., & Rees, G. (2016). Unexpected arousal modulates the influence of sensory noise on confidence. eLife, 5, 1–17. doi: 10.7554/eLife.18103

5. Amita, H., & Hikosaka, O. (2019). Indirect pathway from caudate tail mediates rejection of bad objects in periphery. Science Advances, 5(8), eaaw9297. doi: 10.1126/sciadv.aaw9297

6. Baayen, H., Vasishth, S., Kliegl, R., & Bates, D. (2017). The cave of shadows: Addressing the human factor with generalized additive mixed models. Journal of Memory and Language, 94(5), 206–234. doi: 10.1016/j.jml.2016.11.006

7. Barr, D. J., Levy, R., Scheepers, C., & Tily, H. J. (2013). Random effects structure for confirmatory hypothesis testing: Keep it maximal. Journal of Memory and Language, 68(3), 255–278. doi: 10.1016/j.jml.2012.11.001

8. Bates, D., Mächler, M., Bolker, B., & Walker, S. (2015). Fitting linear mixed-effects models using lme4. Journal of Statistical Software, 67, 1–48. doi: 10.18637/jss.v067.i01

9. Beatty, J. (1982). Task-evoked pupillary responses, processing load, and the structure of processing resources. Psychological Bulletin, 91(2), 276–292. doi: 10.1037/0033-2909.91.2.276

10. Berke, J. D. (2018). What does dopamine mean? Nature Neuroscience, 21(6), 787–793. doi: 10.1038/s41593-018-0152-y

11. Bijleveld, E., Custers, R., & Aarts, H. (2009). The unconscious eye opener: Pupil dilation reveals strategic recruitment of resources upon presentation of subliminal reward cues. Psychological Science, 20(11), 1313–1315. doi: 10.1111/j.1467-9280.2009.02443.x

12. Blanchard, D. C. (2017). Translating dynamic defense patterns from rodents to people. Neuroscience & Biobehavioral Reviews, 76, 22–28. doi: 10.1016/j.neubiorev.2016.11.001

13. Bornert, P., & Bouret, S. (2021). Locus coeruleus neurons encode the subjective difficulty of triggering and executing actions. PLOS Biology, 19(12), e3001487. doi: 10.1371/journal.pbio.3001487

14. Cavanagh, J. F., Eisenberg, I., Guitart-Masip, M., Huys, Q. J. M., & Frank, M. J. (2013). Frontal theta overrides Pavlovian learning biases. Journal of Neuroscience, 33(19), 8541–8548. doi: 10.1523/JNEUROSCI.5754-12.2013

15. Chen, K., Schlagenhauf, F., Sebold, M., Kuitunen-Paul, S., Chen, H., Huys, Q. J. M., … Garbusow, M. (2023). The association of non-drug-related Pavlovian-to-instrumental transfer effect in nucleus accumbens with relapse in alcohol dependence: A replication. Biological Psychiatry, 93(6), 558–565. doi: 10.1016/j.biopsych.2022.09.017

16. Cyders, M. A., Littlefield, A. K., Coffey, S., & Karyadi, K. A. (2014). Examination of a short English version of the UPPS-P Impulsive Behavior Scale. Addictive Behaviors, 39(9), 1372–1376. doi: 10.1016/j.addbeh.2014.02.013

17. da Silva Castanheira, K., LoParco, M., & Otto, A. R. (2020). Task-evoked pupillary responses track effort exertion: Evidence from task-switching. Cognitive, Affective, & Behavioral Neuroscience, 1–15. doi: 10.3758/s13415-020-00843-z

18. D’Ascenzo, S., Iani, C., Guidotti, R., Laeng, B., & Rubichi, S. (2016). Practice-induced and sequential modulations in the Simon task: Evidence from pupil dilation. International Journal of Psychophysiology, 110, 187–193. doi: 10.1016/j.ijpsycho.2016.08.002

19. Daw, N. D., Niv, Y., & Dayan, P. (2005). Uncertainty-based competition between prefrontal and dorsolateral striatal systems for behavioral control. Nature Neuroscience, 8(12), 1704–1711. doi: 10.1038/nn1560

20. Dayan, P. (2014). Rationalizable irrationalities of choice. Topics in Cognitive Science, 6(2), 204–228. doi: 10.1111/tops.12082

21. Dayan, P., Niv, Y., Seymour, B., & Daw, N. (2006). The misbehavior of value and the discipline of the will. Neural Networks, 19(8), 1153–1160. doi: 10.1016/j.neunet.2006.03.002

22. de Gee, J. W., Colizoli, O., Kloosterman, N. A., Knapen, T., Nieuwenhuis, S., & Donner, T. H. (2017). Dynamic modulation of decision biases by brainstem arousal systems. eLife, 6(Lc), 1–36. doi: 10.7554/eLife.23232

23. de Gee, J. W., Correa, C. M. C., Weaver, M., Donner, T. H., & van Gaal, S. (2021). Pupil dilation and the slow wave ERP reflect surprise about choice outcome resulting from intrinsic variability in decision confidence. Cerebral Cortex, 1–14. doi: 10.1093/cercor/bhab032

24. Dippel, G., Mückschel, M., Ziemssen, T., & Beste, C. (2017). Demands on response inhibition processes determine modulations of theta band activity in superior frontal areas and correlations with pupillometry – Implications for the norepinephrine system during inhibitory control. NeuroImage, 157(June), 575–585. doi: 10.1016/j.neuroimage.2017.06.037

25. Dixon, M. L., & Christoff, K. (2012). The decision to engage cognitive control is driven by expected reward-value: Neural and behavioral evidence. PLoS ONE, 7(12), e51637. doi: 10.1371/journal.pone.0051637

26. Dorfman, H. M., & Gershman, S. J. (2019). Controllability governs the balance between Pavlovian and instrumental action selection. Nature Communications, 10(1), 5826. doi: 10.1038/s41467-019-13737-7

27. Fiedler, S., Schulte-Mecklenbeck, M., Renkewitz, F., & Orquin, J. L. (2020). Guideline for reporting standards of eye-tracking research in decision sciences. PsyArXiv.

28. Frank, M. J. (2006). Hold your horses: A dynamic computational role for the subthalamic nucleus in decision making. Neural Networks, 19(8), 1120–1136. doi: 10.1016/j.neunet.2006.03.006

29. Grogan, J. P., Sandhu, T. R., Hu, M. T., & Manohar, S. G. (2020). Dopamine promotes instrumental motivation, but reduces reward-related vigour. eLife, 9, e58321. doi: 10.7554/eLife.58321

30. Guitart-Masip, M., Duzel, E., Dolan, R., & Dayan, P. (2014). Action versus valence in decision making. Trends in Cognitive Sciences, 18(4), 194–202. doi: 10.1016/j.tics.2014.01.003

31. Guitart-Masip, M., Huys, Q. J. M., Fuentemilla, L., Dayan, P., Duzel, E., & Dolan, R. J. (2012). Go and no-go learning in reward and punishment: Interactions between affect and effect. NeuroImage, 62(1), 154–166. doi: 10.1016/j.neuroimage.2012.04.024

32. Hamid, A. A. (2021). Dopaminergic specializations for flexible behavioral control: Linking levels of analysis and functional architectures. Current Opinion in Behavioral Sciences, 41, 175–184. doi: 10.1016/j.cobeha.2021.07.005

33. Hamid, A. A., Frank, M. J., & Moore, C. I. (2021). Wave-like dopamine dynamics as a mechanism for spatiotemporal credit assignment. Cell, 184(10), 2733–2749.e16. doi: 10.1016/j.cell.2021.03.046

34. Hamid, A. A., Pettibone, J. R., Mabrouk, O. S., Hetrick, V. L., Schmidt, R., Vander Weele, C. M., … Berke, J. D. (2016). Mesolimbic dopamine signals the value of work. Nature Neuroscience, 19(1), 117–126. doi: 10.1038/nn.4173

35. Hashemi, M. M., Gladwin, T. E., de Valk, N. M., Zhang, W., Kaldewaij, R., van Ast, V., … Roelofs, K. (2019). Neural dynamics of shooting decisions and the switch from freeze to fight. Scientific Reports, 9(1), 4240. doi: 10.1038/s41598-019-40917-8

36. Hess, E. H., & Polt, J. M. (1964). Pupil size in relation to mental activity during simple problem-solving. Science, 143(3611), 1190–1192. doi: 10.1126/science.143.3611.1190

37. Hoeks, B., & Levelt, W. J. M. (1993). Pupillary dilation as a measure of attention: A quantitative system analysis. Behavior Research Methods, Instruments, & Computers, 25(1), 16–26. doi: 10.3758/BF03204445

38. Huys, Q. J. M., Gölzer, M., Friedel, E., Heinz, A., Cools, R., Dayan, P., & Dolan, R. J. (2016). The specificity of Pavlovian regulation is associated with recovery from depression. Psychological Medicine, 46(05), 1027–1035. doi: 10.1017/S0033291715002597

39. Joshi, S., & Gold, J. I. (2019). Pupil size as a window on neural substrates of cognition. Trends in Cognitive Sciences, (December), 1–24. doi: 10.31234/osf.io/dvsme

40. Joshi, S., Li, Y., Kalwani, R. M., & Gold, J. I. (2016). Relationships between pupil diameter and neuronal activity in the locus coeruleus, colliculi, and cingulate cortex. Neuron, 89(1), 221–234. doi: 10.1016/j.neuron.2015.11.028

41. Kahneman, D. (1973). Attention and effort. Englewood Cliffs, NJ: Prentice Hall.

42. Kahneman, D. (2011). Thinking, fast and slow. New York, NY: Farrar, Strauss, and Giroux.

43. Kawagoe, R., Takikawa, Y., & Hikosaka, O. (1998). Expectation of reward modulates cognitive signals in the basal ganglia. Nature Neuroscience, 1(5), 411–416. doi: 10.1038/1625

44. Keramati, M., Dezfouli, A., & Piray, P. (2011). Speed/accuracy trade-off between the habitual and the goal-directed processes. PLoS Computational Biology, 7(5), e1002055. doi: 10.1371/journal.pcbi.1002055

45. Kim, H. F., Amita, H., & Hikosaka, O. (2017). Indirect pathway of caudal basal ganglia for rejection of valueless visual objects. Neuron, 94(4), 920–930.e3. doi: 10.1016/j.neuron.2017.04.033

46. Klaassen, F. H., Held, L., Figner, B., O’Reilly, J. X., Klumpers, F., de Voogd, L. D., & Roelofs, K. (2021). Defensive freezing and its relation to approach–avoidance decision-making under threat. Scientific Reports, 11(1), 12030. doi: 10.1038/s41598-021-90968-z

47. Kurniawan, I. T., Grueschow, M., & Ruff, C. C. (2021). Anticipatory energization revealed by pupil and brain activity guides human effort-based decision making. Journal of Neuroscience, 41(29), 6328–6342. doi: 10.1523/JNEUROSCI.3027-20.2021

48. Lin, H., Saunders, B., Hutcherson, C. A., & Inzlicht, M. (2018). Midfrontal theta and pupil dilation parametrically track subjective conflict (but also surprise) during intertemporal choice. NeuroImage, 172(August 2017), 838–852. doi: 10.1016/j.neuroimage.2017.10.055

49. Lloyd, B., de Voogd, L. D., Mäki-Marttunen, V., & Nieuwenhuis, S. (2023). Pupil size reflects activation of subcortical ascending arousal system nuclei during rest. eLife, 12, e84822. doi: 10.7554/eLife.84822

50. Loewenstein, G., & O’Donoghue, T. (2004, May 4). *Animal spirits: Affective and deliberative processes in economic behavior* [SSRN Scholarly Paper]. Rochester, NY. doi: 10.2139/ssrn.539843

51. Lundqvist, D., Flykt, A., & Öhman, A. (1998). Karolinska directed emotional faces [Database of standardized facial images] (pp. 171–176). Stockholm, Sweden: CD ROM from Department of Clinical Neuroscience, Psychology section, Karolinska Institutet.

52. Ly, V., Huys, Q. J. M., Stins, J. F., Roelofs, K., & Cools, R. (2014). Individual differences in bodily freezing predict emotional biases in decision making. Frontiers in Behavioral Neuroscience, 8. Retrieved from https://www.frontiersin.org/articles/10.3389/fnbeh.2014.00237

53. Mahlberg, J., Seabrooke, T., Weidemann, G., Hogarth, L., Mitchell, C. J., & Moustafa, A. A. (2021). Human appetitive Pavlovian-to-instrumental transfer: A goal-directed account. Psychological Research, 85(2), 449–463. doi: 10.1007/s00426-019-01266-3

54. Manohar, S. G., Chong, T. T.-J., Apps, M. A. J., Batla, A., Stamelou, M., Jarman, P. R., … Husain, M. (2015). Reward pays the cost of noise reduction in motor and cognitive control. Current Biology, 25(13), 1707–1716. doi: 10.1016/j.cub.2015.05.038

55. Maris, E., & Oostenveld, R. (2007). Nonparametric statistical testing of EEG- and MEG-data. Journal of Neuroscience Methods, 164(1), 177–190. doi: 10.1016/j.jneumeth.2007.03.024

56. Megemont, M., McBurney-Lin, J., & Yang, H. (2022). Pupil diameter is not an accurate real-time readout of locus coeruleus activity. eLife, 11, 1–17. doi: 10.7554/eLife.70510

57. Merscher, A.-S., & Gamer, M. (2024). Fear lies in the eyes of the beholder—Robust evidence for reduced gaze dispersion upon avoidable threat. Psychophysiology, 61(1), e14421. doi: 10.1111/psyp.14421

58. Merscher, A.-S., Tovote, P., Pauli, P., & Gamer, M. (2022). Centralized gaze as an adaptive component of defensive states in humans. Proceedings of the Royal Society B: Biological Sciences, 289(1975), 20220405. doi: 10.1098/rspb.2022.0405

59. Metcalfe, J., & Mischel, W. (1999). A hot/cool-system analysis of delay of gratification: Dynamics of willpower. Psychological Review, 106(1), 3–19. doi: 10.1037/0033-295X.106.1.3

60. Milli, S., Lieder, F., & Griffiths, T. L. (2021). A rational reinterpretation of dual-process theories. Cognition, 217, 104881. doi: 10.1016/j.cognition.2021.104881

61. Mkrtchian, A., Aylward, J., Dayan, P., Roiser, J. P., & Robinson, O. J. (2017). Modeling avoidance in mood and anxiety disorders using reinforcement learning. Biological Psychiatry, 82(7), 532–539. doi: 10.1016/j.biopsych.2017.01.017

62. Mohebi, A., Pettibone, J. R., Hamid, A. A., Wong, J.-M. T., Vinson, L. T., Patriarchi, T., … Berke, J. D. (2019). Dissociable dopamine dynamics for learning and motivation. Nature, 570(7759), 65–70. doi: 10.1038/s41586-019-1235-y

63. Moutoussis, M., Bullmore, E. T., Goodyer, I. M., Fonagy, P., Jones, P. B., Dolan, R. J., & Dayan, P. (2018). Change, stability, and instability in the Pavlovian guidance of behaviour from adolescence to young adulthood. PLOS Computational Biology, 14(12), e1006679. doi: 10.1371/journal.pcbi.1006679

64. Murphy, P. R., O’Connell, R. G., O’Sullivan, M., Robertson, I. H., & Balsters, J. H. (2014). Pupil diameter covaries with BOLD activity in human locus coeruleus. Human Brain Mapping, 35(8), 4140–4154. doi: 10.1002/hbm.22466

65. Murphy, P. R., Robertson, I. H., Balsters, J. H., & O’Connell, R. G. (2011). Pupillometry and P3 index the locus coeruleus-noradrenergic arousal function in humans. Psychophysiology, 48(11), 1532–1543. doi: 10.1111/j.1469-8986.2011.01226.x

66. Nicola, S. M., Woodward Hopf, F., & Hjelmstad, G. O. (2004). Contrast enhancement: A physiological effect of striatal dopamine? Cell and Tissue Research, 318(1), 93–106. doi: 10.1007/s00441-004-0929-z

67. Nieuwenhuis, S., Aston-Jones, G., & Cohen, J. D. (2005). Decision making, the P3, and the locus coeruleus—Norepinephrine system. Psychological Bulletin, 131(4), 510–532. doi: 10.1037/0033-2909.131.4.510

68. Nord, C. L., Lawson, R. P., Huys, Q. J. M., Pilling, S., & Roiser, J. P. (2018). Depression is associated with enhanced aversive Pavlovian control over instrumental behaviour. Scientific Reports, 8(1), 12582. doi: 10.1038/s41598-018-30828-5

69. O’Doherty, J. P., Cockburn, J., & Pauli, W. M. (2017). Learning, reward, and decision making. Annual Review of Psychology, 68(1), 73–100. doi: 10.1146/annurev-psych-010416-044216

70. Ousdal, O. T., Huys, Q. J., Milde, A. M., Craven, A. R., Ersland, L., Endestad, T., … Dolan, R. J. (2018). The impact of traumatic stress on Pavlovian biases. Psychological Medicine, 48(02), 327–336. doi: 10.1017/S003329171700174X

71. Park, J., Coddington, L. T., & Dudman, J. T. (2020). Basal ganglia circuits for action specification. Annual Review of Neuroscience, 43(1), annurev-neuro-070918-050452. doi: 10.1146/annurev-neuro-070918-050452

72. Queirazza, F., Steele, J. D., Krishnadas, R., Cavanagh, J., & Philiastides, M. G. (2023). Functional magnetic resonance imaging signatures of Pavlovian and instrumental valuation systems during a modified orthogonalized Go/No-Go task. Journal of Cognitive Neuroscience, 35(12), 2089–2109. doi: 10.1162/jocn_a_02062

73. R Core Team. (2022). R: A language and environment for statistical computing. In R Foundation for Statistical Computing. Vienna, Austria: R Foundation for Statistical Computing. Retrieved from www.R-project.org

74. Richer, F., & Beatty, J. (1985). Pupillary dilations in movement preparation and execution. Psychophysiology, 22(2), 204–207. doi: 10.1111/j.1469-8986.1985.tb01587.x

75. Richer, F., Silverman, C., & Beatty, J. (1983). Response selection and initiation in speeded reactions: A pupillometric analysis. Journal of Experimental Psychology: Human Perception and Performance, 9(3), 360–370. doi: 10.1037/0096-1523.9.3.360

76. Roelofs, K. (2017). Freeze for action: Neurobiological mechanisms in animal and human freezing. Philosophical Transactions of the Royal Society B: Biological Sciences, 372(1718), 20160206. doi: 10.1098/rstb.2016.0206

77. Roelofs, K., & Dayan, P. (2022). Freezing revisited: Coordinated autonomic and central optimization of threat coping. Nature Reviews Neuroscience, 23(9), 568–580. doi: 10.1038/s41583-022-00608-2

78. Rondeel, E., Van Steenbergen, H., Holland, R., & van Knippenberg, A. (2015). A closer look at cognitive control: Differences in resource allocation during updating, inhibition and switching as revealed by pupillometry. Frontiers in Human Neuroscience, 9. Retrieved from https://www.frontiersin.org/articles/10.3389/fnhum.2015.00494

79. Rösler, L., & Gamer, M. (2019). Freezing of gaze during action preparation under threat imminence. Scientific Reports, 9(1), 17215. doi: 10.1038/s41598-019-53683-4

80. Schacht, A., Dimigen, O., & Sommer, W. (2010). Emotions in cognitive conflicts are not aversive but are task specific. Cognitive, Affective, & Behavioral Neuroscience, 10(3), 349–356. doi: 10.3758/CABN.10.3.349

81. Schad, D. J., Rapp, M. A., Garbusow, M., Nebe, S., Sebold, M., Obst, E., … Huys, Q. J. M. (2020). Dissociating neural learning signals in human sign- and goal-trackers. Nature Human Behaviour, 4(2), 201–214. doi: 10.1038/s41562-019-0765-5

82. Schmidt, R., & Berke, J. D. (2017). A pause-then-cancel model of stopping: Evidence from basal ganglia neurophysiology. Philosophical Transactions of the Royal Society B: Biological Sciences, 372(1718). doi: 10.1098/rstb.2016.0202

83. Shadmehr, R., Reppert, T. R., Summerside, E. M., Yoon, T., & Ahmed, A. A. (2019). Movement vigor as a reflection of subjective economic utility. Trends in Neurosciences, 42(5), 323–336. doi: 10.1016/j.tins.2019.02.003

84. Shenhav, A., Musslick, S., Lieder, F., Kool, W., Griffiths, T. L., Cohen, J. D., & Botvinick, M. M. (2017). Toward a rational and mechanistic account of mental effort. Annual Review of Neuroscience, 40(1), 99–124. doi: 10.1146/annurev-neuro-072116-031526

85. Shiffrin, R. M., & Schneider, W. (1977). Controlled and automatic human information processing: II. Perceptual learning, automatic attending and a general theory. Psychological Review, 84(2), 127–190. doi: 10.1037/0033-295X.84.2.127

86. Singmann, H., Bolker, B., Westfall, J., & Aust, F. (2018). afex: Analysis of factorial experiments. Retrieved from https://cran.r-project.org/package=afex

87. Spielberger, C., Gorssuch, R., Lushene, P., Vagg, P., & Jacobs, G. (1983). Manual for the State-Trait Anxiety Inventory. Moutain View, CA: Consulting Psychologists Press.

88. Strauch, C., Wang, C., Einhäuser, W., Van der Stigchel, S., & Naber, M. (2022). Pupillometry as an integrated readout of distinct attentional networks. Trends in Neurosciences, 1–13. doi: 10.1016/j.tins.2022.05.003

89. Swart, J. C., Frank, M. J., Määttä, J. I., Jensen, O., Cools, R., & den Ouden, H. E. M. (2018). Frontal network dynamics reflect neurocomputational mechanisms for reducing maladaptive biases in motivated action. PLOS Biology, 16(10), e2005979. doi: 10.1371/journal.pbio.2005979

90. Swart, J. C., Froböse, M. I., Cook, J. L., Geurts, D. E., Frank, M. J., Cools, R., & den Ouden, H. E. (2017). Catecholaminergic challenge uncovers distinct Pavlovian and instrumental mechanisms of motivated (in)action. eLife, 6, e22169. doi: 10.7554/eLife.22169

91. Syed, E. C. J., Grima, L. L., Magill, P. J., Bogacz, R., Brown, P., & Walton, M. E. (2016). Action initiation shapes mesolimbic dopamine encoding of future rewards. Nature Neuroscience, 19(1), 34–36. doi: 10.1038/nn.4187

92. Tachibana, Y., & Hikosaka, O. (2012). The primate ventral pallidum encodes expected reward value and regulates motor action. Neuron, 76(4), 826–837. doi: 10.1016/j.neuron.2012.09.030

93. Treadway, M. T., Buckholtz, J. W., Schwartzman, A. N., Lambert, W. E., & Zald, D. H. (2009). Worth the ‘EEfRT’? The effort expenditure for rewards task as an objective measure of motivation and anhedonia. PLoS ONE, 4(8), e6598. doi: 10.1371/journal.pone.0006598

94. Turner, R. S., & Desmurget, M. (2010). Basal ganglia contributions to motor control: A vigorous tutor. Current Opinion in Neurobiology, 20(6), 704–716. doi: 10.1016/j.conb.2010.08.022

95. Urai, A. E., Braun, A., & Donner, T. H. (2017). Pupil-linked arousal is driven by decision uncertainty and alters serial choice bias. Nature Communications, 8, 14637. doi: 10.1038/ncomms14637

96. Van der Molen, M. W., Boomsma, D. I., Jennings, J. R., & Nieuwboer, R. T. (1989). Does the heart know what the eye sees? A cardiac/ pupillometric analysis of motor preparation and response execution. Psychophysiology, 26(1), 70–80. doi: 10.1111/j.1469-8986.1989.tb03134.x

97. van der Wel, P., & van Steenbergen, H. (2018). Pupil dilation as an index of effort in cognitive control tasks: A review. Psychonomic Bulletin & Review, 25(6), 2005–2015. doi: 10.3758/s13423-018-1432-y

98. van Rij, J., Hendriks, P., van Rijn, H., Baayen, R. H., & Wood, S. N. (2019). Analyzing the time course of pupillometric data. Trends in Hearing, 23, 2331216519832483. doi: 10.1177/2331216519832483

99. van Steenbergen, H., & Band, G. P. H. (2013). Pupil dilation in the Simon task as a marker of conflict processing. Frontiers in Human Neuroscience, 7(May), 215. doi: Artn 215\nDoi 10.3389/Fnhum.2013.00215

100. Varazzani, C., San-Galli, A., Gilardeau, S., & Bouret, S. (2015). Noradrenaline and dopamine neurons in the reward/effort trade-off: A direct electrophysiological comparison in behaving monkeys. Journal of Neuroscience, 35(20), 7866–7877. doi: 10.1523/JNEUROSCI.0454-15.2015

101. Walton, M. E., & Bouret, S. (2018). What is the relationship between dopamine and effort? Trends in Neurosciences, 42(2), 1–13. doi: 10.1016/j.tins.2018.10.001

102. Wessel, J. R. (2018). Surprise: A more realistic framework for studying action stopping? Trends in Cognitive Sciences, 22(9), 741–744. doi: 10.1016/j.tics.2018.06.005

103. Wessel, J. R., & Aron, A. R. (2017). On the globality of motor suppression: Unexpected events and their influence on behavior and cognition. Neuron, 93(2), 259–280. doi: 10.1016/j.neuron.2016.12.013

104. Westbrook, A., Frank, M. J., & Cools, R. (2021). A mosaic of cost–benefit control over cortico-striatal circuitry. Trends in Cognitive Sciences, 25(8), 710–721. doi: 10.1016/j.tics.2021.04.007

105. Wickham, H. (2016). ggplot2: Elegant graphics for data analysis. New York, NY: Springer-Verlag. Retrieved from https://ggplot2.tidyverse.org

106. Willenbockel, V., Sadr, J., Fiset, D., Horne, G. O., Gosselin, F., & Tanaka, J. W. (2010). Controlling low-level image properties: The SHINE toolbox. Behavior Research Methods, 42(3), 671– 684. doi: 10.3758/BRM.42.3.671

107. Zénon, A., Sidibé, M., & Olivier, E. (2014). Pupil size variations correlate with physical effort perception. Frontiers in Behavioral Neuroscience, 8(AUG), 1–8. doi: 10.3389/fnbeh.2014.00286

## Supplementary References

109. Allen, M., Frank, D., Schwarzkopf, D. S., Fardo, F., Winston, J. S., Hauser, T. U., & Rees, G. (2016). Unexpected arousal modulates the influence of sensory noise on confidence. eLife, 5, 1–17. doi: 10.7554/eLife.18103

110. Aston-Jones, G., & Cohen, J. D. (2005). An integrative theory of locus coeruleus-norepinephrine function: Adaptive gain and optimal performance. Annual Review of Neuroscience, 28, 403–450. doi: 10.1146/annurev.neuro.28.061604.135709

111. Cyders, M. A., Littlefield, A. K., Coffey, S., & Karyadi, K. A. (2014). Examination of a short English version of the UPPS-P Impulsive Behavior Scale. Addictive Behaviors, 39(9), 1372–1376. doi: 10.1016/j.addbeh.2014.02.013

112. Eldar, E., Cohen, J. D., & Niv, Y. (2013). The effects of neural gain on attention and learning. Nature Neuroscience, 16(8), 1146–1153. doi: 10.1038/nn.3428

113. Gilzenrat, M. S., Nieuwenhuis, S., Jepma, M., & Cohen, J. D. (2010). Pupil diameter tracks changes in control state predicted by the adaptive gain theory of locus coeruleus function. *Cognitive*, Affective & Behavioral Neuroscience, 10(2), 252–269. doi: 10.3758/CABN.10.2.252

114. Mudrik, L., & Deouell, L. Y. (2022). Neuroscientific evidence for processing without awareness. Annual Review of Neuroscience, 45(1), 403–423. doi: 10.1146/annurev-neuro-110920-033151

115. Muller, T. H., Mars, R. B., Behrens, T. E., & O’Reilly, J. X. (2019). Control of entropy in neural models of environmental state. eLife, 8, 1–30. doi: 10.7554/eLife.39404

116. Skora, L. I., Livermore, J. J. A., Dienes, Z., Seth, A. K., & Scott, R. B. (2023). Feasibility of unconscious instrumental conditioning: A registered replication. Cortex, 159, 101–117. doi: 10.1016/j.cortex.2022.12.003

117. Spielberger, C., Gorssuch, R., Lushene, P., Vagg, P., & Jacobs, G. (1983). Manual for the State-Trait Anxiety Inventory. Moutain View, CA: Consulting Psychologists Press.

118. Vadillo, M. A., Malejka, S., Lee, D. Y. H., Dienes, Z., & Shanks, D. R. (2022). Raising awareness about measurement error in research on unconscious mental processes. Psychonomic Bulletin & Review, 29(1), 21–43. doi: 10.3758/s13423-021-01923-y

